## Supplemental Figures for "*Wolbachia* endosymbionts manipulate GSC self-renewal and differentiation to reinforce host fertility"

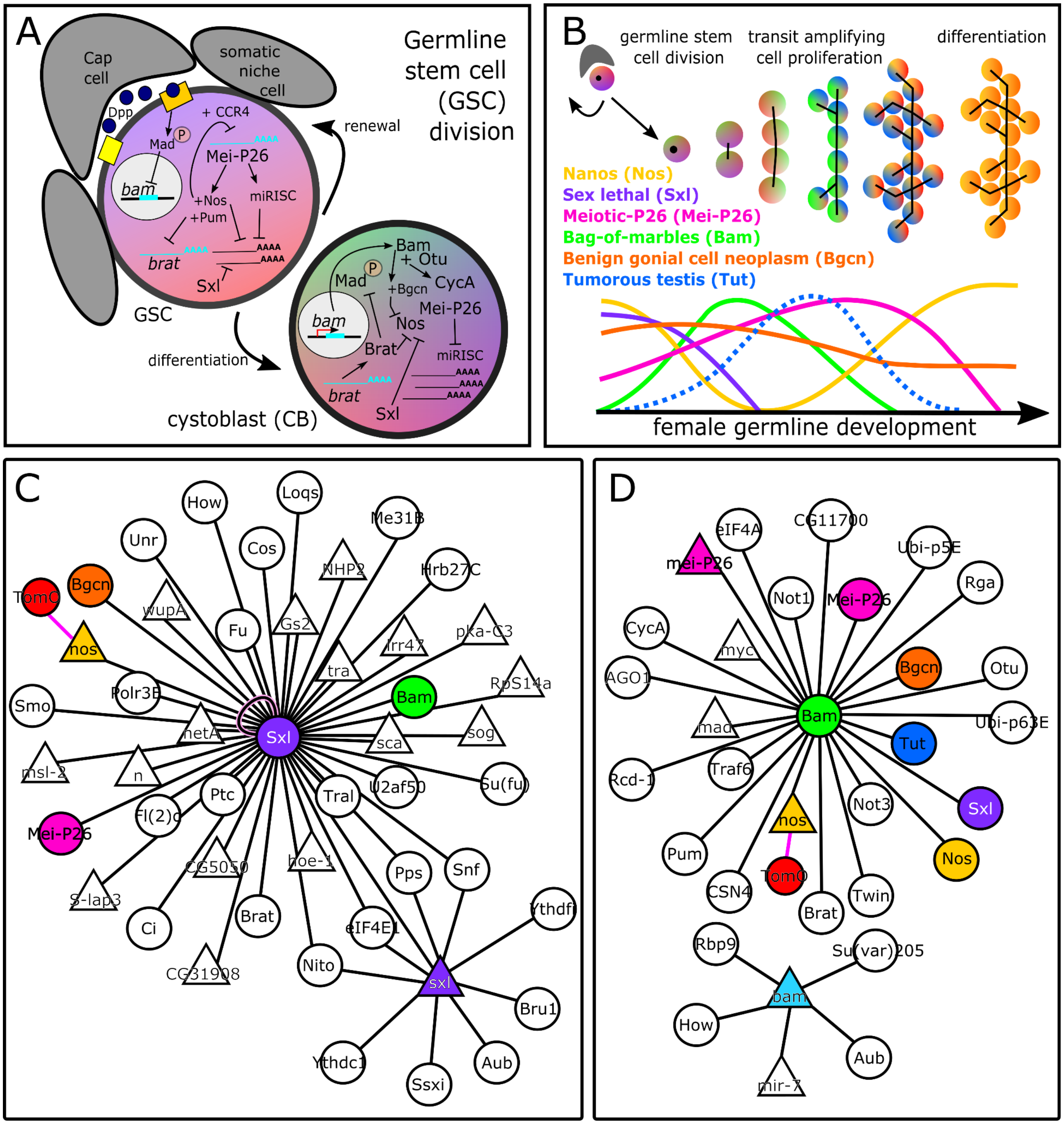


**fig S1**. Key genetic regulators of germline stem cell (GSC) maintenance and differentiation in *D. melanogaster* oogenesis. A) Diagram of the GSC niche illustrating a subset of the genes that shift in expression in the cystoblast following GSC mitosis. B) Model of the relative levels of protein expression during germline cyst development. C,D) esyN interaction networks for C) *sxl* and D) *bam* gene products (supporting references in table S1).

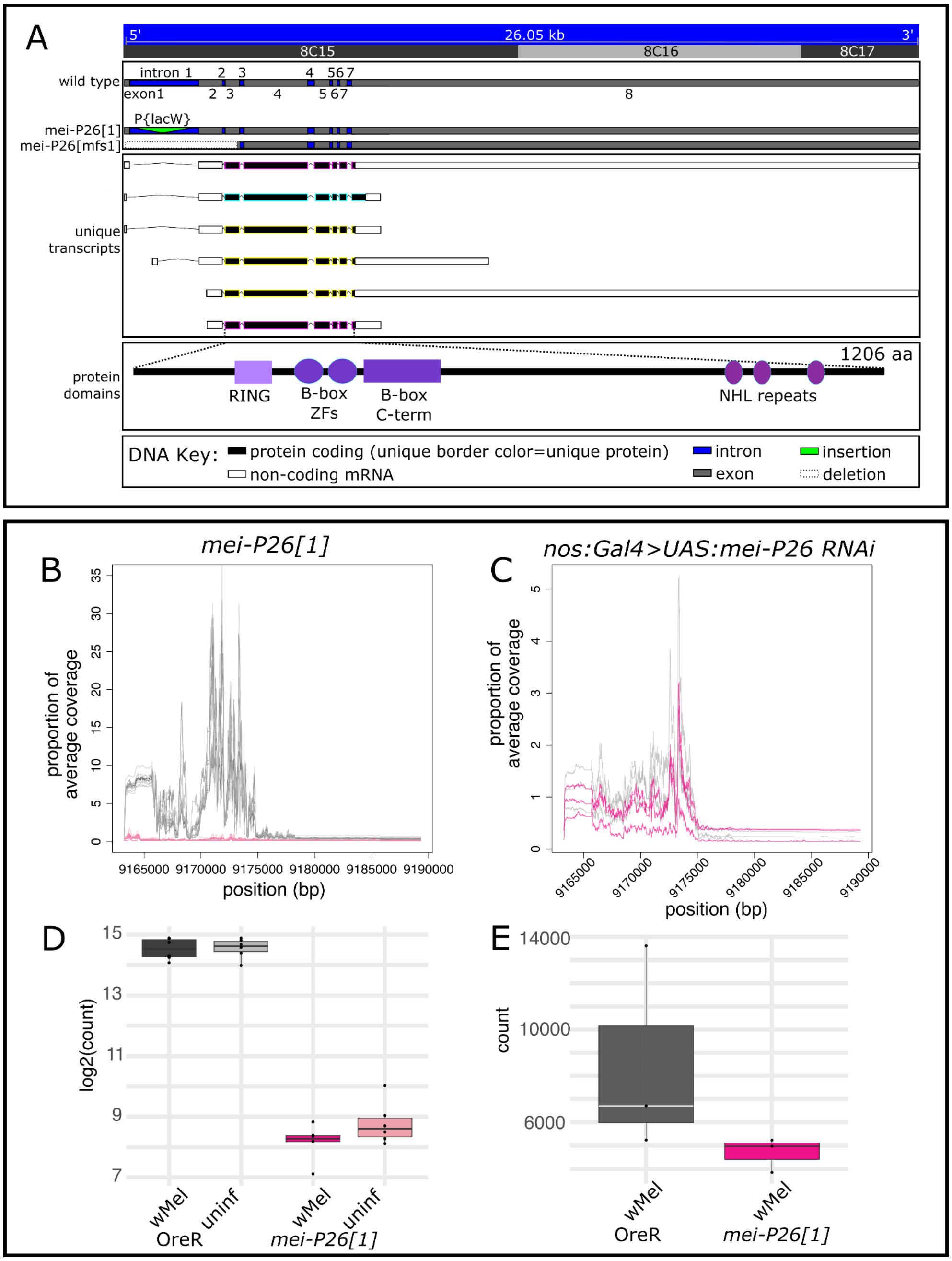


**fig S2.** *Drosophila mei-P26* genetic resources and gene expression characterization. A) Genomic map and gene model for *mei-P26* and the studied alleles. The insertion of a P{lacW} transposon in the first intron of *mei-P26[1]* impacts the RING domain. The *mei-P26[mfs1]* allele was generated by deletion of this insertion and 0.7-1.6kb of DNA flanking each side of the insertion site. B,C) mei-P26 transcript coverage and D,E) Kallisto Kallisto normalized transcript counts for *D. melanogaster mei-P26* transcripts from B,D) *mei-P26[1]* and OreR wMel-infected vs uninfected ovaries and C,E) nos:Gal4>UAS:meiP26RNAi vs OreR wMel-infected ovaries.

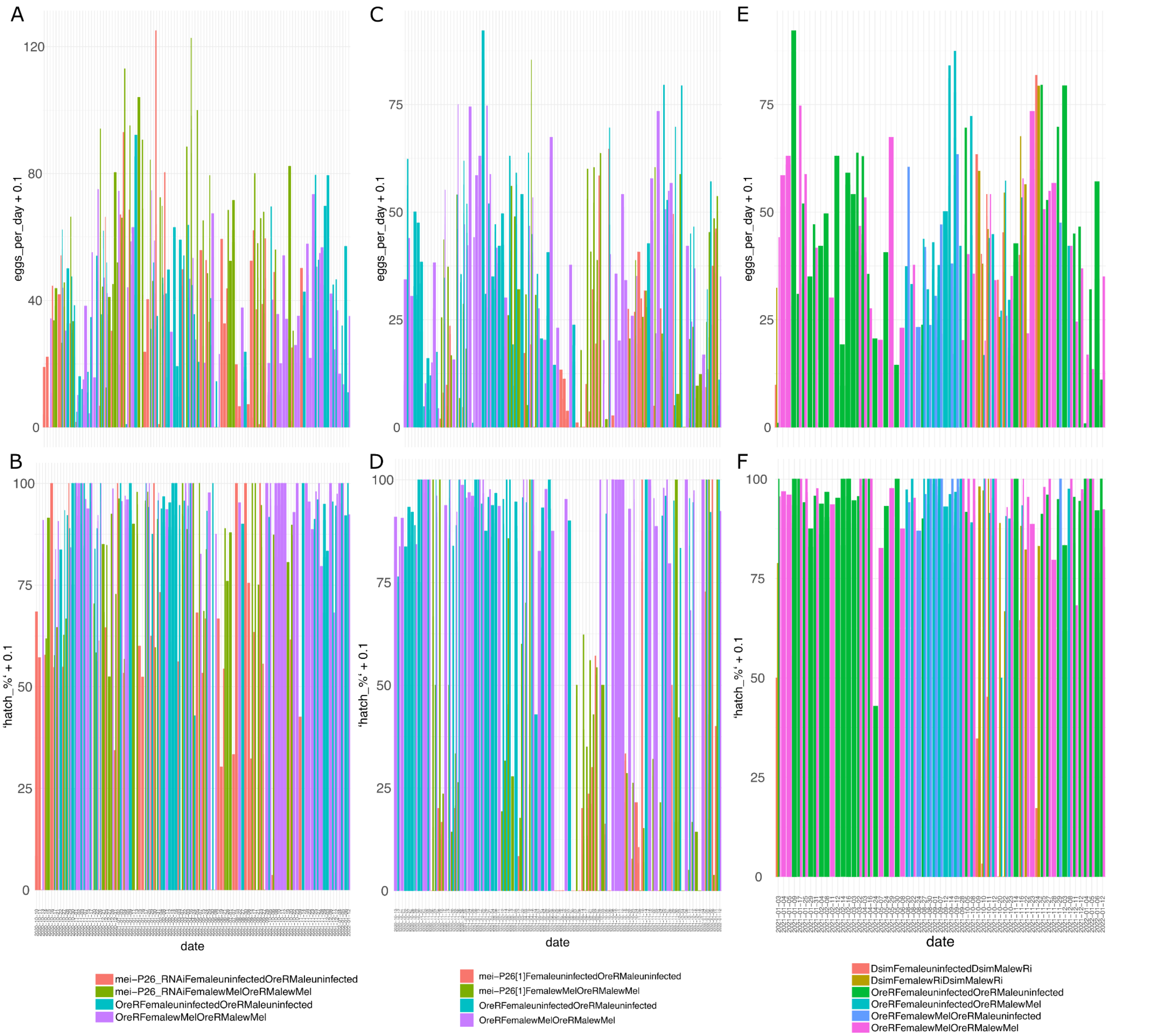


**fig S3.** Fecundity data acquisition plots vs time. A,B) Female *mei-P26* RNAi, C,D) Female *mei-P26[1]*, and E,F) CI assays. Both A,C,E) egg lay rates and B,D,F) hatch rates were consistent over time, across genetic crosses, and across fecundity crosses. A factor of 0.1 was added to the y-axis values as an offset to see zero-lay and zero-percent hatch data points.

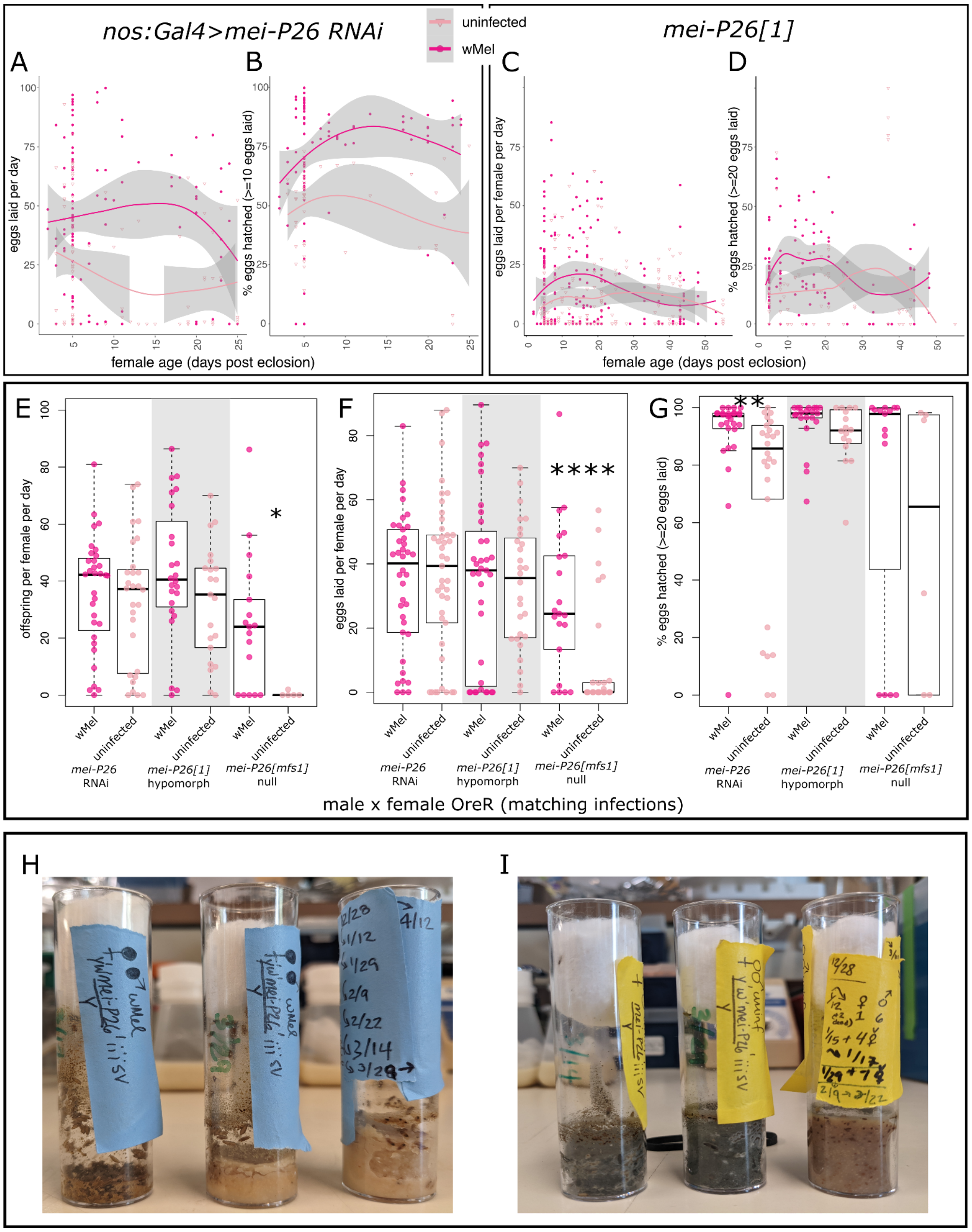


**fig S4.** Infection with wMel rescues *mei-P26* function in females and males. A-B) Hypomorphic *mei-P26[1]* and C,D) *nos:Gal4>mei-P26RNAi D. melanogaster* female fecundity vs. age, fit with a local polynomial regression (dark gray bounds=95% confidence intervals). Infection with wMel elevates offspring production across the female lifespan through increasing the number of eggs laid and the proportion of those eggs that hatch. E-G) Male *mei-P26* rescue: wMel infection produced significantly higher rates of D) overall offspring production, broken into F) egg lay and F) egg hatch, in RNAi, hypomorphic, and null *mei-P26[1]* knockdown male flies mated to wild-type females of the same age and infection status. Wilcoxon rank sum * = p<0.05, ** = 0.01, **** = 1e-4. H,I) Homozygous hypomorphic *mei-P26[1]* stocks H) infected with wMel *Wolbachia* or I) uninfected. Mold growth is uninhibited in the uninfected stocks due to embryo and larval death, which both feeds and fails to stop mold. Infection enables stable robust stock persistence and adequate larval production, which inhibit mold growth. Both stocks were started at the same time (see 12/28 on the label). The wMel-infected stock never needed any adults added, whereas the uninfected stock produced too few offspring and had to be supplemented at every vial flip to keep the stock going artificially. We ended this after a few months and the uninfected stock fully died out.

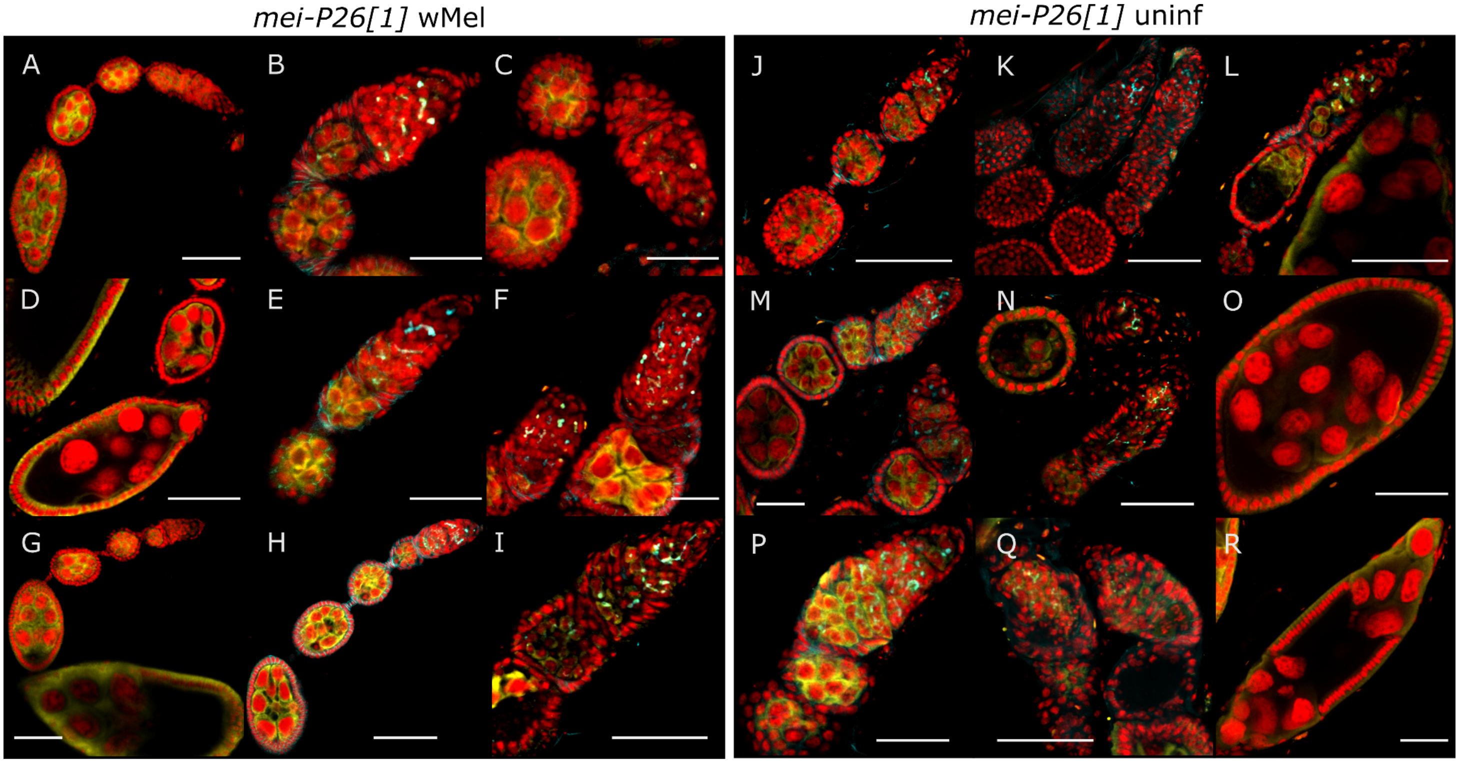


**fig S5.** Hypomorphic *mei-P26[1]* ovarioles and germaria exhibit a range of A-I) wMel-infected and J-R) uninfected phenotypes. Red = PI DNA staining, yellow = anti-Vas staining, and cyan = anti-Hts staining. Scale bars A,D,G,H,I,J-L,N,O,Q,R = 50 μm; B,C,E,F,M,P = 25 μm.

A

| genotype | infection | experiments | total mothers | total offspring | F:M sex ratio | Normal females | Normal males | NDJ females | NDJ males |
|---|---|---|---|---|---|---|---|---|---|
| mei-P26[1] | uninfected | 8 | 48 | 485 | 1.14 | 246 | 224 | 12 | 3 |
| mei-P26[1] | wMel | 7 | 24 | 273 | 0.96 | 124 | 138 | 10 | 1 |

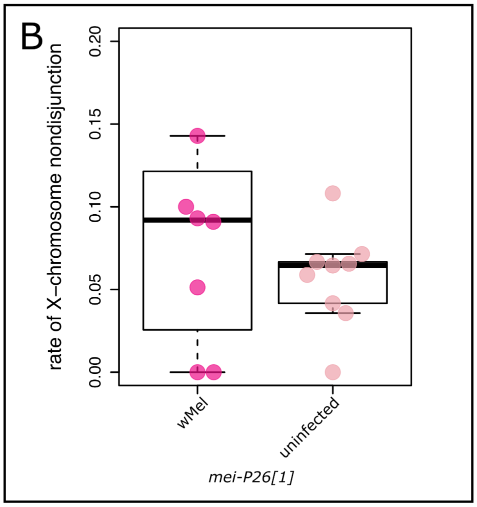


**fig S6.** Infection with wMel does not rescue *mei-P26's* function in meiosis. A) Table containing X-chromosome nondisjunction experimental data. B) Beeswarm boxplot of the rate of X-chromosome non-disjunction(NDJ) in each experiment. There was no significant difference between infected and uninfected *mei-P26[1]* females.

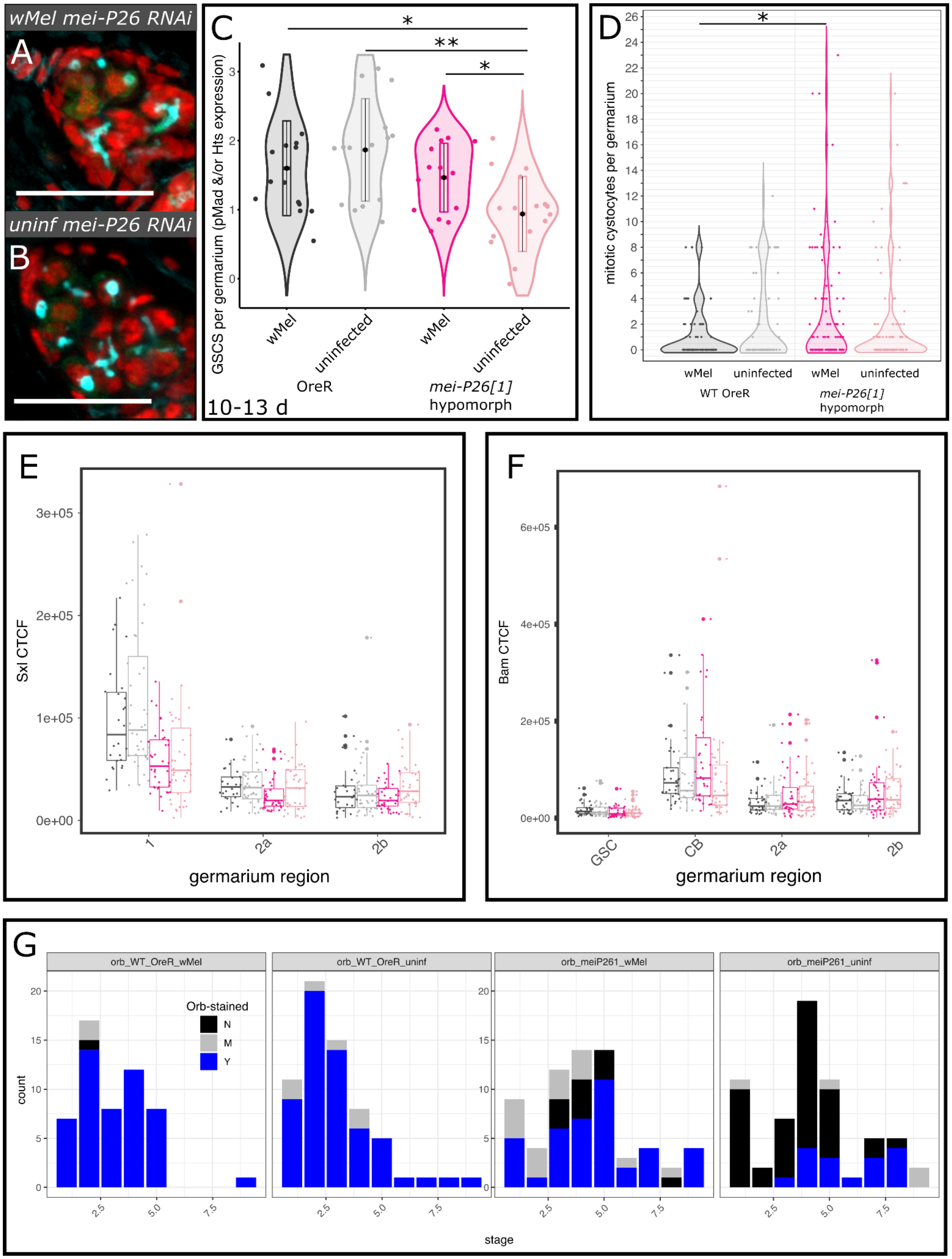


**fig S7.** GSC maintenance and germline differentiation are rescued by wMel infection. A,B) Confocal mean projections of *D. melanogaster* germaria stained with antibodies against Hts and pMad. RNAi knockdown of *mei-P26* does not affect GSC maintenance (Figure 2E). C) Violin plots of the number of GSCs per germarium in 10-13 day old females. As fully functional GSCs express pMad and have Hts-labeled spectrosomes, each was weighted by half and allows for partial scores. Wilcoxon rank sum * = p<0.05, ** = 0.01. D) F-K) Violin plots of the number of mitotic cystocytes per germarium detected by pH3 expression. E,F) Bar-scatter plots of total E) Sxl and F) Bam fluorescence expression levels across the germarium, by region. G) 1D barplots of oocyte-specific Orb staining among germline cysts, distributed across cyst developmental stages.

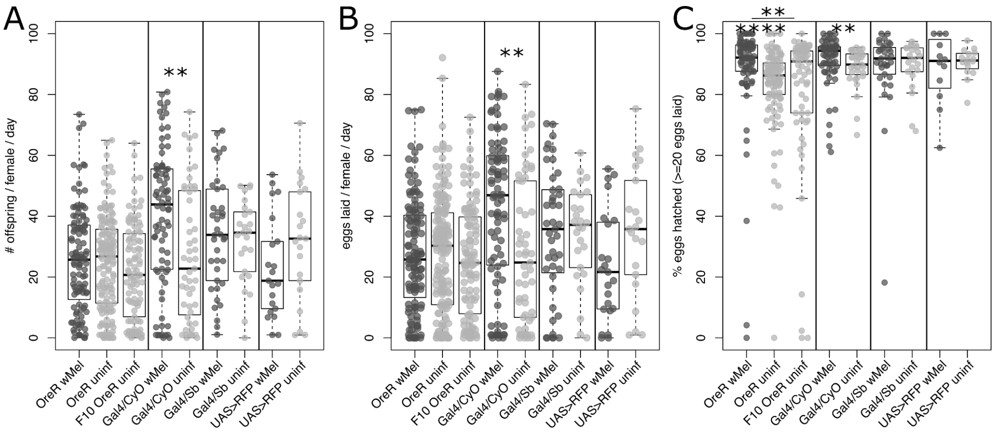


**fig S8.** The wMel strain of *Wolbachia* is a beneficial manipulator of host reproduction. A-C) Beeswarm boxplots showing that wMel infection elevates wild-type *D. melanogaster* fertility relative to uninfected flies of the same genotype. A) overall offspring production, B) egg lay, and C) egg hatch were variably impacted in different “wild-type” genotypes. D) *D. melanogaster* eggs laid per female per day plot against female age, fit with a local polynomial regression (dark gray bounds=95% confidence intervals).

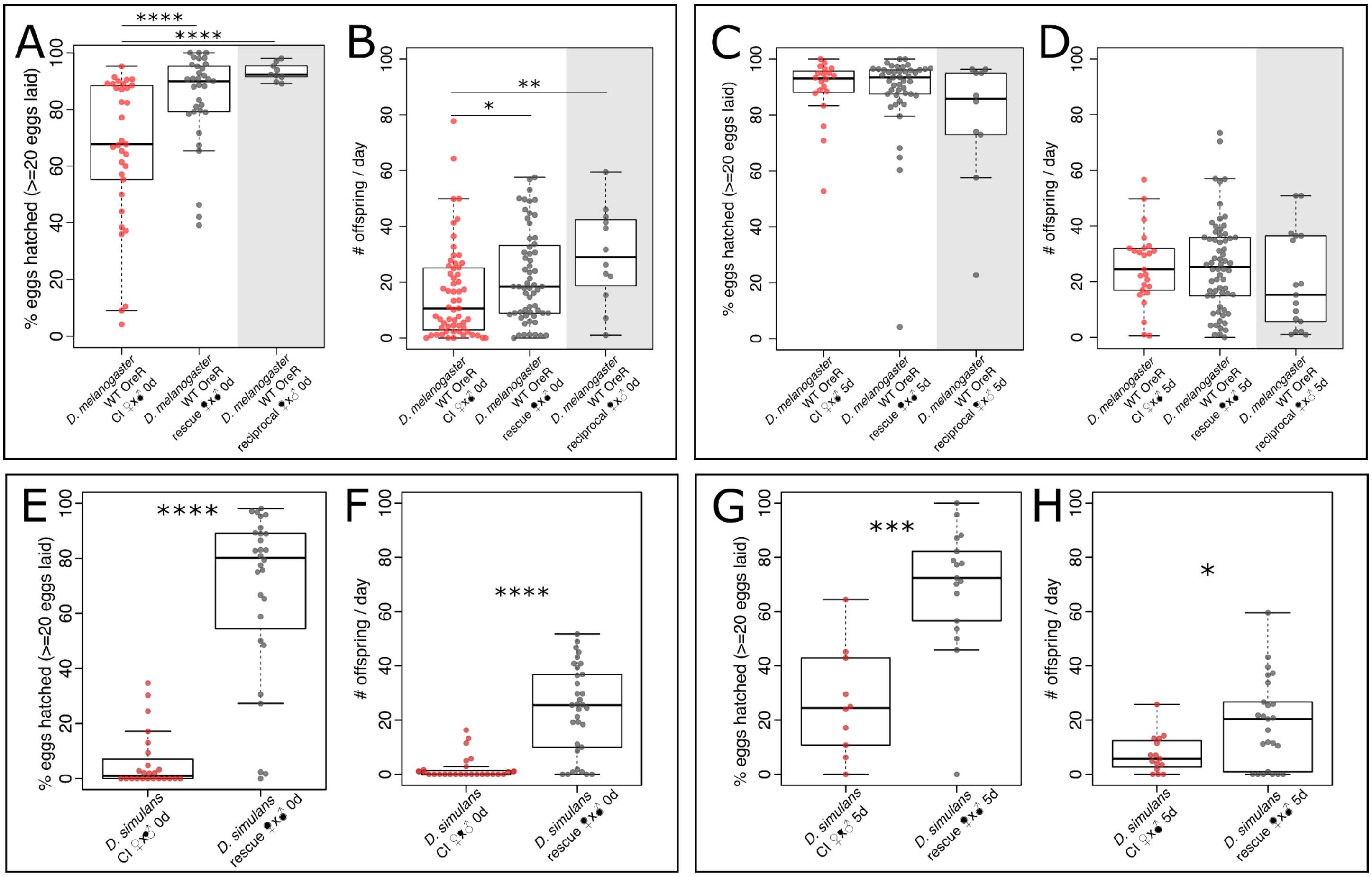


**fig S9.** Cytoplasmic incompatibility (CI) differs in strength between *Drosophila-Wolbachia* associations and weakens with male age. A-D) Beeswarm box plots of A,C) egg hatch rate and B,D) offspring production of uninfected and wMel-infected *D. melanogaster* OreR females mated to A,B) zero-day-old and C,D) 5-day-old wMel-infected males. E-H) Beeswarm box plots of E,G) egg hatch rate and F,G) offspring production in uninfected and wRi-infected *D. simulans* females mated to A,B) zero-day-old and C,D) 5-day-old wRi-infected males. Wilcoxon rank sum * = p<0.05, ** = 0.01, *** = 0.001, **** = 1e-4.

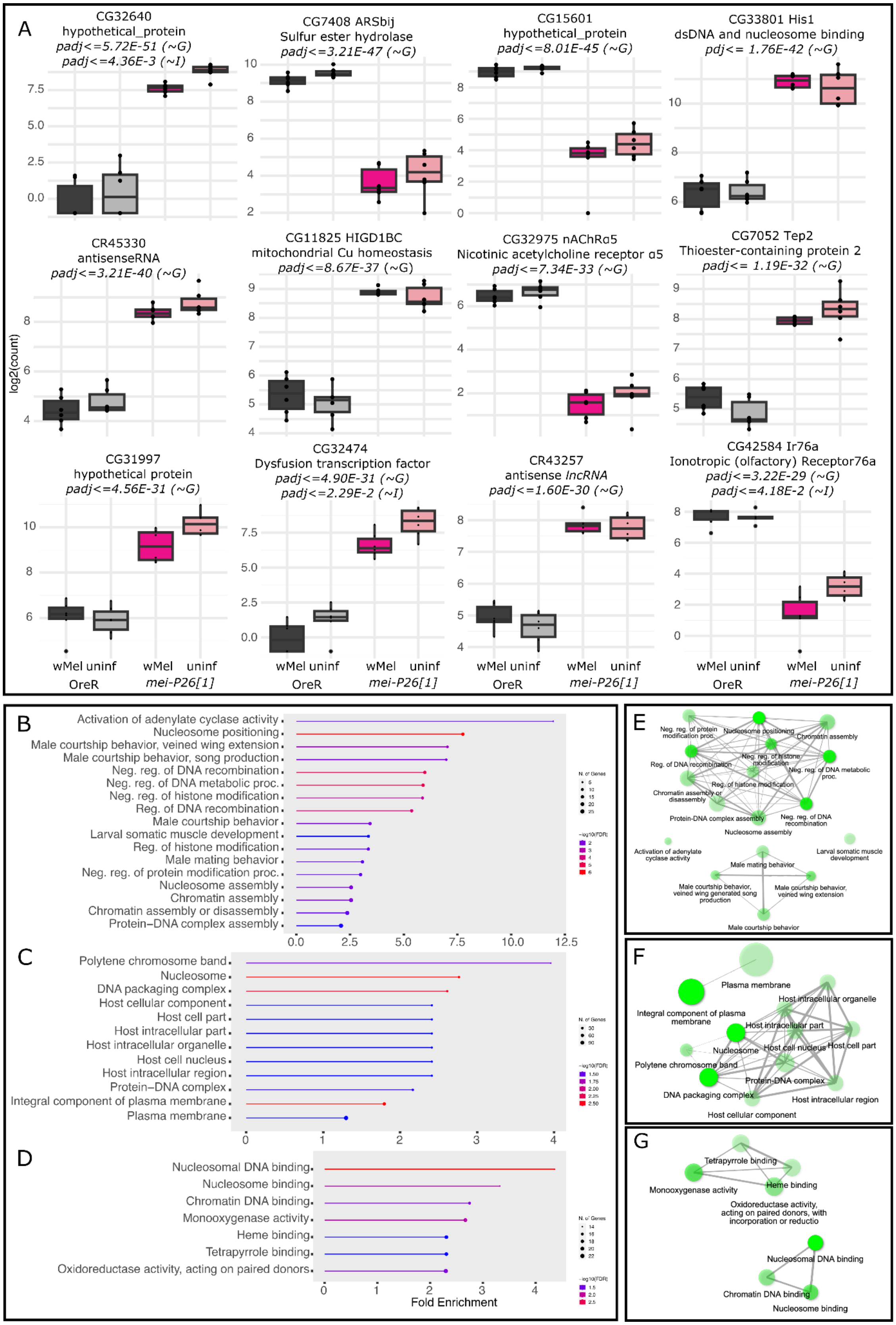


**fig S10.** *D. melanogaster* genes significantly differentially expressed due to genotype (Wald Test ~*G* vs ~*G*+*I*+*G***I*) reveal that *mei-P26* is required for the regulation of many genes. A) Kallisto normalized transcript counts for *D. melanogaster* genes (top 15 hits (including Fig 7E) *padj<=2.0E-30*; see Fig 7 and fig S2 for other plots). Barplots are colored by group: dark gray = wMel-infected OreR, light gray = uninfected OreR, dark pink

= wMel-infected *mei-P26[1]*, light pink = uninfected *mei-P26[1].* B-G) GO analysis for mei-P26-associated DE genes reveal an enrichment for processes involving chromatin, recombination, protein-protein interactions, and muscle cell differentiation. GO enrichment B-D) category plots and E-G) term interaction networks for the categories of B,E) biological process, C,F) cellular component, and D,G) molecular function.

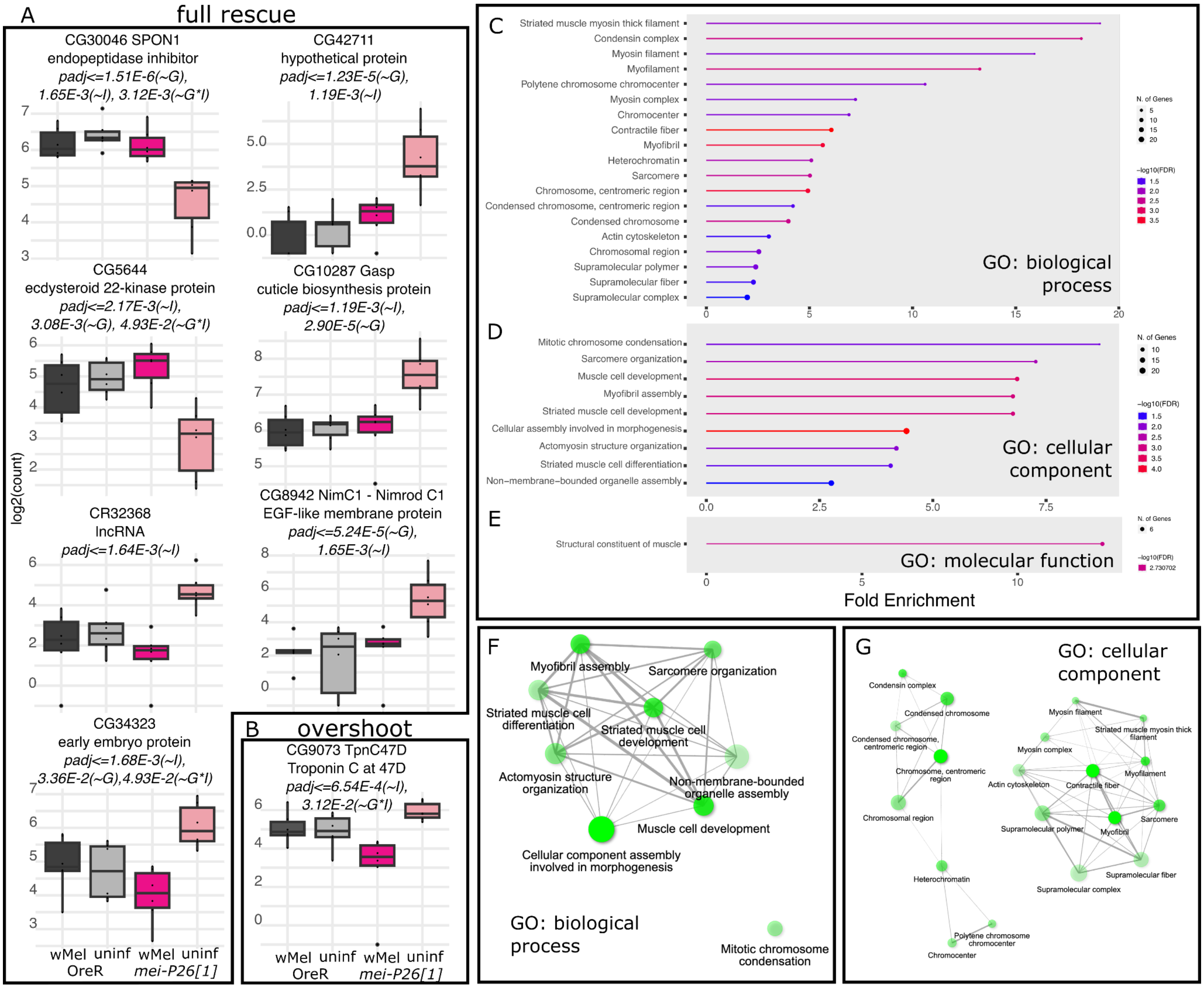


**fig S11**. *D. melanogaster* genes significantly differentially expressed due to infection state (Wald Test *~I* vs *~G+I+G*I*) reveal an enrichment of rescue events, genes for which wMel-infected *mei-P26[1]* ovaries exhibit OreR expression levels. A,B) Kallisto normalized transcript counts for *D. melanogaster* genes exhibiting A) rescue and B) overshoot of OreR expression levels (padj<=0.002; see Fig 7 for other plots). Barplots are colored by group: dark gray = wMel-infected OreR, light gray = uninfected OreR, dark pink = wMel-infected *mei-P26[1]*, light pink = uninfected *mei-P26[1].* C-G) GO analysis for infection DE genes reveal an abundance of cytoskeletal and chromatin components. GO enrichment C-E) category plots and F,G) term interaction networks for the categories of C,F) biological process, D,G) cellular component, and E) molecular function (no network for a single term).

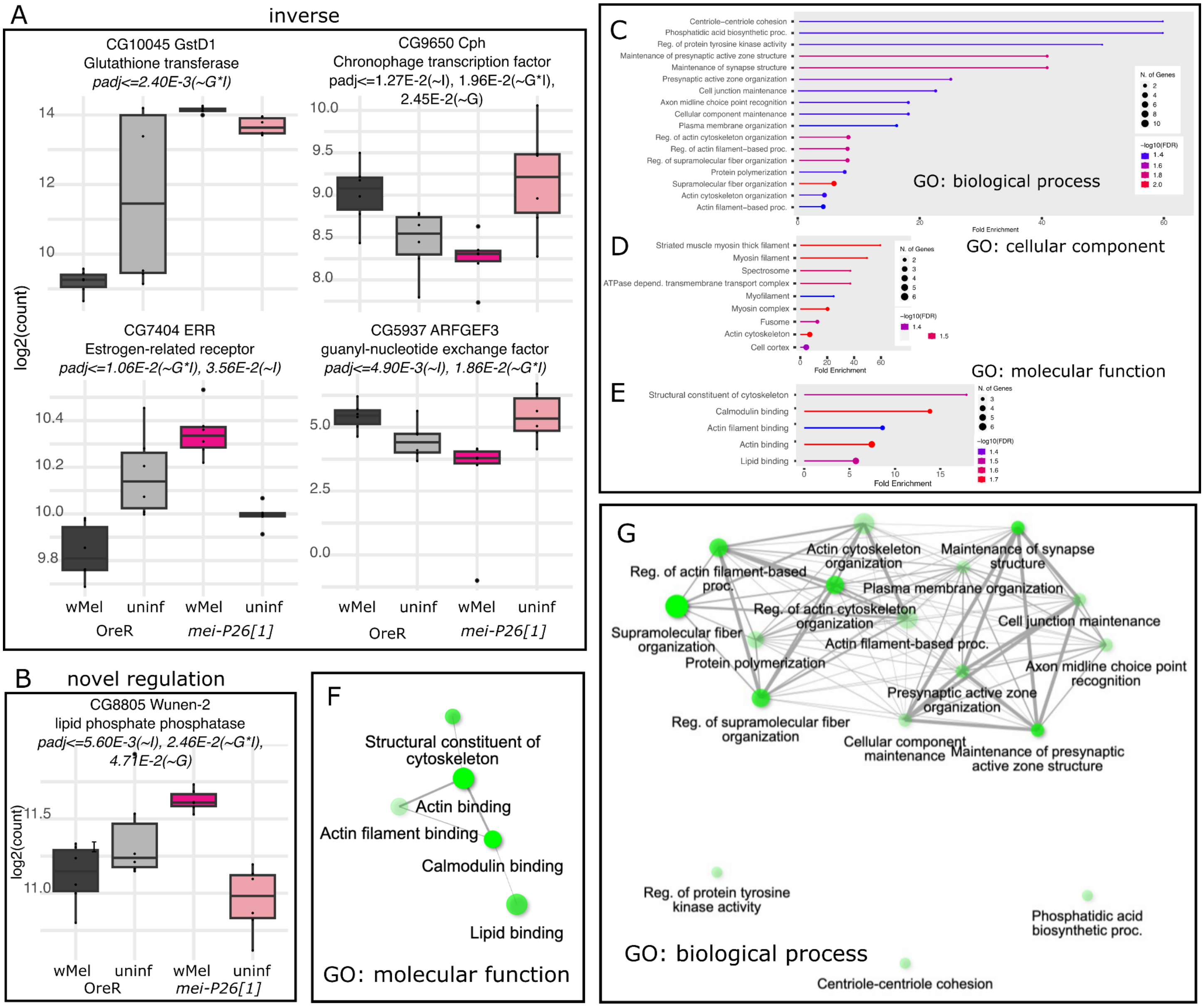


**fig S12.** *D. melanogaster* genes significantly differentially expressed due to the joint infection-by-genotype state (Wald Test ~G*I vs ~G+I+G*I) reveal an enrichment of reversal DEG, genes for which wMel-infected ovaries exhibit inverse DE patterns for *mei-P26[1]* and OreR ovaries. A-C) Kallisto normalized transcript counts for *D. melanogaster* genes exhibiting A) inverse, B) undershoot, and C) novel regulation of OreR expression levels (padj<=0.02; see Fig 7 for other plots). Barplots are colored by group: dark gray = wMel-infected OreR, light gray = uninfected OreR, dark pink = wMel-infected *mei-P26[1]*, light pink = uninfected *mei-P26[1].* D-G) GO analysis for infection DE genes reveal cytoskeletal and membrane factors. GO enrichment C-E) category plots and F,G) term interaction networks for the categories of C,G) biological process, D) cellular component (see Fig 8D for network), and E,F) molecular function.

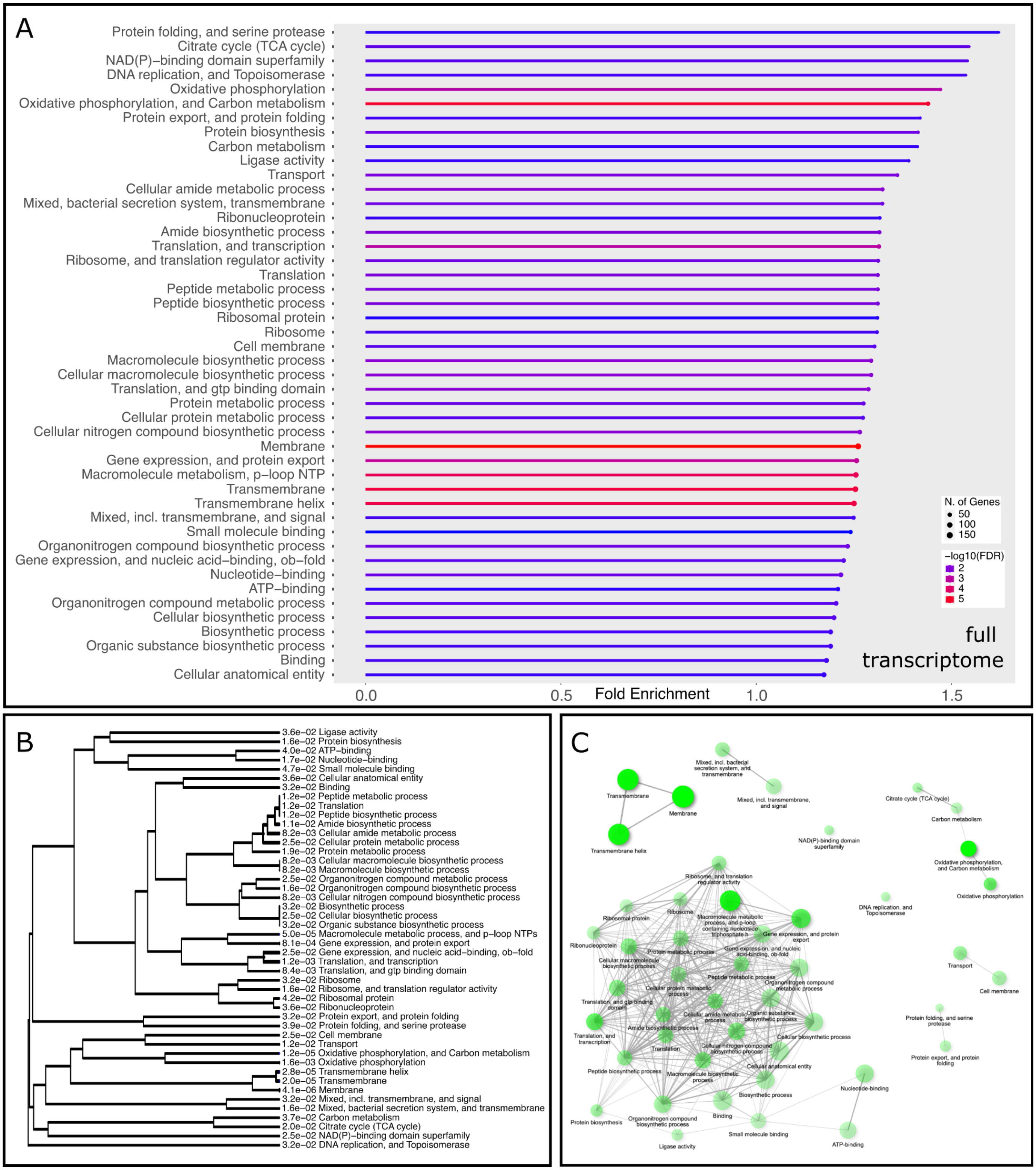


**fig S13.** wMel *Wolbachia* transcriptome differential expression GO categories of all expressed genes reveal cell maintenance, central metabolic, and membrane-associated processes. GO enrichment A) category plot, B) hierarchical clustering tree, and C) term interaction network the biological process category.

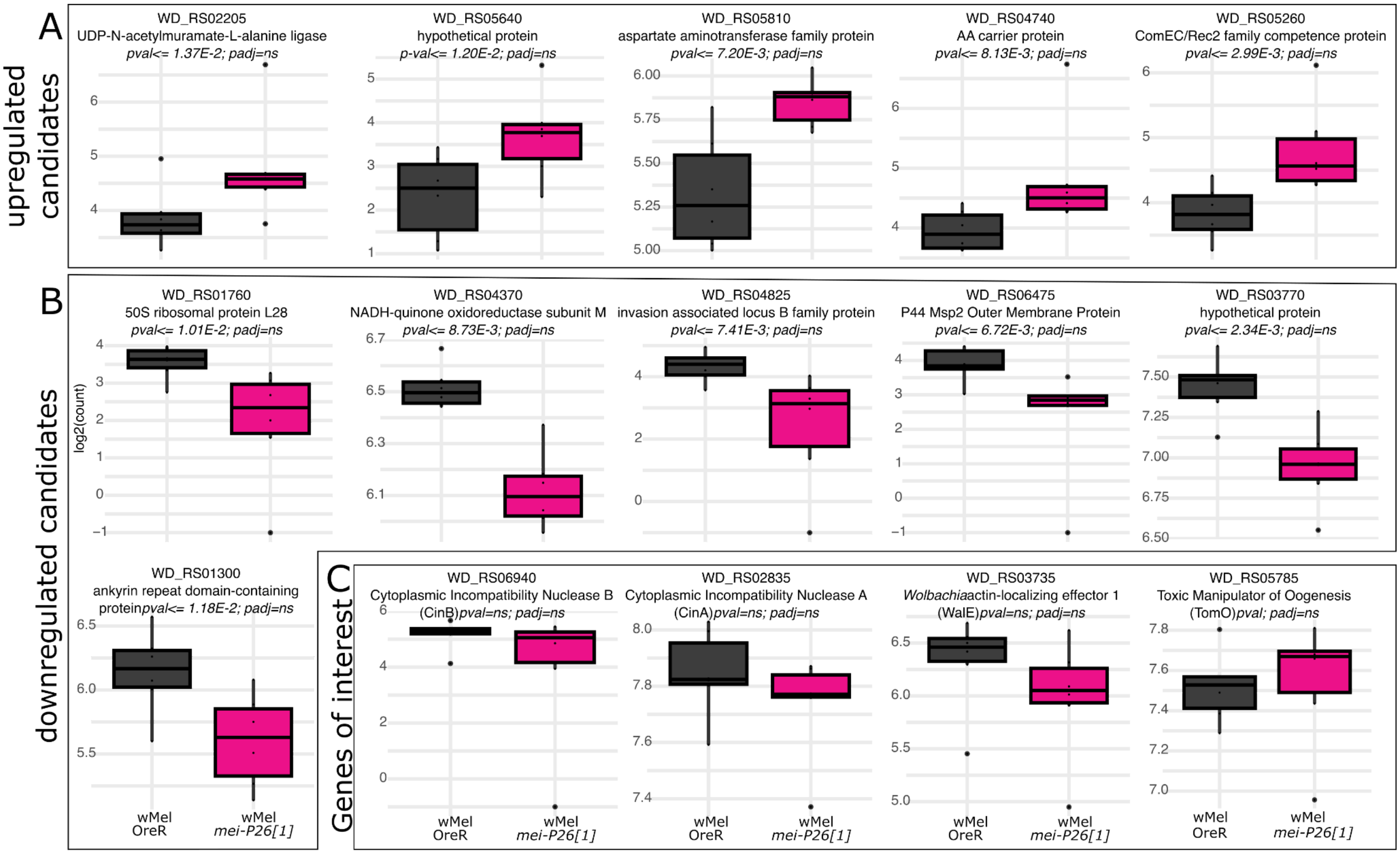


**figS14.** Kallisto normalized transcript counts for *w*Mel *Wolbachia* A,B) differential expression candidate genes (pending deeper sampling) and C) genes of interest from the literature. Barplots are colored by group: dark gray = wMel-infected OreR, light gray = uninfected OreR, dark pink = wMel-infected *mei-P26[1]*, light pink = uninfected *mei-P26[1].* Wald Test genotype-association p-values and adjusted p-values (padj).

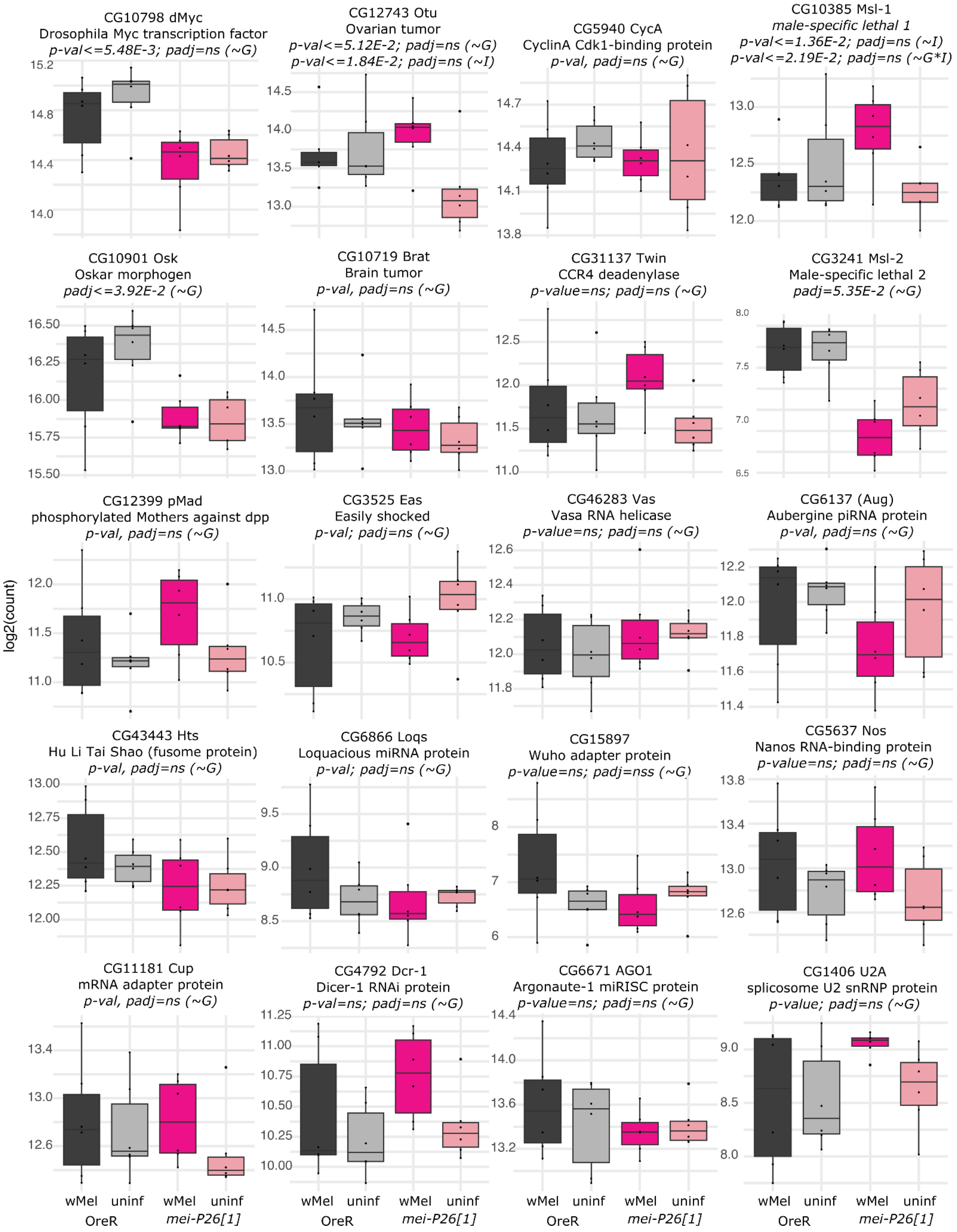


**fig S15.** Non-significant Kallisto normalized transcript counts for a subset of essential *D. melanogaster* oogenesis genes from the literature selected based upon their known interactions with *mei-P26*, *sxl*, *bam,* or germ plasm formation (a mid-stage 9 of oogenesis). Barplots are colored by group: dark gray = wMel-infected OreR, light gray = uninfected OreR, dark pink = wMel-infected *mei-P26[1]*, light pink = uninfected *mei-P26[1].*
