## Supplemental Tables for "*Wolbachia* endosymbionts manipulate GSC self-renewal and differentiation to reinforce host fertility"

| **source** | **type** | **target** | **citations** |
|---|---|---|---|
| AGO1" | physical | bam | pubmed:33907499 |
| AGO1" | physical | mei-P26 | pubmed:22438571, pubmed:18528333, pubmed:29354790 |
| bam" | physical | aub | pubmed:28441530 |
| bgcn" | physical | bam | pubmed:19556547, pubmed:25119050, pubmed:25412508, pubmed:19470484, pubmed:23526974, pubmed:20018853, pubmed:26291077, pubmed:28190776, pubmed:23122292 |
| brat" | physical | bam | pubmed:33907499, pubmed:29354790 |
| CG11700" | physical | bam | pubmed:28484036 |
| CG6304" | physical | mei-P26 | pubmed:22036573 |
| CSN4" | physical | bam | pubmed:25119050 |
| CycA" | physical | bam | pubmed:28484036 |
| eIF4A" | physical | bam | pubmed:19556547 |
| gw" | physical | mei-P26 | pubmed:22438571 |
| how" | physical | bam | pubmed:20362539 |
| how" | physical | Sxl | pubmed:23788626 |
| Mad" | physical | bam | pubmed:31097674 |
| mei-P26" | physical | aub | pubmed:28441530 |
| mei-P26" | physical | bam | pubmed:23122292 |
| mei-P26" | physical | bam | pubmed:23526974, pubmed:29354790 |
| mei-P26" | physical | bgcn | pubmed:23122292 |
| mei-P26" | physical | bgcn | pubmed:23526974, pubmed:29354790 |
| mei-P26" | physical | nos | pubmed:32654316, pubmed:23526974 |
| mei-P26" | physical | nos | pubmed:22438571 |
| mei-P26" | physical | orb | pubmed:22438571 |
| mei-P26" | physical | pum | pubmed:24286029, pubmed:30590052 |
| mei-P26" | physical | twin | pubmed:24286029 |

| | | | |
|---|---|---|---|
| mir-137" | physical | mei-P26 | pubmed:20400939 |
| mir-7" | physical | bam | pubmed:19758565 |
| mir-ban" | physical | mei-P26 | pubmed:20400939 |
| Myc" | physical | bam | pubmed:33907499 |
| nos" | physical | bam | pubmed:32654316 |
| nos" | physical | bam | pubmed:20018853 |
| Not1" | physical | bam | pubmed:29255063 |
| Not3" | physical | bam | pubmed:29255063 |
| otu" | physical | bam | pubmed:28484036, pubmed:30879902 |
| pum" | physical | bam | pubmed:20018853 |
| Rbp9 | physical | bam | pubmed:10082516 |
| Rcd-1" | physical | bam | pubmed:29255063 |
| Rga" | physical | bam | pubmed:29255063 |
| Su(var)205" | physical | bam | pubmed:30867469 |
| Sxl | physical | CG31908 | pubmed:21663794 |
| Sxl" | physical | aub | pubmed:28441530 |
| Sxl" | physical | bam | pubmed:23526974 |
| Sxl" | physical | bgcn | pubmed:23526974, pubmed:29354790 |
| Sxl" | physical | brat | pubmed:29354790 |
| Sxl" | physical | bru1 | pubmed:17067567 |
| Sxl" | physical | CG5050 | pubmed:21663794 |
| Sxl" | physical | ci | pubmed:14597576, pubmed:17284519 |
| Sxl" | physical | cos | pubmed:14597576, pubmed:17284519 |
| Sxl" | physical | eIF4E1 | pubmed:21829374 |
| Sxl" | physical | fl(2)d | pubmed:18245840 |
| Sxl" | physical | fu | pubmed:14597576 |
| Sxl" | physical | Gs2 | pubmed:34029324 |
| Sxl" | physical | hoe1 | pubmed:21663794 |

| | | | |
|---|---|---|---|
| Sxl" | physical | Hrb27C | pubmed:29635389 |
| Sxl" | physical | loqs | pubmed:23788626 |
| Sxl" | physical | Lrr47 | pubmed:21663794 |
| Sxl" | physical | me31B | pubmed:23788626 |
| Sxl" | physical | mei-P26 | pubmed:23526974, pubmed:29354790 |
| Sxl" | physical | msl-2 | pubmed:9144292, pubmed:29089381, pubmed:25209665, pubmed:30562515, pubmed:18203923, pubmed:21663794, pubmed:16452509, pubmed:29635389, pubmed:23788626, pubmed:14532129 |
| Sxl" | physical | N | pubmed:17276344 |
| Sxl" | physical | NetA | pubmed:30562515 |
| Sxl" | physical | NHP2 | pubmed:29845608 |
| Sxl" | physical | nito | pubmed:26324914 |
| Sxl" | physical | nos | pubmed:29845608, pubmed:22645327, pubmed:23526974, pubmed:32654316 |
| Sxl" | physical | Pka-C3 | pubmed:30562515 |
| Sxl" | physical | Polr3E | pubmed:10521666 |
| Sxl" | physical | pps | pubmed:20221253 |
| Sxl" | physical | ptc | pubmed:17284519 |
| Sxl" | physical | RpS14a | pubmed:21663794 |
| Sxl" | physical | S-Lap3 | pubmed:21663794 |
| Sxl" | physical | sca | pubmed:30562515 |
| Sxl" | physical | smo | pubmed:17284519 |
| Sxl" | physical | snf | pubmed:19727396, pubmed:20221253 |
| Sxl" | physical | snf | pubmed:19013444, pubmed:18245840, pubmed:20221253 |
| Sxl" | physical | sog | pubmed:30562515 |
| Sxl" | physical | ssx | pubmed:30590805 |
| Sxl" | physical | Su(fu) | pubmed:14597576 |
| Sxl" | physical | Sxl | pubmed:30562515, pubmed:28675155, pubmed:21829374, pubmed:27919081, pubmed:26324914, pubmed:20221253 |
| Sxl" | physical | Sxl | pubmed:17284519 |

| | | | |
|---|---|---|---|
| Sxl" | physical | tra | pubmed:20221253, pubmed:34029324, pubmed:25453831, pubmed:30562515 |
| Sxl" | physical | tral | pubmed:23788626 |
| Sxl" | physical | U2af50 | pubmed:18245840 |
| Sxl" | physical | Unr | pubmed:23788626, pubmed:16452509, pubmed:18203923 |
| Sxl" | physical | Ythdc1 | pubmed:28675155, pubmed:27919081 |
| Sxl" | physical | Ythdf | pubmed:28675155 |
| Traf6" | physical | bam | pubmed:30879902 |
| tut | physical | mei-P26 | pubmed:25412508 |
| tut" | physical | bam | pubmed:28190776, pubmed:25412508 |
| twin" | physical | bam | pubmed:29255063, pubmed:28190776, pubmed:26549449 |
| U2A" | physical | mei-P26 | pubmed:27035939 |
| Ubi-p5E" | physical | bam | pubmed:28484036 |
| Ubi-p63E" | physical | bam | pubmed:28484036 |
| vas" | physical | mei-P26 | pubmed:19952109 |
| wuho" | physical | mei-P26 | pubmed:31941704 |
| wupA" | physical | Sxl | pubmed:34029324 |
| bam | suppressible | mei-P26 | pubmed:23122292 |
| U2A | suppressible | mei-P26 | pubmed:27035939 |
| mei-P26 | suppressible | eas | pubmed:15937125 |
| mei-P26 | suppressible | Dcr-1 | pubmed:20400939 |
| mei-P26 | suppressible | twin | pubmed:24286029 |
| mei-P26 | suppressible | jus | pubmed:15937125 |
| loqs | suppressible | mei-P26 | pubmed:18528333 |
| bam | enhanceable | mei-P26 | pubmed:10924472 |
| vas | enhanceable | mei-P26 | pubmed:19952109 |

**table S1**. esyN references for *sxl* and *bam* interactions in Supplemental fig S1C and *mei-P26* interactions in Fig 9.

| interaction | sex | reference |
|---|---|---|
| Bam is a translational repressor in female GSCs | females | Shen, R., Weng, C., Yu, J., and Xie, T. (2009). eIF4A controls germline stem cell self-renewal by directly inhibiting BAM function in the Drosophila ovary. Proceedings of the National Academy of Sciences 106, 11623–11628. |
| Bam and Bgcn bind the *nos* 3'-UTR and inhibit Nos translation in females | females | Li, Y., Minor, N.T., Park, J.K., McKearin, D.M., and Maines, J.Z. (2009). Bam and Bgcn antagonize Nanos-dependent germ-line stem cell maintenance. Proceedings of the National Academy of Sciences 106, 9304–9309. |
| Dpp GSC niche signalling silences *bam* transcription in females | females | Chen, D., and McKearin, D. (2003). Dpp Signaling Silences bam Transcription Directly to Establish Asymmetric Divisions of Germline Stem Cells. Current Biology 13, 1786–1791. 10.1016/j.cub.2003.09.033. |
| BamF localizes to the fusome/spectrosome continuously and BamC localizes to the cytoplasm in CB-> 8-cell cysts | females | McKearin, D., and Ohlstein, B. (1995). A role for the Drosophila Bag-of-marbles protein in the differentiation of cystoblasts from germline stem cells. Development 121, 2937–2947. |
| Bam alters chromatin methylation to activate gene expression in the female germline | females | Mukai, M., Hira, S., Nakamura, K., Nakamura, S., Kimura, H., Sato, M., and Kobayashi, S. (2015). H3K36 Trimethylation-Mediated Epigenetic Regulation is Activated by Bam and Promotes Germ Cell Differentiation During Early Oogenesis in Drosophila. Biology Open 4, 119–124. 10.1242/bio.201410850. |
| Bam binds Otu to deubiquitinate CycA during female TA mitoses | females | Ji, S., Li, C., Hu, L., Liu, K., Mei, J., Luo, Y., Tao, Y., Xia, Z., Sun, Q., and Chen, D. (2017). Bam-dependent deubiquitinase complex can disrupt germ-line stem cell maintenance by targeting cyclin A. Proceedings of the National Academy of Sciences 114, 6316–6321. 10.1073/pnas.1619188114. |
| Bam interacts with Bgcn and the CCR4 deadenylase complex to repress GSC maintenance factors in the female germline | females | Sgromo, A., Raisch, T., Backhaus, C., Keskeny, C., Alva, V., Weichenrieder, O., and Izaurralde, E. (2018). Drosophila Bag-of-marbles directly interacts with the CAF40 subunit of the CCR4–NOT complex to elicit repression of mRNA targets. RNA 24, 381–395. 10.1261/rna.064584.117. |
| In males, Tut binds the *mei-P26* 3'-UTR, which when bound to Bam and Bgcn represses Mei-P26 translation | males | Chen, D., Wu, C., Zhao, S., Geng, Q., Gao, Y., Li, X., Zhang, Y., and Wang, Z. (2014). Three RNA Binding Proteins Form a Complex to Promote Differentiation of Germline Stem Cell Lineage in Drosophila. PLoS Genet 10, e1004797. 10.1371/journal.pgen.1004797. |
| Bam and Bgcn bind the *mei-P26* 3'-UTR and inhibit Mei-P26 translation in male spermatocytes | males | Insco, M.L., Bailey, A.S., Kim, J., Olivares, G.H., Wapinski, O.L., Tam, C.H., and Fuller, M.T. (2012). A Self-Limiting Switch Based on Translational Control Regulates the Transition from Proliferation to Differentiation in an Adult Stem Cell Lineage. Cell Stem Cell 11, 689–700. 10.1016/j.stem.2012.08.012. |
| Bam is translationally regulated by miRNA binding to its 3'-UTR in the male germline | males | Eun, S.H., Stoiber, P.M., Wright, H.J., McMurdie, K.E., Choi, C.H., Gan, Q., Lim, C., and Chen, X. (2013). MicroRNAs downregulate Bag of marbles to ensure proper terminal differentiation in the Drosophila male germline. Development 140, 23–30. 10.1242/dev.086397. |
| Bam is translationally regulated by miR-7 in the male germline | males | Pek, J.W., Lim, A.K., and Kai, T. (2009). Drosophila Maelstrom Ensures Proper Germline Stem Cell Lineage Differentiation by Repressing microRNA-7. Developmental Cell 17, |

| | | |
|---|---|---|
| | | 417–424. 10.1016/j.devcel.2009.07.017. |
| Bam transcription is silenced by a histone linker protein | males | Carbonell, A., Pérez-Montero, S., Climent-Cantó, P., Reina, O., and Azorín, F. (2017). The Germline Linker Histone dBigH1 and the Translational Regulator Bam Form a Repressor Loop Essential for Male Germ Stem Cell Differentiation. Cell Reports 21, 3178–3189. 10.1016/j.celrep.2017.11.060. |
| mei-P26 genetically interacts with Bam as a dominant enhancer | females | Page, S.L., McKim, K.S., Deneen, B., Van Hook, T.L., and Hawley, R.S. (2000). Genetic Studies of <em>mei-P26</em> Reveal a Link Between the Processes That Control Germ Cell Proliferation in Both Sexes and Those That Control Meiotic Exchange in Drosophila. Genetics 155, 1757. |
| mei-P26 binds Ago1 through its NHL domain and inhibits miRNA production; loquacious knockdown suppresses mei-P26 knockdown phenotype | females | Neumüller, R.A., Betschinger, J., Fischer, A., Bushati, N., Poernbacher, I., Mechtler, K., Cohen, S.M., and Knoblich, J.A. (2008). Mei-P26 regulates microRNAs and cell growth in the Drosophila ovarian stem cell lineage. Nature 454, 241–245. 10.1038/nature07014. |
| Vas activates *mei-P26* translation by 3'-UTR binding and interaction with eIF5B | females | Liu, N., Han, H., and Lasko, P. (2009). Vasa promotes Drosophila germline stem cell differentiation by activating mei-P26 translation by directly interacting with a (U)-rich motif in its 3' UTR. Genes & Development 23, 2742–2752. 10.1101/gad.1820709. |
| mei-P26 down-regulates eIF4E in GSCs | females | Song, Y., and Lu, B. (2011). Regulation of cell growth by Notch signaling and its differential requirement in normal vs. tumor-forming stem cells in Drosophila. Genes Dev. 25, 2644–2658. 10.1101/gad.171959.111. |
| mei-P26 inhibits Orb by Ago1-based miRNA-binding to the Orb 3'-UTR; mei-P26 also down-regulates Brat (and Bam because Brat--\|pMad) expression in the GSC | females | Li, Y., Maines, J.Z., Tastan, O.Y., McKearin, D.M., and Buszczak, M. (2012). Mei-P26 regulates the maintenance of ovarian germline stem cells by promoting BMP signaling. Development 139, 1547–1556. 10.1242/dev.077412. |
| CCR4-NOT works with Nos and Pum to deadenylate mei-P26 in the GSC; CCR4 was present in the GSCs as well as in other cells in the germarium (Figure 1C) where it was mostly cytoplasmic and accumulated in discrete foci, as reported in other cell types in the ovary and embryo (Rouget et al., 2010; Temme et al., 2004; Zaessinger et al., 2006). | females | Joly, W., Chartier, A., Rojas-Rios, P., Busseau, I., and Simonelig, M. (2013). The CCR4 Deadenylase Acts with Nanos and Pumilio in the Fine-Tuning of Mei-P26 Expression to Promote Germline Stem Cell Self-Renewal. Stem Cell Reports 1, 411–424. 10.1016/j.stemcr.2013.09.007. |
| "Mei-P26 associates with Bam, Bgcn and Sxl and nanos mRNA during early cyst development, suggesting that this protein helps to repress the translation of nanos mRNA." | females | Li, Y., Zhang, Q., Carreira-Rosario, A., Maines, J.Z., McKearin, D.M., and Buszczak, M. (2013). Mei-P26 Cooperates with Bam, Bgcn and Sxl to Promote Early Germline Development in the Drosophila Ovary. PLoS ONE 8, e58301. 10.1371/journal.pone.0058301. |
| "Mei-P26regulates PGC development" | females | Jankovics, F., Henn, L., Bujna, Á., Vilmos, P., Spirohn, K., Boutros, M., and Erdélyi, M. (2014). Functional Analysis of the Drosophila Embryonic Germ Cell Transcriptome by RNA Interference. PLoS ONE 9, e98579. 10.1371/journal.pone.0098579. |
| Mei-P26-Bgcn-Bam-Sxl-Brat-Ago1-miR980/miR-1 --\| nos 3'-UTR | females | Malik, S., Jang, W., and Kim, C. (2017). Protein Interaction Mapping of Translational Regulators Affecting Expression of the Critical Stem Cell Factor Nos. Development & Reproduction 21, 449–456. 10.12717/DR.2017.21.4.449. |

| | | |
|---|---|---|
| sisR-1, a stable intronic sequence RNA, negatively regulates mei-P26 in the GSC (through deadenylation) | females | Wong, J.T., Akhbar, F., Ng, A.Y.E., Tay, M.L.-I., Loi, G.J.E., and Pek, J.W. (2017). DIP1 modulates stem cell homeostasis in Drosophila through regulation of sisR-1. Nat Commun 8, 759. 10.1038/s41467-017-00684-4. |
| Wh regulates Mei-p26, and these proteins function together in multiple contexts to control GSC maintenance and differentiation | females | Rastegari, E., Kajal, K., Tan, B.-S., Huang, F., Chen, R.-H., Hsieh, T.-S., and Hsu, H.-J. (2020). WD40 protein Wuho controls germline homeostasis via TRIM-NHL tumor suppressor Mei-p26 in Drosophila. Development 147, dev182063. 10.1242/dev.182063. |
| Mei-P26 structure and RNA binding targets | females | Salerno-Kochan, A., Horn, A., Ghosh, P., Nithin, C., Kościelniak, A., Meindl, A., Strauss, D., Krutyhołowa, R., Rossbach, O., Bujnicki, J.M., et al. (2022). Molecular insights into RNA recognition and gene regulation by the TRIM-NHL protein Mei-P26. Life Sci. Alliance 5, e202201418. 10.26508/lsa.202201418. |
| Aub represses Mei-P26 translation through deadenylation | females | Rojas-Ríos, P., Chartier, A., Pierson, S., and Simonelig, M. (2017). Aubergine and pi RNA s promote germline stem cell self‐renewal by repressing the proto‐oncogene Cbl. EMBO J 36, 3194–3211. 10.15252/embj.201797259. |
| "mei-P26 mutant cystoblasts fail to downregulate dMyc protein, suggesting a role of Mei-P26 in dMyc repression during the stem cell-cystoblast transition" | females | Rhiner, C., Díaz, B., Portela, M., Poyatos, J.F., Fernández-Ruiz, I., López-Gay, J.M., Gerlitz, O., and Moreno, E. (2009). Persistent competition among stem cells and their daughters in the Drosophila ovary germline niche. Development 136, 995–1006. 10.1242/dev.033340. |
| Tut binds the long isoform of the mei-P26 3'-UTR; Bam binds Tut on its N-terminus and Bgcn on its C-Terminus to regulate Mei-P26 | males | Chen, D., Wu, C., Zhao, S., Geng, Q., Gao, Y., Li, X., Zhang, Y., and Wang, Z. (2014). Three RNA Binding Proteins Form a Complex to Promote Differentiation of Germline Stem Cell Lineage in Drosophila. PLoS Genet 10, e1004797. 10.1371/journal.pgen.1004797. |
| Mei-P26 facilitates the accumulation of Bam, and then Bam with Bgcn represses the translation of mei-P26 | males | Insco, M.L., Bailey, A.S., Kim, J., Olivares, G.H., Wapinski, O.L., Tam, C.H., and Fuller, M.T. (2012). A Self-Limiting Switch Based on Translational Control Regulates the Transition from Proliferation to Differentiation in an Adult Stem Cell Lineage. Cell Stem Cell 11, 689–700. 10.1016/j.stem.2012.08.012. |
| U2A is involved in mei-P26 splicing | males | Wu, H., Sun, L., Wen, Y., Liu, Y., Yu, J., Mao, F., Wang, Y., Tong, C., Guo, X., Hu, Z., et al. (2016). Major spliceosome defects cause male infertility and are associated with nonobstructive azoospermia in humans. Proc Natl Acad Sci USA 113, 4134–4139. 10.1073/pnas.1513682113. |
| Bruno binds to the sxl mRNA 3'-UTR to repress translation | females | Wang, Z., and Lin, H. (2007). Sex-lethal is a target of Bruno-mediated translational repression in promoting the differentiation of stem cell progeny during Drosophila oogenesis. Developmental Biology 302, 160–168. 10.1016/j.ydbio.2006.09.016. |
| Bam requires Sxl for differentiation | females | Chau, J., Kulnane, L.S., and Salz, H.K. (2009). Sex-lethal Facilitates the Transition From Germline Stem Cell to Committed Daughter Cell in the Drosophila Ovary. Genetics 182, 121–132. 10.1534/genetics.109.100693. |

| | | |
|---|---|---|
| Sxl is required for cell autonomous PGC fate deterimination | females | Hashiyama, K., Hayashi, Y., and Kobayashi, S. (2011). Drosophila Sex lethal Gene Initiates Female Development in Germline Progenitors. Science 333, 885–888. 10.1126/science.1208146. |
| Sxl binds the nos 3'-UTR to down-regulate Nos translation | females | Chau, J., Kulnane, L.S., and Salz, H.K. (2012). Sex-lethal enables germline stem cell differentiation by down-regulating Nanos protein levels during Drosophila oogenesis. Proceedings of the National Academy of Sciences 109, 9465–9470. 10.1073/pnas.1120473109. |
| Sxl review | females | Moschall, R., Gaik, M., and Medenbach, J. (2017). Promiscuity in post-transcriptional control of gene expression: Drosophila sex-lethal and its regulatory partnerships. FEBS Lett 591, 1471–1488. 10.1002/1873-3468.12652. |
| Sxl binds to 3'-UTRs to control the length distribution of all transcripts | females | Sandler, J.E., Irizarry, J., Stepanik, V., Dunipace, L., Amrhein, H., and Stathopoulos, A. (2018). A Developmental Program Truncates Long Transcripts to Temporally Regulate Cell Signaling. Developmental Cell 47, 773-784.e6. 10.1016/j.devcel.2018.11.019. |
| SXL functions with SETDB1 in the assembly of H3K9me3 silencing islands in germ cells | females | Smolko, A.E., Shapiro-Kulnane, L., and Salz, H.K. (2018). The H3K9 methyltransferase SETDB1 maintains female identity in Drosophila germ cells. Nat Commun 9, 4155. 10.1038/s41467-018-06697-x. |
| Sxl transcription is repressed by histone lysine methyltransferase (HKMT) Eggless (Egg/dSETDB1), which catalyzes methylation of Histone H3 lysine 9 (H3K9) | females | Clough, E., Tedeschi, T., and Hazelrigg, T. (2014). Epigenetic regulation of oogenesis and germ stem cell maintenance by the Drosophila histone methyltransferase Eggless/dSetDB1. Developmental Biology 388, 181–191. 10.1016/j.ydbio.2014.01.014. |
| Sxl alters poly-A lengths in the female germline | females | Gawande, B., Robida, M.D., Rahn, A., and Singh, R. (2006). Drosophila Sex-lethal protein mediates polyadenylation switching in the female germline. EMBO J 25, 1263–1272. 10.1038/sj.emboj.7601022. |
| wMel TomO rescues Sxl function in GSC maintenance through derepression of Nos translation | females | Ote, M., Ueyama, M., and Yamamoto, D. (2016). Wolbachia protein TomO targets nanos mRNA and restores germ stem cells in Drosophila sex-lethal mutants. Current Biology 26, 2223–2232. 10.1016/j.cub.2016.06.054. |
| wPip's TomO sequence rescues Sxl in D. melanogaster via nos translational derepression | females | Ote, M., and Yamamoto, D. (2018). Enhancing Nanos expression via the bacterial TomO protein is a conserved strategy used by the symbiont Wolbachia to fuel germ stem cell maintenance in infected Drosophila females. Archives of Insect Biochemistry and Physiology, e21471. 10.1002/arch.21471. |
| "TomO associates with orb mRNA, inhibiting interaction with the translation repressor Cup, leading to the precocious translation of Orb" | females | Ote, M., and Yamamoto, D. (2018). The Wolbachia protein TomO interacts with a host RNA to induce polarization defects in Drosophila oocytes. Archives of Insect Biochemistry and Physiology 99, e21475. 10.1002/arch.21475. |

**table S2.** Annotated references used to make fig S1A,B and the table in Fig. 9C.

**table S3**. Full fecundity dataset (n=3002). See “Table_S3_fecundity_assays - all_data.tsv” file.

| category | group1 | group2 | n1 | mean1 | n2 | mean2 | differential offspring/day: mean1-mean2 | proportion offspring/day: mean2/mean1 | test | p-value |
|---|---|---|---|---|---|---|---|---|---|---|
| wild type (WT) fertility | WT_OreR_wMel | WT_OreR_uninf | 115 | 25.72 | 164 | 24.75 | 0.96 | 0.96 | Wilcoxon rank sum | 8.09E-01 |
| | WT_OreR_uninf | WT_F10_OreR_uninf | "" | "" | 102 | 21.88 | 2.87 | 0.88 | Wilcoxon rank sum | 1.81E-01 |
| | WT_OreR_wMel | WT_F10_OreR_uninf | "" | "" | "" | "" | 3.84 | 0.85 | Wilcoxon rank sum | 1.23E-01 |
| | nos:Gal4>RFP_wMel | nos:Gal4>RFP_uninf | 23 | 23.03 | 21 | 31.58 | -8.55 | 1.37 | Wilcoxon rank sum | 1.25E-01 |
| | WT_OreR_wMel | nos:Gal4>RFP_wMel | "" | "" | "" | "" | -2.69 | 0.90 | Wilcoxon rank sum | 4.75E-01 |
| | WT_OreR_uninf | nos:Gal4>RFP_uninf | "" | "" | "" | "" | 6.83 | 1.28 | Wilcoxon rank sum | 1.05E-01 |
| | CyO/nos:Gal4_wMel | CyO/nos:Gal4_uninf | 83 | 40.26 | 55 | 27.09 | 13.18 | 0.67 | Wilcoxon rank sum | 2.20E-03 |
| | Sb/nos:Gal4_wMel | Sb/nos:Gal4_uninf | 43 | 34.39 | 26 | 31.93 | 2.46 | 0.93 | Wilcoxon rank sum | 7.41E-01 |
| F mei-P26 knockdown | nos:Gal4>meiP26RNAi_F_wMel | nos:Gal4>meiP26RNAi_F_uninf | 66 | 37.02 | 60 | 18.38 | 18.64 | 0.50 | Wilcoxon rank sum | 1.07E-04 |
| | mei-P26[1]_F_wMel | mei-P26[1]_F_uninf | 73 | 6.96 | 45 | 1.61 | 5.35 | 0.23 | Wilcoxon rank sum | 9.37E-05 |
| | mei-P26[1/mfs1]_F_wMel | mei-P26[1/mfs1]_F_uninf | 42 | 4.56 | 37 | 0.18 | 4.38 | 0.04 | Fisher's exact test (with/without_offspring) | 1.03E-05 |
| | mei-P26[mfs1]_F_wMel | mei-P26[mfs1]_F_uninf | 25 | 0.76 | 19 | 0.00 | 0.76 | 0.00 | Fisher's exact test (with/without_offspring) | 5.14E-06 |
| WT vs F mei-P26 knockdown | WT_OreR_wMel | nos:Gal4>meiP26RNAi_F_wMel | "" | "" | "" | "" | -11.30 | 1.44 | Wilcoxon rank sum | 1.15E-02 |
| | WT_OreR_uninf | nos:Gal4>meiP26RNAi_F_uninf | "" | "" | "" | "" | 6.38 | 0.74 | Wilcoxon rank sum | 1.26E-02 |
| | WT_OreR_wMel | nos:Gal4>meiP26RNAi_F_uninf | "" | "" | "" | "" | 7.34 | 0.71 | Wilcoxon rank sum | 1.24E-02 |
| | WT_OreR_uninf | nos:Gal4>meiP26RNAi_F_wMel | "" | "" | "" | "" | 12.26 | 1.50 | Wilcoxon rank sum | 5.78E-03 |
| | nos:Gal4>RFP_wMel | nos:Gal4>meiP26RNAi_F_wMel | "" | "" | "" | "" | -13.98 | 1.61 | Wilcoxon rank sum | 3.49E-02 |
| | nos:Gal4>RFP_uninf | nos:Gal4>meiP26RNAi_F_uninf | "" | "" | "" | "" | 13.20 | 0.58 | Wilcoxon rank sum | 5.42E-03 |
| | nos:Gal4>RFP_wMel | nos:Gal4>meiP26RNAi_F_uninf | "" | "" | "" | "" | 4.65 | 0.80 | Wilcoxon rank sum | 3.42E-01 |
| | nos:Gal4>RFP_uninf | nos:Gal4>meiP26RNAi_F_wMel | "" | "" | "" | "" | -5.44 | 1.17 | Wilcoxon rank sum | 7.18E-01 |
| | WT_OreR_uninf | mei-P26[1]_F_wMel | "" | "" | "" | "" | -12.26 | 1.50 | Wilcoxon rank sum | 5.78E-03 |
| | WT_OreR_wMel | mei-P26[1]_F_wMel | "" | "" | "" | "" | 18.76 | 0.27 | Wilcoxon rank sum | 3.45E-12 |
| | WT_OreR_uninf | mei-P26[1]_F_uninf | "" | "" | "" | "" | 23.15 | 0.06 | Wilcoxon rank sum | 2.26E-14 |
| | WT_OreR_wMel | mei-P26[1/mfs1]_F_wMel | "" | "" | "" | "" | 21.15 | 0.18 | Wilcoxon rank sum | 5.01E-10 |
| | WT_OreR_uninf | mei-P26[1/mfs1]_F_uninf | "" | "" | "" | "" | 24.57 | 0.01 | Wilcoxon rank sum | 3.40E-04 |
| | WT_OreR_wMel | mei-P26[mfs1]_F_wMel | "" | "" | "" | "" | 24.96 | 0.03 | Wilcoxon rank sum | 1.66E-09 |

| category | group1 | group2 | n1 | mean1 | n2 | mean2 | differential egg laid/day: mean1-mean2 | proportion egg laid/day: mean2/mean1 | test | **p-value** |
|---|---|---|---|---|---|---|---|---|---|---|
| | WT_OreR_uninf | mei-P26[mfs1]_F_uninf | "" | "" | "" | "" | 24.75 | 0.00 | Fisher's exact test (with/without_offspring) | 2.20E-16 |
| M mei-P26 knockdown | nos:Gal4>meiP26RNAi_M_wMel | nos:Gal4>meiP26RNAi_M_uninf | 35 | 36.11 | 35 | 32.45 | 3.67 | 0.90 | Wilcoxon rank sum | 4.41E-01 |
| | mei-P26[1]_M_wMel | mei-P26[1]_M_uninf | 32 | 43.38 | 21 | 32.49 | 10.89 | 0.75 | Wilcoxon rank sum | 1.58E-01 |
| | mei-P26[mfs1]_M_wMel | mei-P26[mfs1]_M_uninf | 21 | 24.84 | 24 | 0.40 | 24.45 | 0.02 | Wilcoxon rank sum | 2.85E-02 |
| WT vs M mei-P26 knockdown | WT_OreR_wMel | nos:Gal4>meiP26RNAi_M_wMel | "" | "" | "" | "" | -10.40 | 1.40 | Wilcoxon rank sum | 3.72E-03 |
| | WT_OreR_uninf | nos:Gal4>meiP26RNAi_M_uninf | "" | "" | "" | "" | -7.69 | 1.31 | Wilcoxon rank sum | 6.54E-02 |
| | WT_OreR_wMel | mei-P26[1]_M_wMel | "" | "" | "" | "" | -17.66 | 1.69 | Wilcoxon rank sum | 3.96E-04 |
| | WT_OreR_uninf | mei-P26[1]_M_uninf | "" | "" | "" | "" | -7.74 | 1.31 | Wilcoxon rank sum | 6.60E-02 |
| | WT_OreR_wMel | mei-P26[mfs1]_M_wMel | "" | "" | "" | "" | 0.88 | 0.97 | Wilcoxon rank sum | 4.35E-01 |
| | WT_OreR_uninf | mei-P26[mfs1]_M_uninf | "" | "" | "" | "" | 24.36 | 0.02 | Wilcoxon rank sum | 4.28E-04 |
| CI crosses | WT_OreR_Dmel_reciprocal-5d | WT_OreR_wMel | 60 | 22.50 | "" | "" | -3.21 | 1.14 | Wilcoxon rank sum | 2.74E-01 |
| | WT_OreR_Dmel_reciprocal-5d | WT_OreR_uninf | "" | "" | "" | "" | -2.25 | 1.10 | Wilcoxon rank sum | 3.25E-01 |
| | WT_OreR_Dmel_CI-0d | WT_OreR_Dmel_rescue-0d | 70 | 16.11 | 69 | 22.08 | -5.98 | 1.37 | Wilcoxon rank sum | 1.71E-02 |
| | WT_OreR_Dmel_CI-0d | WT_OreR_Dmel_reciprocal-0d | "" | "" | 14 | 29.69 | -13.59 | 1.84 | Wilcoxon rank sum | 1.06E-02 |
| | WT_OreR_Dmel_rescue-0d | WT_OreR_Dmel_reciprocal-0d | "" | "" | "" | "" | -7.61 | 1.34 | Wilcoxon rank sum | 1.71E-01 |
| | WT_OreR_Dmel_CI-5d | WT_OreR_Dmel_rescue-5d | 26 | 25.12 | 69 | 25.89 | -0.77 | 1.03 | Wilcoxon rank sum | 9.27E-01 |
| | WT_OreR_Dmel_CI-5d | WT_OreR_Dmel_reciprocal-5d | "" | "" | "" | "" | 2.62 | 0.90 | Wilcoxon rank sum | 3.63E-01 |
| | Dsimulans_CI-0d | Dsimulans_rescue-0d | 31 | 2.24 | 33 | 23.88 | -21.64 | 10.66 | Wilcoxon rank sum | 6.53E-07 |
| | Dsimulans_CI-5d | Dsimulans_rescue-5d | 17 | 7.51 | 27 | 19.00 | -11.49 | 2.53 | Wilcoxon rank sum | 5.00E-02 |

**table S4.** Fecundity statistics: offspring produced per female per day in single female-by-single male crosses. Experimental genotypes, infection statuses, and sexes are listed. The mate for each cross was OreR, of the same infection status, and of the opposite sex as the experimental fly. Males were aged 3-6 days, except for the young male CI crosses, which were aged zero days (distinguished with “-0d” and “-5d” labels). P-values <0.01 are in light green and <0.05 are in dark green for clarity.

| | | | | | | | | | | |
|---|---|---|---|---|---|---|---|---|---|---|
| wild type (WT) fertility | WT_OreR_wMel | WT_OreR_uninf | 115 | 27.49 | 164 | 27.69 | -0.20 | 1.01 | Wilcoxon rank sum | 9.09E-01 |
| | WT_OreR_uninf | WT_F10_OreR_uninf | "" | "" | 102 | 25.62 | 2.07 | 0.93 | Wilcoxon rank sum | 4.78E-01 |
| | WT_OreR_wMel | WT_F10_OreR_uninf | "" | "" | "" | "" | 1.87 | 0.93 | Wilcoxon rank sum | 5.18E-01 |
| | nos:Gal4>RFP_wMel | nos:Gal4>RFP_uninf | 23 | 23.42 | 21 | 34.87 | -11.46 | 1.49 | Wilcoxon rank sum | 7.22E-02 |
| | WT_OreR_wMel | nos:Gal4>RFP_wMel | "" | "" | "" | "" | -4.08 | 0.85 | Wilcoxon rank sum | 3.15E-01 |
| | WT_OreR_uninf | nos:Gal4>RFP_uninf | "" | "" | "" | "" | 7.18 | 1.26 | Wilcoxon rank sum | 1.15E-01 |
| | CyO/nos:Gal4_wMel | CyO/nos:Gal4_uninf | 83 | 42.36 | 55 | 28.43 | 13.93 | 0.67 | Wilcoxon rank sum | 2.22E-03 |
| | Sb/nos:Gal4_wMel | Sb/nos:Gal4_uninf | 43 | 34.21 | 26 | 34.17 | 0.03 | 1.00 | Wilcoxon rank sum | 1.00E+00 |
| F mei-P26 knockdown | nos:Gal4>meiP26RNAi_F_wMel | nos:Gal4>meiP26RNAi_F_uninf | 66 | 45.89 | 60 | 28.89 | 17.01 | 0.63 | Wilcoxon rank sum | 1.17E-03 |
| | mei-P26[1]_F_wMel | mei-P26[1]_F_uninf | 73 | 16.09 | 45 | 6.09 | 10.00 | 0.38 | Wilcoxon rank sum | 2.82E-02 |
| | mei-P26[1/mfs1]_F_wMel | mei-P26[1/mfs1]_F_uninf | 42 | 8.40 | 37 | 0.18 | 8.22 | 0.02 | Wilcoxon rank sum | 4.71E-05 |
| | mei-P26[mfs1]_F_wMel | mei-P26[mfs1]_F_uninf | 25 | 2.52 | 19 | 0.00 | 2.52 | 0.00 | Wilcoxon rank sum | 3.04E-05 |
| WT vs F mei-P26 knockdown | WT_OreR_wMel | nos:Gal4>meiP26RNAi_F_wMel | "" | "" | "" | "" | -18.40 | 1.67 | Wilcoxon rank sum | 6.99E-05 |
| | WT_OreR_uninf | nos:Gal4>meiP26RNAi_F_uninf | "" | "" | "" | "" | -1.20 | 1.04 | Wilcoxon rank sum | 5.15E-01 |
| | WT_OreR_wMel | nos:Gal4>meiP26RNAi_F_uninf | "" | "" | "" | "" | -1.40 | 1.05 | Wilcoxon rank sum | 5.29E-01 |
| | WT_OreR_uninf | nos:Gal4>meiP26RNAi_F_wMel | "" | "" | "" | "" | 18.20 | 1.66 | Wilcoxon rank sum | 3.01E-05 |
| | nos:Gal4>RFP_wMel | nos:Gal4>meiP26RNAi_F_wMel | "" | "" | "" | "" | -22.48 | 1.96 | Wilcoxon rank sum | 2.17E-03 |
| | nos:Gal4>RFP_uninf | nos:Gal4>meiP26RNAi_F_uninf | "" | "" | "" | "" | 5.99 | 0.83 | Wilcoxon rank sum | 1.68E-01 |
| | nos:Gal4>RFP_wMel | nos:Gal4>meiP26RNAi_F_uninf | "" | "" | "" | "" | -5.47 | 1.23 | Wilcoxon rank sum | 8.34E-01 |
| | nos:Gal4>RFP_uninf | nos:Gal4>meiP26RNAi_F_wMel | "" | "" | "" | "" | -11.02 | 1.32 | Wilcoxon rank sum | 1.76E-01 |
| | WT_OreR_uninf | mei-P26[1]_F_wMel | "" | "" | "" | "" | -18.20 | 1.66 | Wilcoxon rank sum | 3.01E-05 |
| | WT_OreR_wMel | mei-P26[1]_F_wMel | "" | "" | "" | "" | 11.40 | 0.59 | Wilcoxon rank sum | 5.13E-06 |
| | WT_OreR_uninf | mei-P26[1]_F_uninf | "" | "" | "" | "" | 21.60 | 0.22 | Wilcoxon rank sum | 1.03E-11 |
| | WT_OreR_wMel | mei-P26[1/mfs1]_F_wMel | "" | "" | "" | "" | 19.09 | 0.31 | Wilcoxon rank sum | 2.67E-10 |
| | WT_OreR_uninf | mei-P26[1/mfs1]_F_uninf | "" | "" | "" | "" | 27.51 | 0.01 | Wilcoxon rank sum | < 2.2e-16 |
| | WT_OreR_wMel | mei-P26[mfs1]_F_wMel | "" | "" | "" | "" | 24.97 | 0.09 | Wilcoxon rank sum | 1.12E-10 |
| | WT_OreR_uninf | mei-P26[mfs1]_F_uninf | "" | "" | "" | "" | 27.69 | 0.00 | Wilcoxon rank sum | 6.31E-11 |
| M mei-P26 knockdown | nos:Gal4>meiP26RNAi_M_wMel | nos:Gal4>meiP26RNAi_M_uninf | 38 | 34.84 | 41 | 36.30 | -1.46 | 1.04 | Wilcoxon rank sum | 9.10E-01 |
| | mei-P26[1]_M_wMel | mei-P26[1]_M_uninf | 35 | 34.64 | 28 | 33.47 | 1.18 | 0.97 | Wilcoxon rank sum | 9.83E-01 |
| | mei-P26[mfs1]_M_wMel | mei-P26[mfs1]_M_uninf | 21 | 28.79 | 28 | 2.51 | 26.29 | 0.09 | Wilcoxon rank sum | 6.10E-06 |

| | | | | | | | | | | |
|---|---|---|---|---|---|---|---|---|---|---|
| WT vs M mei-P26 knockdown | WT_OreR_wMel | nos:Gal4>meiP26RNAi_M_wMel | "" | "" | "" | "" | -7.35 | 1.27 | Wilcoxon rank sum | 3.58E-02 |
| | WT_OreR_uninf | nos:Gal4>meiP26RNAi_M_uninf | "" | "" | "" | "" | -8.61 | 1.31 | Wilcoxon rank sum | 3.90E-02 |
| | WT_OreR_wMel | mei-P26[1]_M_wMel | "" | "" | "" | "" | -7.15 | 1.26 | Wilcoxon rank sum | 2.32E-01 |
| | WT_OreR_uninf | mei-P26[1]_M_uninf | "" | "" | "" | "" | -5.78 | 1.21 | Wilcoxon rank sum | 1.20E-01 |
| | WT_OreR_wMel | mei-P26[mfs1]_M_wMel | "" | "" | "" | "" | -1.30 | 1.05 | Wilcoxon rank sum | 1.00E+00 |
| | WT_OreR_uninf | mei-P26[mfs1]_M_uninf | "" | "" | "" | "" | 25.18 | 0.09 | Wilcoxon rank sum | 8.59E-12 |

**table S5.** Fecundity statistics: eggs produced per female per day in single female-by-single male crosses. Experimental genotypes, infection statuses, and sexes are listed. The mate for each cross was OreR, of the same infection status, and of the opposite sex as the experimental fly. Males were aged 3-6 days, except for the young male CI crosses, which were aged zero days (distinguished with “-0d” and “-5d” labels). P-values <0.01 are in light green and <0.05 are in dark green for clarity.

| category | group1(lay>=20) | group2(lay>=20) | n1 | mean1 | n2 | mean2 | differential hatch: mean1-mean2 | proportion hatch/day: mean2/mean1 | test | p-value |
|---|---|---|---|---|---|---|---|---|---|---|
| wild type (WT) fertility | WT_OreR_wMel | WT_OreR_uninf | 76 | 88.40 | 104 | 83.18 | 5.22 | 0.94 | Wilcoxon rank sum | 3.46E-07 |
| | WT_OreR_uninf | WT_F10_OreR_uninf | "" | "" | 64 | 80.88 | 2.30 | 0.97 | Wilcoxon rank sum | 1.17E-01 |
| | WT_OreR_wMel | WT_F10_OreR_uninf | "" | "" | "" | "" | 7.51 | 0.92 | Wilcoxon rank sum | 8.98E-03 |
| | nos:Gal4>RFP_wMel | nos:Gal4>RFP_uninf | 12 | 88.49 | 16 | 90.50 | -2.01 | 1.02 | Wilcoxon rank sum | 9.82E-01 |
| | WT_OreR_wMel | nos:Gal4>RFP_wMel | "" | "" | "" | "" | 0.09 | 1.00 | Wilcoxon rank sum | 7.15E-01 |
| | WT_OreR_uninf | nos:Gal4>RFP_uninf | "" | "" | "" | "" | 7.32 | 1.09 | Wilcoxon rank sum | 5.01E-03 |
| | CyO/nos:Gal4_wMel | CyO/nos:Gal4_uninf | 66 | 91.31 | 31 | 88.83 | 2.47 | 0.97 | Wilcoxon rank sum | 2.08E-03 |
| | Sb/nos:Gal4_wMel | Sb/nos:Gal4_uninf | 33 | 88.66 | 22 | 89.55 | -0.89 | 1.01 | Wilcoxon rank sum | 8.37E-01 |
| F mei-P26 knockdown | nos:Gal4>meiP26RNAi_F_wMel | nos:Gal4>meiP26RNAi_F_uninf | 53 | 74.45 | 33 | 49.63 | 24.81 | 0.67 | Wilcoxon rank sum | 2.64E-08 |
| | mei-P26[1]_F_wMel | mei-P26[1]_F_uninf | 27(73) | 24.91 | 5(45) | 18.47 | 6.43 | 0.74 | Fisher's exact test (with/without_hatch) | 2.55E-03 |
| | mei-P26[1/mfs1]_F_wMel | mei-P26[1/mfs1]_F_uninf | 7(42) | 27.74 | 5(35) | 0.00 | 27.74 | 0.00 | Wilcoxon rank sum<br>(Fisher’s exact test) | 4.20E-03<br>(1.00E+00) |
| | mei-P26[mfs1]_F_wMel | mei-P26[mfs1]_F_uninf | 5(25) | 0.00 | 5(19) | 0.00 | 0.00 | 0.00 | NA - no females laid >=20 eggs/day | NA |
| WT vs F mei-P26 knockdown | WT_OreR_wMel | nos:Gal4>meiP26RNAi_F_wMel | "" | "" | "" | "" | 13.95 | 0.84 | Wilcoxon rank sum | 9.11E-07 |
| | WT_OreR_uninf | nos:Gal4>meiP26RNAi_F_uninf | "" | "" | "" | "" | 33.55 | 0.60 | Wilcoxon rank sum | 4.32E-15 |
| | WT_OreR_wMel | nos:Gal4>meiP26RNAi_F_uninf | "" | "" | "" | "" | 38.76 | 0.56 | Wilcoxon rank sum | 3.57E-14 |
| | WT_OreR_uninf | nos:Gal4>meiP26RNAi_F_wMel | "" | "" | "" | "" | -8.73 | 0.90 | Wilcoxon rank sum | 1.38E-02 |

| | | | | | | | | | | |
|---|---|---|---|---|---|---|---|---|---|---|
| | nos:Gal4>RFP_wMel | nos:Gal4>meiP26RNAi_F_wMel | "" | "" | "" | "" | 14.04 | 0.84 | Wilcoxon rank sum | 1.42E-02 |
| | nos:Gal4>RFP_uninf | nos:Gal4>meiP26RNAi_F_uninf | "" | "" | "" | "" | 40.86 | 0.55 | Wilcoxon rank sum | 1.93E-08 |
| | nos:Gal4>RFP_wMel | nos:Gal4>meiP26RNAi_F_uninf | "" | "" | "" | "" | 38.85 | 0.56 | Wilcoxon rank sum | 8.26E-07 |
| | nos:Gal4>RFP_uninf | nos:Gal4>meiP26RNAi_F_wMel | "" | "" | "" | "" | 16.05 | 0.82 | Wilcoxon rank sum | 3.56E-03 |
| | WT_OreR_uninf | mei-P26[1]_F_wMel | "" | "" | "" | "" | 8.73 | 0.90 | Wilcoxon rank sum | 1.38E-02 |
| | WT_OreR_wMel | mei-P26[1]_F_wMel | "" | "" | "" | "" | 63.49 | 0.28 | Wilcoxon rank sum | 4.82E-13 |
| | WT_OreR_uninf | mei-P26[1]_F_uninf | "" | "" | "" | "" | 64.71 | 0.22 | Wilcoxon rank sum | 2.93E-04 |
| | WT_OreR_wMel | mei-P26[1/mfs1]_F_wMel | "" | "" | "" | "" | 60.66 | 0.31 | Wilcoxon rank sum | 5.32E-05 |
| | WT_OreR_uninf | mei-P26[1/mfs1]_F_uninf | "" | "" | "" | "" | 83.18 | 0.00 | Wilcoxon rank sum | 1.97E-04 |
| | WT_OreR_wMel | mei-P26[mfs1]_F_wMel | "" | "" | "" | "" | 88.40 | 0.00 | Wilcoxon rank sum | 2.41E-04 |
| | WT_OreR_uninf | mei-P26[mfs1]_F_uninf | "" | "" | "" | "" | 83.18 | 0.00 | Wilcoxon rank sum | 1.97E-04 |
| M mei-P26 knockdown | nos:Gal4>meiP26RNAi_M_wMel | nos:Gal4>meiP26RNAi_M_uninf | 28 | 90.58 | 26 | 72.61 | 17.97 | 0.80 | Wilcoxon rank sum | 3.42E-03 |
| | mei-P26[1]_M_wMel | mei-P26[1]_M_uninf | 21 | 94.86 | 16 | 90.64 | 4.22 | 0.96 | Wilcoxon rank sum | 1.03E-01 |
| | mei-P26[mfs1]_M_wMel | mei-P26[mfs1]_M_uninf | 15 | 71.00 | 2 | 0.00 | 71.00 | 0.00 | Wilcoxon rank sum | 1.07E-01 |
| WT vs M mei-P26 knockdown | WT_OreR_wMel | nos:Gal4>meiP26RNAi_M_wMel | "" | "" | "" | "" | -2.18 | 1.02 | Wilcoxon rank sum | 1.65E-02 |
| | WT_OreR_uninf | nos:Gal4>meiP26RNAi_M_uninf | "" | "" | "" | "" | 10.57 | 0.87 | Wilcoxon rank sum | 9.00E-01 |
| | WT_OreR_wMel | mei-P26[1]_M_wMel | "" | "" | "" | "" | -6.46 | 1.07 | Wilcoxon rank sum | 4.72E-04 |
| | WT_OreR_uninf | mei-P26[1]_M_uninf | "" | "" | "" | "" | -7.46 | 1.09 | Wilcoxon rank sum | 5.00E-03 |
| | WT_OreR_wMel | mei-P26[mfs1]_M_wMel | "" | "" | "" | "" | 17.39 | 0.80 | Wilcoxon rank sum | 4.01E-01 |
| | WT_OreR_uninf | mei-P26[mfs1]_M_uninf | "" | "" | "" | "" | 83.18 | 0.00 | Wilcoxon rank sum | 1.73E-02 |
| CI crosses | WT_OreR_Dmel_reciprocal-5d | WT_OreR_wMel | 10 | 78.17 | "" | "" | -10.22 | 1.13 | Wilcoxon rank sum | 0.612 |
| | WT_OreR_Dmel_reciprocal-5d | WT_OreR_uninf | "" | "" | "" | "" | -5.01 | 1.06 | Wilcoxon rank sum | 9.91E-04 |
| | WT_OreR_Dmel_CI-0d | WT_OreR_Dmel_rescue-0d | 33 | 66.10 | 33 | 84.03 | -17.93 | 1.27 | Wilcoxon rank sum | 4.63E-04 |
| | WT_OreR_Dmel_CI-0d | WT_OreR_Dmel_reciprocal-0d | "" | "" | 9 | 93.21 | -27.11 | 1.41 | Wilcoxon rank sum | 4.56E-05 |
| | WT_OreR_Dmel_rescue-0d | WT_OreR_Dmel_reciprocal-0d | "" | "" | "" | "" | -9.18 | 1.11 | Wilcoxon rank sum | 1.25E-01 |
| | WT_OreR_Dmel_CI-5d | WT_OreR_Dmel_rescue-5d | 20 | 89.22 | 45 | 88.66 | 0.56 | 0.99 | Wilcoxon rank sum | 9.15E-01 |
| | WT_OreR_Dmel_CI-5d | WT_OreR_Dmel_reciprocal-5d | "" | "" | "" | "" | 11.05 | 0.88 | Wilcoxon rank sum | 2.09E-01 |
| | WT_OreR_Dmel_rescue-5d | WT_OreR_Dmel_reciprocal-5d | "" | "" | "" | "" | 10.49 | 0.88 | Wilcoxon rank sum | 9.72E-02 |
| | Dsimulans_CI-0d | Dsimulans_rescue-0d | 24 | 6.07 | 28 | 68.43 | -62.36 | 11.27 | Wilcoxon rank sum | 5.64E-08 |
| | Dsimulans_CI-5d | Dsimulans_rescue-5d | 10 | 26.52 | 17 | 69.04 | -42.51 | 2.60 | Wilcoxon rank sum | 3.63E-04 |

**table S6.** Fecundity statistics: percentage of eggs that hatched from single female-by-single male crosses that laid >= 20 eggs. Experimental genotypes, infection statuses, and sexes are listed. The mate for each cross was OreR, of the same infection status, and of the opposite sex as the experimental fly. Males were aged 3-6 days, except for the young male CI crosses, which were aged zero days (distinguished with “-0d” and “-5d” labels). Sample counts (n1,n2) in parentheses are for Fisher Exact Tests (samples with hatched eggs vs no hatched eggs, opposed to % hatch for samples with 20 or more eggs laid). P-values <0.01 are in light green and <0.05 are in dark green for clarity.

| group1 | group2 | test | egg lay (#/female/day) p-value | %egg hatch (>10 eggs laid) p-value | offspring (#/female/day) p-value |
|---|---|---|---|---|---|
| WT_OreR_wMel | WT_OreR_uninf | Kolmogorov-Smirnov | 7.84E-02 | 7.06E-09 | 6.02E-02 |
| meiP26RNAi_F_wMel | meiP26RNAi_F_uninf | Kolmogorov-Smirnov | 5.92E-06 | 1.48E-11 | 3.87E-08 |
| meiP261_F_wMel | meiP261_F_uninf | Kolmogorov-Smirnov | 3.41E-02 | 6.36E-05 | 1.18E-04 |

**table S7.** Fecundity versus age statistics

| category | group1 | group2 | n1 | mean1 | n2 | mean2 | test | **p-value** |
|---|---|---|---|---|---|---|---|---|
| wild type (WT) | WT_OreR_wMel-5d | WT_OreR_uninf-5d | 8 | 2.313 | 13 | 1.923 | Wilcoxon rank sum | 1.81E-01 |
| | WT_OreR_wMel-10d | WT_OreR_uninf-10d | 15 | 1.600 | 15 | 1.867 | Wilcoxon rank sum | 3.47E-01 |
| | WT_OreR_wMel-31d | WT_OreR_uninf-31d | 14 | 1.321 | 15 | 1.533 | Wilcoxon rank sum | 7.64E-01 |
| | WT_OreR_wMel-5d | WT_OreR_wMel-10d | "" | "" | "" | "" | Wilcoxon rank sum | 1.93E-02 |
| | WT_OreR_uninf-5d | WT_OreR_uninf-10d | "" | "" | "" | "" | Wilcoxon rank sum | 8.24E-01 |
| | WT_OreR_wMel-10d | WT_OreR_wMel-31d | "" | "" | "" | "" | Wilcoxon rank sum | 0.2655 |
| | WT_OreR_uninf-10d | WT_OreR_uninf-31d | "" | "" | "" | "" | Wilcoxon rank sum | 0.249 |
| F mei-P26 knockdown | meiP26RNAi_F_wMel-5d | meiP26RNAi_F_uninf-5d | 31 | 2.048 | 23 | 1.848 | Wilcoxon rank sum | 6.68E-01 |
| | meiP261_F_wMel-5d | meiP261_F_uninf-5d | 42 | 1.024 | 73 | 0.575 | Wilcoxon rank sum | 2.88E-04 |
| | meiP26RNAi_F_wMel-5d | meiP261_F_wMel-5d | "" | "" | "" | "" | Wilcoxon rank sum | 4.47E-08 |
| | meiP26RNAi_F_wMel-5d | meiP261_F_uninf-5d | "" | "" | "" | "" | Wilcoxon rank sum | 4.04E-14 |
| | meiP26RNAi_F_uninf-5d | meiP261_F_uninf-5d | "" | "" | "" | "" | Wilcoxon rank sum | 3.63E-08 |
| | meiP26RNAi_F_uninf-5d | meiP261_F_wMel-5d | "" | "" | "" | "" | Wilcoxon rank sum | 3.15E-04 |
| | meiP261_F_wMel-10d | meiP261_F_uninf-10d | 14 | 1.464 | 16 | 0.938 | Wilcoxon rank sum | 1.48E-02 |
| | meiP261_F_wMel-5d | meiP261_F_wMel-10d | "" | "" | "" | "" | Wilcoxon rank sum | 1.32E-02 |
| | meiP261_F_uninf-5d | meiP261_F_uninf-10d | "" | "" | "" | "" | Wilcoxon rank sum | 1.37E-02 |
| WT vs F mei-P26 knockdown | WT_OreR_wMel-5d | meiP26RNAi_F_wMel-5d | "" | "" | "" | "" | Wilcoxon rank sum | 1.94E-01 |
| | WT_OreR_uninf-5d | meiP26RNAi_F_uninf-5d | "" | "" | "" | "" | Wilcoxon rank sum | 1.00E+00 |
| | WT_OreR_uninf-5d | meiP26RNAi_F_wMel-5d | "" | "" | "" | "" | Wilcoxon rank sum | 6.12E-01 |
| | WT_OreR_wMel-5d | meiP26RNAi_F_uninf-5d | "" | "" | "" | "" | Wilcoxon rank sum | 2.33E-01 |
| | WT_OreR_uninf-5d | meiP261_F_uninf-5d | "" | "" | "" | "" | Wilcoxon rank sum | 3.18E-07 |
| | WT_OreR_wMel-5d | meiP261_F_wMel-5d | "" | "" | "" | "" | Wilcoxon rank sum | 1.18E-04 |
| | WT_OreR_wMel-5d | meiP261_F_uninf-5d | "" | "" | "" | "" | Wilcoxon rank sum | 3.36E-06 |
| | WT_OreR_uninf-5d | meiP261_F_wMel-5d | "" | "" | "" | "" | Wilcoxon rank sum | 2.75E-04 |
| | WT_OreR_wMel-10d | meiP261_F_wMel-10d | "" | "" | "" | "" | Wilcoxon rank sum | 6.66E-01 |
| | WT_OreR_wMel-10d | meiP261_F_uninf-10d | "" | "" | "" | "" | Wilcoxon rank sum | 1.10E-02 |
| | WT_OreR_uninf-10d | meiP261_F_wMel-10d | "" | "" | "" | "" | Wilcoxon rank sum | 1.40E-01 |
| | WT_OreR_uninf-10d | meiP261_F_uninf-10d | "" | "" | "" | "" | Wilcoxon rank sum | 1.24E-03 |

**table S8.** Germline stem cell (GSC) counts per germarium.

| category | group1 | group2 | n germaria | n GSCs | n pHH3+ GSCs | proportion GSCs mitotic | n germaria | n GSCs | n pHH3+ GSCs | proportion GSCs mitotic | test | p-value |
|---|---|---|---|---|---|---|---|---|---|---|---|---|
| wild type (WT) | WT_OreR_wMel | WT_OreR_uninf | 81 | 179 | 10.000 | 0.056 | 68 | 143 | 9 | 0.063 | Fisher's Exact Test | 8.16E-01 |
| mei-P26 knockdown | meiP261_F_wMel | meiP261_F_uninf | 111 | 152 | 9.000 | 0.059 | 111 | 115 | 1 | 0.009 | Fisher's Exact Test | 4.69E-02 |
| WT vs F mei-P26 knockdown | WT_OreR_uninf | meiP261_F_uninf | "" | "" | | "" | "" | "" | | "" | Fisher's Exact Test | 4.62E-02 |
| | WT_OreR_wMel | meiP261_F_wMel | "" | "" | | "" | "" | "" | | "" | Fisher's Exact Test | 1.00E+00 |
| | WT_OreR_wMel | meiP261_F_uninf | "" | "" | | "" | "" | "" | | "" | Fisher's Exact Test | 5.51E-02 |
| | WT_OreR_uninf | meiP261_F_wMel | "" | "" | | "" | "" | "" | | "" | Fisher's Exact Test | 1.00E+00 |

**table S9.** Number of GSCs in mitosis (anti-pHH3-positive staining), per germarium.

| category | group1 | group2 | n germaria | avg # mitotic cystocytes / germarium | n germaria | avg # mitotic cystocytes / germarium | test | **p-value** |
|---|---|---|---|---|---|---|---|---|
| wild type (WT) | WT_OreR_wMel | WT_OreR_uninf | 81 | 1.079 | 68 | 1.957 | Wilcoxon Rank Sum Test | 3.01E-01 |
| mei-P26 knockdown | meiP261_F_wMel | meiP261_F_uninf | 111 | 2.545 | 111 | 1.817 | Fisher's Exact Test | 1.40E-01 |
| WT vs F mei-P26 knockdown | WT_OreR_uninf | meiP261_F_uninf | "" | "" | "" | "" | Fisher's Exact Test | 9.52E-01 |
| | WT_OreR_wMel | meiP261_F_wMel | "" | "" | "" | "" | Fisher's Exact Test | 1.34E-02 |
| | WT_OreR_wMel | meiP261_F_uninf | "" | "" | "" | "" | Fisher's Exact Test | 2.64E-01 |
| | WT_OreR_uninf | meiP261_F_wMel | "" | "" | "" | "" | Fisher's Exact Test | 2.26E-01 |

**table S10.** Number of cystocytes (CC) in mitosis (anti-pHH3-positive staining), per germarium.

| category | group1 | group2 | n1 | n2 | test | region 1 - relative fluor mean1 | region 1 - relative fluor mean2 | differential region 1 - \|mean1-mean2\| | relative fluor region 1 p-value | region 2a - relative fluor mean1 | region 2a - relative fluor mean2 | differential region 2a - \|mean1-mean2\| | relative fluor region 2a p-value | region 2b - relative fluor mean1 | region 2b - relative fluor mean2 | differential region 2b - \|mean1-mean2\| | relative fluor region 2b p-value |
|---|---|---|---|---|---|---|---|---|---|---|---|---|---|---|---|---|---|
| wild type (WT) | WT_OreR_wMel-5d | WT_OreR_uninf-5d | 26 | 40 | Wilcoxon rank sum | 0.644 | 0.639 | 0.005 | 5.02E-01 | 0.236 | 0.197 | 0.039 | 1.42E-02 | 0.172 | 0.155 | 0.02 | 2.64E-01 |
| F mei-P26 knockdown | meiP261_F_wMel-5d | meiP261_F_uninf-5d | 36 | 39 | Wilcoxon rank sum | 0.527 | 0.466 | 0.061 | 8.40E-03 | 0.222 | 0.236 | 0.014 | 1.77E-01 | 0.211 | 0.262 | 0.05 | 1.72E-02 |
| WT vs F mei-P26 knockdown | WT_OreR_uninf-5d | meiP261_F_uninf-5d | | | Wilcoxon rank sum | | | 0.173 | 5.67E-10 | | | 0.039 | 8.60E-04 | | | 0.11 | 2.30E-07 |
| | WT_OreR_wMel-5d | meiP261_F_wMel-5d | | | Wilcoxon rank sum | | | 0.117 | 2.87E-04 | | | 0.014 | 8.82E-01 | | | 0.04 | 2.21E-02 |
| | WT_OreR_wMel-5d | meiP261_F_uninf-5d | | | Wilcoxon rank sum | | | 0.178 | 9.90E-08 | | | 0.000 | 3.08E-01 | | | 0.09 | 5.33E-05 |
| | WT_OreR_uninf-5d | meiP261_F_wMel-5d | | | Wilcoxon rank sum | | | 0.112 | 3.78E-06 | | | 0.025 | 9.10E-02 | | | 0.06 | 1.16E-03 |

**table S11.** Sxl expression by germarium region, measured by fluorescence intensity.

| category | group1 | group2 | n1 | n2 | test | GSC - relative fluor mean 1 | GSC - relative fluor mean2 | differential GSC - \|mean1-mean2\| | relative fluor GSC p-value | CB - relative fluor mean1 | CB - relative fluor mean2 | differential CB - \|mean1-mean2\| | relative fluor CB p-value | region 2a - relative fluor mean1 | region 2a - relative fluor mean2 | differential region 2a - \|mean1-mean2\| | relative fluor region 2a p-value | region 2b - relative fluor mean1 | region 2b - relative fluor mean2 | differential region 2b - \|mean1-mean2\| | relative fluor region 2b p-value |
|---|---|---|---|---|---|---|---|---|---|---|---|---|---|---|---|---|---|---|---|---|---|
| wild type (WT) | WT_OreR_wMel-5d | WT_OreR_uninf-5d | 33 | 29 | Wilcoxon rank sum | 0.102 | 0.089 | 0.012 | 1.72E-01 | 0.481 | 0.469 | 0.011 | 6.14E-01 | 0.174 | 0.195 | 0.021 | 1.72E-01 | 0.206 | 0.204 | 0.002 | 7.05E-01 |
| F mei-P26 knockdown | meiP261_F_wMel-5d | meiP261_F_uninf-5d | 34 | 39 | Wilcoxon rank sum | 0.056 | 0.069 | 0.012 | 4.25E-01 | 0.502 | 0.397 | 0.105 | 1.60E-02 | 0.179 | 0.220 | 0.041 | 6.21E-02 | 0.210 | 0.263 | 0.053 | 1.66E-01 |
| WT vs F mei-P26 knockdown | WT_OreR_uninf-5d | meiP261_F_uninf-5d | | | Wilcoxon rank sum | | | 0.021 | 1.42E-03 | | | 0.073 | 9.18E-02 | | | 0.025 | 1.74E-01 | | | 0.059 | 3.33E-02 |
| | WT_OreR_wMel-5d | meiP261_F_wMel-5d | | | Wilcoxon rank sum | | | 0.046 | 1.28E-06 | | | 0.021 | 4.58E-01 | | | 0.005 | 7.70E-01 | | | 0.004 | 6.14E-01 |
| | WT_OreR_wMel-5d | meiP261_F_uninf-5d | | | Wilcoxon rank sum | | | 0.033 | 4.32E-05 | | | 0.084 | 3.40E-02 | | | 0.046 | 1.99E-02 | | | 0.057 | 3.50E-02 |
| | WT_OreR_uninf-5d | meiP261_F_wMel-5d | | | Wilcoxon rank sum | | | 0.033 | 5.97E-05 | | | 0.033 | 3.27E-01 | | | 0.016 | 3.27E-01 | | | 0.006 | 5.15E-01 |

**table S12.** Bam expression by germarium region, measured by fluorescence intensity.

| category | group1 | group2 | n1 | n2 | test | GSC - relative Bam/pMad fluor mean1 | GSC - relative Bam/pMad fluor mean2 | differential Bam/pMad GSC - \|mean1-mean2\| | relative Bam/pMad fluor GSC p-value | CB - relative Bam/pMad fluor mean1 | CB - relative Bam/pMad fluor mean2 | differential Bam/pMad CB - \|mean1-mean2\| | relative Bam/pMad fluor CB p-value |
|---|---|---|---|---|---|---|---|---|---|---|---|---|---|
| wild type (WT) | WT_OreR_wMel-5d | WT_OreR_uninf-5d | 21 | 29 | Wilcoxon rank sum | 0.467 | 0.593 | 0.126 | 4.43E-02 | 1.841 | 1.767 | 0.074 | 5.11E-02 |
| F mei-P26 knockdown | meiP261_F_wMel-5d | meiP261_F_uninf-5d | 20 | 27 | Wilcoxon rank sum | 0.640 | 1.422 | 0.783 | 9.82E-03 | 2.814 | 2.142 | 0.673 | 1.43E-03 |
| WT vs F mei-P26 knockdown | WT_OreR_uninf-5d | meiP261_F_uninf-5d | | | Wilcoxon rank sum | | | 0.829 | 4.04E-04 | | | 0.374 | 2.82E-01 |
| | WT_OreR_wMel-5d | meiP261_F_wMel-5d | | | Wilcoxon rank sum | | | 0.173 | 9.08E-01 | | | 0.973 | 2.13E-02 |
| | WT_OreR_wMel-5d | meiP261_F_uninf-5d | | | Wilcoxon rank sum | | | 0.955 | 2.51E-04 | | | 0.300 | 3.86E-01 |
| | WT_OreR_uninf-5d | meiP261_F_wMel-5d | | | Wilcoxon rank sum | | | 0.047 | 8.57E-02 | | | 1.047 | 1.22E-06 |

**table S13.** Relative Bam vs pMad expression, measured by fluorescence, in GSCs.

| | | | | nurse cells in cyst | | | | nurse cells in cyst | | | | |
|---|---|---|---|---|---|---|---|---|---|---|---|---|
| category | group1 | group2 | n1 | <15 | 15 | >15 | n2 | <15 | 15 | >15 | test | p-value |
| wild type (WT) | WT_nos:Gal4_wMel | WT_nos:Gal4_uninf | 215 | 0 | 212 | 3 | 216 | 0 | 216 | 0 | Fisher's exact test | 1.23E-01 |
| F mei-P26 knockdown | meiP26RNAi_F_wMel | meiP26RNAi_F_uninf | 86 | 10 | 62 | 14 | 190 | 16 | 107 | 67 | Fisher's exact test | 4.46E-03 |
| | meiP261_F_wMel | meiP261_F_uninf | 444 | 37 | 189 | 218 | 124 | 8 | 47 | 69 | Fisher's exact test | 4.44E-01 |
| | meiP26RNAi_F_wMel | meiP261_F_wMel | "" | "" | "" | "" | "" | "" | "" | "" | Fisher's exact test | 2.24E-08 |
| | meiP26RNAi_F_wMel | meiP261_F_uninf | "" | "" | "" | "" | "" | "" | "" | "" | Fisher's exact test | 1.81E-08 |
| | meiP26RNAi_F_uninf | meiP261_F_uninf | "" | "" | "" | "" | "" | "" | "" | "" | Fisher's exact test | 1.63E-03 |
| | meiP26RNAi_F_uninf | meiP261_F_wMel | "" | "" | "" | "" | "" | "" | "" | "" | Fisher's exact test | 3.87E-03 |
| WT vs F mei-P26 knockdown | WT_nos:Gal4_wMel | meiP26RNAi_F_wMel | "" | "" | "" | "" | "" | "" | "" | "" | Fisher's exact test | 2.72E-12 |
| | WT_nos:Gal4_uninf | meiP26RNAi_F_uninf | "" | "" | "" | "" | "" | "" | "" | "" | Fisher's exact test | < 2.2e-16 |
| | WT_nos:Gal4_uninf | meiP26RNAi_F_wMel | "" | "" | "" | "" | "" | "" | "" | "" | Fisher's exact test | 5.94E-15 |
| | WT_nos:Gal4_wMel | meiP26RNAi_F_uninf | "" | "" | "" | "" | "" | "" | "" | "" | Fisher's exact test | < 2.2e-16 |
| | WT_nos:Gal4_uninf | meiP261_F_uninf | "" | "" | "" | "" | "" | "" | "" | "" | Fisher's exact test | < 2.2e-16 |
| | WT_nos:Gal4_wMel | meiP261_F_wMel | "" | "" | "" | "" | "" | "" | "" | "" | Fisher's exact test | < 2.2e-16 |
| | WT_nos:Gal4_wMel | meiP261_F_uninf | "" | "" | "" | "" | "" | "" | "" | "" | Fisher's exact test | < 2.2e-16 |
| | WT_nos:Gal4_uninf | meiP261_F_wMel | "" | "" | "" | "" | "" | "" | "" | "" | Fisher's exact test | < 2.2e-16 |
| F mei-P26 OE | meiP26OE_F_wMel | meiP26OE_F_uninf | 101 | 7 | 85 | 9 | 138 | 0 | 131 | 7 | Fisher's exact test | 1.63E-03 |
| mei-P26 OE vs mei-P26 knockdown | meiP26OE_F_wMel | meiP26RNAi_F_wMel | "" | "" | "" | "" | "" | "" | "" | "" | Fisher's exact test | 1.36E-01 |
| | meiP26OE_F_wMel | meiP261_F_wMel | "" | "" | "" | "" | "" | "" | "" | "" | Fisher's exact test | 1.31E-15 |
| | meiP26OE_F_wMel | meiP26RNAi_F_uninf | "" | "" | "" | "" | "" | "" | "" | "" | Fisher's exact test | 6.80E-07 |
| | meiP26OE_F_wMel | meiP261_F_uninf | "" | "" | "" | "" | "" | "" | "" | "" | Fisher's exact test | 2.87E-14 |
| | meiP26OE_F_uninf | meiP26RNAi_F_wMel | "" | "" | "" | "" | "" | "" | "" | "" | Fisher's exact test | 7.96E-07 |
| | meiP26OE_F_uninf | meiP261_F_wMel | "" | "" | "" | "" | "" | "" | "" | "" | Fisher's exact test | < 2.2e-16 |
| | meiP26OE_F_uninf | meiP26RNAi_F_uninf | "" | "" | "" | "" | "" | "" | "" | "" | Fisher's exact test | 4.19E-16 |
| | meiP26OE_F_uninf | meiP261_F_uninf | "" | "" | "" | "" | "" | "" | "" | "" | Fisher's exact test | < 2.2e-16 |
| WT vs F mei-P26 OE | meiP26OE_F_wMel | WT_nos:Gal4_wMel | "" | "" | "" | "" | "" | "" | "" | "" | Fisher's exact test | 1.23E-06 |
| | meiP26OE_F_wMel | WT_nos:Gal4_uninf | "" | "" | "" | "" | "" | "" | "" | "" | Fisher's exact test | 4.73E-09 |
| | meiP26OE_F_uninf | WT_nos:Gal4_wMel | "" | "" | "" | "" | "" | "" | "" | "" | Fisher's exact test | 5.24E-02 |
| | meiP26OE_F_uninf | WT_nos:Gal4_uninf | "" | "" | "" | "" | "" | "" | "" | "" | Fisher's exact test | 1.24E-03 |

**table S14.** Tumorous germline cyst counts. Normal cysts contain 16 germline-derived cells, 15 nurse cells and one oocyte. Cysts containing greater or less than 15 nurse cells were scored as tumorous or abnormal, respectively.

| category | group1 | group2 | n1 | Y | M | N | n2 | Y | M | N | test | p-value |
|---|---|---|---|---|---|---|---|---|---|---|---|---|
| wild type (WT) | WT_OreR_wMel-5d | WT_OreR_uninf-5d | 53 | 50 | 2 | 1 | 64 | 58 | 6 | 0 | Fisher's Exact Test | 2.09E-01 |
| F mei-P26 knockdown | meiP261_F_wMel-5d | meiP261_F_uninf-5d | 66 | 40 | 15 | 11 | 63 | 16 | 4 | 43 | Fisher's Exact Test | 5.85E-09 |
| WT vs F mei-P26 knockdown | WT_OreR_uninf-5d | meiP261_F_uninf-5d | "" | "" | "" | "" | "" | "" | "" | "" | Fisher's Exact Test | < 2.2e-16 |
| | WT_OreR_wMel-5d | meiP261_F_wMel-5d | "" | "" | "" | "" | "" | "" | "" | "" | Fisher's Exact Test | 5.61E-05 |
| | WT_OreR_wMel-5d | meiP261_F_uninf-5d | "" | "" | "" | "" | "" | "" | "" | "" | Fisher's Exact Test | 4.51E-16 |
| | WT_OreR_uninf-5d | meiP261_F_wMel-5d | "" | "" | "" | "" | "" | "" | "" | "" | Fisher's Exact Test | 2.97E-05 |

**table S15.** Counts of germline cysts exhibiting oocyte-specific Orb expression (Y), unclear staining (M), or no specific expression, indicating developmentally abnormal cysts lacking specified oocytes.

| protocol | Sample ID | # Read pairs | Yield (Mbases) | Mean Quality Score | % Bases >= 30 | Dmel mapped reads | Dmel % | Dmel coverage (calculated) | wMel mapped reads | wMel % | wMel coverage (calculated) |
|---|---|---|---|---|---|---|---|---|---|---|---|
| ribodepletion | mei-P261uninfected-1 | 16566114 | 4970 | 35.28 | 90.31 | 26794060 | 80.87% | 21.73 | 142 | 0.00% | 0.02 |
| ribodepletion | mei-P261uninfected-2 | 24371726 | 7312 | 35.23 | 90.06 | 39374606 | 80.78% | 31.93 | 1644 | 0.00% | 0.22 |
| ribodepletion | mei-P261uninfected-3 | 20726752 | 6218 | 35.2 | 89.96 | 31465602 | 75.91% | 25.52 | 130 | 0.00% | 0.02 |
| ribodepletion | mei-P261uninfected-4 | 22725501 | 6818 | 34.99 | 89.1 | 33541934 | 73.80% | 27.20 | 4742 | 0.01% | 0.65 |
| ribodepletion | mei-P261uninfected-5 | 25921666 | 7776 | 35.29 | 90.4 | 41658014 | 80.35% | 33.79 | 92 | 0.00% | 0.01 |
| ribodepletion | mei-P261uninfected-6 | 21289976 | 6387 | 35.38 | 90.8 | 32541844 | 76.43% | 26.39 | 48 | 0.00% | 0.01 |
| ribodepletion | mei-P261wMel-1 | 30205483 | 9061 | 32.9 | 80.16 | 36362650 | 60.19% | 29.49 | 57237 | 0.09% | 7.83 |
| ribodepletion | mei-P261wMel-3 | 22849556 | 6855 | 35.36 | 90.65 | 35184986 | 76.99% | 28.54 | 62646 | 0.14% | 8.57 |
| ribodepletion | mei-P261wMel-4 | 26232731 | 7870 | 35.37 | 90.71 | 42053352 | 80.15% | 34.11 | 118236 | 0.23% | 16.18 |
| ribodepletion | mei-P261wMel-5 | 23786557 | 7136 | 35.41 | 90.93 | 39184612 | 82.37% | 31.78 | 87424 | 0.18% | 11.96 |
| ribodepletion | mei-P261wMel-7 | 26403318 | 7921 | 35.34 | 90.59 | 41693804 | 78.96% | 33.82 | 96438 | 0.18% | 13.20 |
| ribodepletion | mei-P261wMel-8 | 24804635 | 7441 | 35.33 | 90.55 | 39188718 | 78.99% | 31.78 | 80004 | 0.16% | 10.95 |
| ribodepletion | OreRF10uninfected-4 | 21514878 | 6454 | 35.41 | 90.95 | 33544436 | 77.96% | 27.21 | 1670 | 0.00% | 0.23 |
| ribodepletion | OreRF10uninfected-5 | 21887003 | 6566 | 35.4 | 90.9 | 33751420 | 77.10% | 27.37 | 978 | 0.00% | 0.13 |
| ribodepletion | OreRF10uninfected-6 | 22033197 | 6610 | 35.13 | 89.72 | 32014844 | 72.65% | 25.97 | 926 | 0.00% | 0.13 |
| ribodepletion | OreRuninfected-1 | 22902974 | 6871 | 35.42 | 91 | 38142430 | 83.27% | 30.94 | 1000 | 0.00% | 0.14 |
| ribodepletion | OreRuninfected-2 | 19424258 | 5827 | 35.3 | 90.37 | 32687632 | 84.14% | 26.51 | 556 | 0.00% | 0.08 |
| ribodepletion | OreRuninfected-3 | 25244059 | 7573 | 35.25 | 90.13 | 39497260 | 78.23% | 32.03 | 576 | 0.00% | 0.08 |
| ribodepletion | OreRwMelDB-1 | 23669962 | 7101 | 35.36 | 90.66 | 40049364 | 84.60% | 32.48 | 69074 | 0.15% | 9.45 |
| ribodepletion | OreRwMelDB-2 | 21342747 | 6403 | 35.28 | 90.3 | 31878812 | 74.68% | 25.86 | 129552 | 0.30% | 17.73 |
| ribodepletion | OreRwMelDB-3 | 20997126 | 6299 | 35.33 | 90.52 | 32964512 | 78.50% | 26.74 | 272124 | 0.65% | 37.23 |
| ribodepletion | OreRwMelDB-4 | 26137488 | 7841 | 35.31 | 90.46 | 44312894 | 84.77% | 35.94 | 186630 | 0.36% | 25.54 |
| ribodepletion | OreRwMelDB-6 | 23976948 | 7193 | 35.38 | 90.75 | 38740764 | 80.79% | 31.42 | 310028 | 0.65% | 42.42 |
| ribodepletion | OreRwMelDB-7 | 23816174 | 7145 | 35.37 | 90.75 | 37387940 | 78.49% | 30.32 | 228366 | 0.48% | 31.25 |
| poly-A | nos-meiP26RNAi-wMel -1-resub | 72190715 | 21657 | 35.97 | 93.99 | 137328600 | 95.12% | 111.38 | 3948 | 0.00% | 0.54 |
| poly-A | nos-meiP26RNAi-wMel -2-resub | 65207225 | 19562 | 35.96 | 93.99 | 123616024 | 94.79% | 100.26 | 2204 | 0.00% | 0.30 |

| | | | | | | | | | | | |
|---|---|---|---|---|---|---|---|---|---|---|---|
| poly-A | nos-meiP26RNAi-wMel -3 | 67418318 | 20225 | 35.87 | 93.54 | 126461294 | 93.79% | 102.57 | 6748 | 0.01% | 0.92 |
| poly-A | nosGal4CyO-wMel-1 | 63388028 | 19016 | 35.92 | 93.81 | 120645078 | 95.16% | 97.85 | 7462 | 0.01% | 1.02 |
| poly-A | nosGal4CyO-wMel-2 | 71148881 | 21345 | 35.81 | 93.18 | 133691906 | 93.95% | 108.43 | 5078 | 0.00% | 0.69 |
| poly-A | nosGal4CyO-wMel-3-re sub | 67506892 | 20252 | 35.96 | 93.95 | 128140108 | 94.91% | 103.93 | 3636 | 0.00% | 0.50 |

**table S16.** Transcriptomic dataset generated to test the impacts of mei-P26 knockdown and wMel infection. Data deposited under NCBI BioProjectnumber PRJNA992140.

| gene_id | baseMean | log2FoldChange | lfcSE | stat | pvalue | padj |
|---|---|---|---|---|---|---|
| Dmel_CG32581 | 498.502 | -7.545 | 0.348 | -21.705 | 1.85E-104 | 1.97E-100 |
| Dmel_CG12218 | 12376.792 | 5.652 | 0.285 | 19.841 | 1.31E-87 | 7.01E-84 |
| Dmel_CG42565 | 636.741 | -4.964 | 0.276 | -17.965 | 3.68E-72 | 1.31E-68 |
| Dmel_CR34601 | 1403.059 | 8.261 | 0.467 | 17.692 | 4.87E-70 | 1.30E-66 |
| Dmel_CG32640 | 167.387 | -7.771 | 0.501 | -15.517 | 2.67E-54 | 5.72E-51 |
| Dmel_CG7408 | 343.022 | 5.184 | 0.347 | 14.940 | 1.80E-50 | 3.21E-47 |
| Dmel_CG15601 | 142.969 | 4.618 | 0.317 | 14.557 | 5.24E-48 | 8.01E-45 |
| Dmel_CG33801 | 977.281 | -4.316 | 0.304 | -14.175 | 1.32E-45 | 1.76E-42 |
| Dmel_CR45330 | 212.370 | -3.991 | 0.289 | -13.796 | 2.70E-43 | 3.21E-40 |
| Dmel_CG11825 | 247.162 | -3.656 | 0.277 | -13.206 | 8.11E-40 | 8.67E-37 |
| Dmel_CG33816 | 11696.267 | -9.588 | 0.748 | -12.825 | 1.18E-37 | 1.15E-34 |
| Dmel_CG32975 | 49.803 | 4.811 | 0.385 | 12.492 | 8.24E-36 | 7.34E-33 |
| Dmel_CG7052 | 166.253 | -3.532 | 0.284 | -12.447 | 1.45E-35 | 1.19E-32 |
| Dmel_CG31997 | 498.060 | -4.245 | 0.349 | -12.147 | 5.97E-34 | 4.56E-31 |
| Dmel_CG32474 | 120.332 | -7.181 | 0.592 | -12.135 | 6.88E-34 | 4.90E-31 |
| Dmel_CR43257 | 129.071 | -3.145 | 0.261 | -12.032 | 2.40E-33 | 1.60E-30 |
| Dmel_CG42584 | 108.987 | 4.408 | 0.374 | 11.777 | 5.12E-32 | 3.22E-29 |
| Dmel_CG8357 | 611.808 | -2.670 | 0.229 | -11.662 | 2.00E-31 | 1.19E-28 |
| Dmel_CG32475 | 179.753 | -10.041 | 0.863 | -11.639 | 2.62E-31 | 1.47E-28 |
| Dmel_CG2947 | 1490.588 | 1.955 | 0.171 | 11.448 | 2.42E-30 | 1.29E-27 |
| Dmel_CG4125 | 840.725 | 3.748 | 0.333 | 11.237 | 2.67E-29 | 1.36E-26 |
| Dmel_CR45284 | 114.416 | 9.866 | 0.890 | 11.083 | 1.51E-28 | 7.36E-26 |
| Dmel_CR46123 | 510.503 | -3.518 | 0.323 | -10.877 | 1.48E-27 | 6.87E-25 |
| Dmel_CG31683 | 923.718 | 7.381 | 0.681 | 10.838 | 2.28E-27 | 1.02E-24 |
| Dmel_CR45631 | 72.040 | -5.945 | 0.554 | -10.723 | 7.91E-27 | 3.38E-24 |
| Dmel_CG32600 | 181.352 | 3.164 | 0.296 | 10.680 | 1.26E-26 | 5.17E-24 |
| Dmel_CG10013 | 137.097 | -3.508 | 0.335 | -10.471 | 1.17E-25 | 4.65E-23 |
| Dmel_CG14233 | 178.085 | -2.132 | 0.206 | -10.352 | 4.09E-25 | 1.56E-22 |

| | | | | | | |
|---|---|---|---|---|---|---|
| Dmel_CG6864 | 56.941 | 5.134 | 0.497 | 10.322 | 5.61E-25 | 2.07E-22 |
| Dmel_CG18321 | 353.613 | -2.824 | 0.274 | -10.309 | 6.39E-25 | 2.28E-22 |
| Dmel_CG17684 | 1222.833 | 2.837 | 0.277 | 10.237 | 1.36E-24 | 4.69E-22 |
| Dmel_CG13941 | 124.885 | -2.456 | 0.246 | -10.003 | 1.47E-23 | 4.92E-21 |
| Dmel_CG8376 | 116.256 | -3.033 | 0.304 | -9.985 | 1.77E-23 | 5.75E-21 |
| Dmel_CR45567 | 40.179 | 8.757 | 0.888 | 9.863 | 6.00E-23 | 1.89E-20 |
| Dmel_CG9453 | 343.636 | -2.445 | 0.248 | -9.859 | 6.28E-23 | 1.92E-20 |
| Dmel_CG32647 | 110.757 | -3.051 | 0.313 | -9.739 | 2.05E-22 | 6.09E-20 |
| Dmel_CG34380 | 118.947 | 4.239 | 0.436 | 9.712 | 2.69E-22 | 7.77E-20 |
| Dmel_CG18188 | 45.646 | -3.579 | 0.369 | -9.703 | 2.93E-22 | 8.25E-20 |
| Dmel_CG13138 | 209.177 | 4.323 | 0.449 | 9.625 | 6.27E-22 | 1.72E-19 |
| Dmel_CG18853 | 75.915 | -8.396 | 0.873 | -9.621 | 6.49E-22 | 1.74E-19 |
| Dmel_CG33852 | 7295.263 | 4.955 | 0.520 | 9.520 | 1.74E-21 | 4.53E-19 |
| Dmel_CG1842 | 581.392 | 2.211 | 0.238 | 9.302 | 1.38E-20 | 3.51E-18 |
| Dmel_CG42255 | 42.513 | -3.971 | 0.428 | -9.277 | 1.74E-20 | 4.24E-18 |
| Dmel_CG42699 | 188.299 | 1.987 | 0.214 | 9.278 | 1.72E-20 | 4.24E-18 |
| Dmel_CG6269 | 43.176 | -3.049 | 0.330 | -9.228 | 2.76E-20 | 6.56E-18 |
| Dmel_CG13937 | 148.226 | 4.624 | 0.502 | 9.215 | 3.10E-20 | 7.21E-18 |
| Dmel_CG12112 | 1047.365 | -2.129 | 0.232 | -9.196 | 3.72E-20 | 8.46E-18 |
| Dmel_CG17470 | 57.133 | 4.145 | 0.461 | 8.986 | 2.57E-19 | 5.73E-17 |
| Dmel_CG5559 | 36.773 | 5.491 | 0.616 | 8.919 | 4.69E-19 | 1.02E-16 |
| Dmel_CG15312 | 1267.851 | 1.384 | 0.156 | 8.881 | 6.65E-19 | 1.42E-16 |
| Dmel_CG31792 | 38.167 | -4.478 | 0.506 | -8.851 | 8.70E-19 | 1.82E-16 |
| Dmel_CG9871 | 50.560 | -6.238 | 0.716 | -8.715 | 2.90E-18 | 5.97E-16 |
| Dmel_CG4373 | 307.928 | -4.950 | 0.573 | -8.646 | 5.35E-18 | 1.08E-15 |
| Dmel_CG3616 | 38.041 | -3.219 | 0.374 | -8.597 | 8.15E-18 | 1.61E-15 |
| Dmel_CG12592 | 187.503 | -3.278 | 0.382 | -8.580 | 9.47E-18 | 1.84E-15 |
| Dmel_CG12505 | 3126.415 | -2.939 | 0.344 | -8.542 | 1.32E-17 | 2.52E-15 |
| Dmel_CG5744 | 82.115 | -4.651 | 0.545 | -8.534 | 1.42E-17 | 2.66E-15 |
| Dmel_CG3578 | 148.306 | -1.836 | 0.217 | -8.471 | 2.43E-17 | 4.40E-15 |
| Dmel_CG9170 | 356.887 | -2.386 | 0.282 | -8.472 | 2.41E-17 | 4.40E-15 |

| | | | | | | |
|---|---|---|---|---|---|---|
| Dmel_CG9168 | 50.760 | 3.256 | 0.386 | 8.433 | 3.38E-17 | 6.03E-15 |
| Dmel_CR34602 | 25.163 | 8.166 | 0.970 | 8.417 | 3.85E-17 | 6.75E-15 |
| Dmel_CG12414 | 1841.851 | 1.885 | 0.225 | 8.376 | 5.48E-17 | 9.46E-15 |
| Dmel_CG32823 | 73.681 | 3.720 | 0.452 | 8.232 | 1.85E-16 | 3.13E-14 |
| Dmel_CR46499 | 109.382 | -2.030 | 0.248 | -8.197 | 2.46E-16 | 4.11E-14 |
| Dmel_CG31776 | 24.796 | -8.050 | 0.984 | -8.184 | 2.74E-16 | 4.51E-14 |
| Dmel_CG12520 | 87.565 | -2.304 | 0.285 | -8.082 | 6.36E-16 | 1.03E-13 |
| Dmel_CG14855 | 58.734 | 2.874 | 0.356 | 8.081 | 6.44E-16 | 1.03E-13 |
| Dmel_CG13579 | 22.440 | -8.336 | 1.034 | -8.059 | 7.69E-16 | 1.21E-13 |
| Dmel_CG12275 | 39.817 | -5.503 | 0.693 | -7.944 | 1.96E-15 | 3.03E-13 |
| Dmel_CG8821 | 106.295 | 3.716 | 0.473 | 7.864 | 3.71E-15 | 5.67E-13 |
| Dmel_CG16755 | 68.579 | 2.852 | 0.364 | 7.839 | 4.54E-15 | 6.85E-13 |
| Dmel_CR44264 | 17.002 | 7.192 | 0.922 | 7.803 | 6.05E-15 | 8.99E-13 |
| Dmel_CG2893 | 2410.997 | 1.218 | 0.156 | 7.793 | 6.53E-15 | 9.56E-13 |
| Dmel_CG7002 | 154.856 | -3.554 | 0.457 | -7.781 | 7.19E-15 | 1.04E-12 |
| Dmel_CG32789 | 795.569 | -1.531 | 0.197 | -7.778 | 7.35E-15 | 1.05E-12 |
| Dmel_CG31617 | 8715.429 | 2.006 | 0.258 | 7.769 | 7.90E-15 | 1.10E-12 |
| Dmel_CG33813 | 8715.429 | 2.006 | 0.258 | 7.769 | 7.90E-15 | 1.10E-12 |
| Dmel_CG16957 | 23.805 | 5.408 | 0.697 | 7.759 | 8.59E-15 | 1.18E-12 |
| Dmel_CG7900 | 157.605 | 3.411 | 0.441 | 7.744 | 9.66E-15 | 1.31E-12 |
| Dmel_CG14444 | 6588.810 | -0.958 | 0.124 | -7.713 | 1.23E-14 | 1.64E-12 |
| Dmel_CG6658 | 157.093 | -3.339 | 0.433 | -7.707 | 1.29E-14 | 1.70E-12 |
| Dmel_CG33868 | 2586.412 | -2.187 | 0.284 | -7.690 | 1.47E-14 | 1.92E-12 |
| Dmel_CG15706 | 424.122 | 1.599 | 0.209 | 7.659 | 1.88E-14 | 2.42E-12 |
| Dmel_CR43960 | 157.786 | -2.537 | 0.334 | -7.598 | 3.00E-14 | 3.82E-12 |
| Dmel_CR46481 | 1090.588 | -1.945 | 0.257 | -7.572 | 3.68E-14 | 4.63E-12 |
| Dmel_CR45323 | 25.859 | -6.703 | 0.889 | -7.541 | 4.68E-14 | 5.81E-12 |
| Dmel_CG12846 | 131.084 | -2.264 | 0.302 | -7.490 | 6.87E-14 | 8.44E-12 |
| Dmel_CG6890 | 2345.639 | 1.977 | 0.264 | 7.476 | 7.65E-14 | 9.30E-12 |
| Dmel_CG18405 | 1312.377 | 1.367 | 0.183 | 7.474 | 7.77E-14 | 9.33E-12 |
| Dmel_CG1232 | 80.890 | -3.916 | 0.524 | -7.467 | 8.21E-14 | 9.76E-12 |

| | | | | | | |
|---|---|---|---|---|---|---|
| Dmel_CG31865 | 120.039 | -2.432 | 0.327 | -7.436 | 1.04E-13 | 1.22E-11 |
| Dmel_CG32364 | 249.152 | -2.558 | 0.344 | -7.434 | 1.06E-13 | 1.23E-11 |
| Dmel_CG11144 | 634.856 | -1.765 | 0.239 | -7.392 | 1.45E-13 | 1.67E-11 |
| Dmel_CG1449 | 290.556 | -2.023 | 0.275 | -7.345 | 2.05E-13 | 2.33E-11 |
| Dmel_CG7665 | 49.711 | -2.544 | 0.352 | -7.224 | 5.05E-13 | 5.68E-11 |
| Dmel_CG6293 | 2110.091 | 2.582 | 0.358 | 7.217 | 5.33E-13 | 5.94E-11 |
| Dmel_CG14204 | 32.050 | 2.302 | 0.321 | 7.173 | 7.33E-13 | 8.08E-11 |
| Dmel_CG11354 | 35.882 | -2.429 | 0.341 | -7.125 | 1.04E-12 | 1.14E-10 |
| Dmel_CG14253 | 132.269 | 2.126 | 0.299 | 7.123 | 1.06E-12 | 1.14E-10 |
| Dmel_CG11099 | 22.666 | -5.708 | 0.802 | -7.116 | 1.11E-12 | 1.19E-10 |
| Dmel_CG33837 | 2738.319 | -2.233 | 0.314 | -7.108 | 1.18E-12 | 1.19E-10 |
| Dmel_CG33840 | 2738.319 | -2.233 | 0.314 | -7.108 | 1.18E-12 | 1.19E-10 |
| Dmel_CG33843 | 2738.319 | -2.233 | 0.314 | -7.108 | 1.18E-12 | 1.19E-10 |
| Dmel_CG33846 | 2738.319 | -2.233 | 0.314 | -7.108 | 1.18E-12 | 1.19E-10 |
| Dmel_CG33849 | 2738.319 | -2.233 | 0.314 | -7.108 | 1.18E-12 | 1.19E-10 |
| Dmel_CG33864 | 2738.319 | -2.233 | 0.314 | -7.108 | 1.18E-12 | 1.19E-10 |
| Dmel_CG33655 | 51.654 | 3.304 | 0.467 | 7.074 | 1.50E-12 | 1.50E-10 |
| Dmel_CG32490 | 168.640 | -2.039 | 0.288 | -7.073 | 1.52E-12 | 1.51E-10 |
| Dmel_CR45446 | 23.752 | 5.109 | 0.724 | 7.056 | 1.71E-12 | 1.68E-10 |
| Dmel_CG4587 | 91.859 | -2.577 | 0.366 | -7.044 | 1.87E-12 | 1.82E-10 |
| Dmel_CG8034 | 387.147 | -1.379 | 0.196 | -7.030 | 2.06E-12 | 1.99E-10 |
| Dmel_CG9981 | 21.353 | -6.048 | 0.861 | -7.027 | 2.11E-12 | 2.02E-10 |
| Dmel_CG14102 | 550.744 | 1.420 | 0.203 | 7.003 | 2.50E-12 | 2.37E-10 |
| Dmel_CG6424 | 1178.291 | -1.063 | 0.152 | -6.977 | 3.01E-12 | 2.82E-10 |
| Dmel_CG12548 | 110.057 | 2.956 | 0.424 | 6.971 | 3.14E-12 | 2.92E-10 |
| Dmel_CG31279 | 99.274 | -2.625 | 0.377 | -6.961 | 3.39E-12 | 3.13E-10 |
| Dmel_CG15632 | 24.357 | -6.024 | 0.867 | -6.950 | 3.65E-12 | 3.33E-10 |
| Dmel_CG14808 | 573.015 | -2.613 | 0.376 | -6.941 | 3.89E-12 | 3.52E-10 |
| Dmel_CG14518 | 13.705 | 6.644 | 0.962 | 6.905 | 5.02E-12 | 4.48E-10 |
| Dmel_CR46350 | 65.211 | 2.397 | 0.347 | 6.905 | 5.00E-12 | 4.48E-10 |
| Dmel_CR32881 | 15.023 | 6.709 | 0.976 | 6.877 | 6.13E-12 | 5.42E-10 |

| | | | | | | |
|---|---|---|---|---|---|---|
| Dmel_CG5106 | 180.963 | -2.039 | 0.298 | -6.852 | 7.29E-12 | 6.39E-10 |
| Dmel_CG4573 | 407.261 | 0.986 | 0.144 | 6.838 | 8.02E-12 | 6.97E-10 |
| Dmel_CG15673 | 374.682 | 2.079 | 0.305 | 6.808 | 9.91E-12 | 8.55E-10 |
| Dmel_CG32261 | 8.393 | 6.641 | 0.976 | 6.801 | 1.04E-11 | 8.87E-10 |
| Dmel_CG32540 | 27.797 | -5.848 | 0.863 | -6.776 | 1.23E-11 | 1.05E-09 |
| Dmel_CG40470 | 117.571 | 2.521 | 0.374 | 6.743 | 1.55E-11 | 1.30E-09 |
| Dmel_CG10654 | 255.238 | -1.959 | 0.291 | -6.723 | 1.78E-11 | 1.48E-09 |
| Dmel_CG32319 | 43.380 | 2.630 | 0.396 | 6.640 | 3.15E-11 | 2.61E-09 |
| Dmel_CG13876 | 343.928 | -1.331 | 0.202 | -6.595 | 4.25E-11 | 3.47E-09 |
| Dmel_CG17142 | 69.912 | -1.977 | 0.300 | -6.595 | 4.25E-11 | 3.47E-09 |
| Dmel_CR44603 | 26.607 | -6.925 | 1.050 | -6.592 | 4.33E-11 | 3.51E-09 |
| Dmel_CG13848 | 572.438 | -1.490 | 0.226 | -6.590 | 4.40E-11 | 3.53E-09 |
| Dmel_CG4116 | 39.770 | 2.830 | 0.430 | 6.578 | 4.75E-11 | 3.79E-09 |
| Dmel_CG8321 | 886.301 | -1.178 | 0.181 | -6.515 | 7.27E-11 | 5.76E-09 |
| Dmel_CG2397 | 133.002 | -1.613 | 0.248 | -6.499 | 8.08E-11 | 6.36E-09 |
| Dmel_CR44441 | 70.571 | 2.873 | 0.444 | 6.472 | 9.70E-11 | 7.57E-09 |
| Dmel_CG4666 | 110.138 | -2.471 | 0.383 | -6.451 | 1.11E-10 | 8.62E-09 |
| Dmel_CG32944 | 92.791 | -2.040 | 0.317 | -6.441 | 1.19E-10 | 9.12E-09 |
| Dmel_CR44649 | 14.040 | 5.807 | 0.902 | 6.440 | 1.19E-10 | 9.12E-09 |
| Dmel_CG3592 | 24.833 | 3.723 | 0.579 | 6.435 | 1.24E-10 | 9.39E-09 |
| Dmel_CG15267 | 102.975 | 2.512 | 0.391 | 6.432 | 1.26E-10 | 9.51E-09 |
| Dmel_CG4181 | 124.651 | -2.626 | 0.408 | -6.430 | 1.28E-10 | 9.54E-09 |
| Dmel_CG33861 | 849.240 | -2.803 | 0.438 | -6.394 | 1.61E-10 | 1.20E-08 |
| Dmel_CG5130 | 868.620 | -1.343 | 0.212 | -6.343 | 2.26E-10 | 1.67E-08 |
| Dmel_CG42343 | 28.745 | -2.528 | 0.399 | -6.331 | 2.43E-10 | 1.78E-08 |
| Dmel_CG12405 | 52.075 | -1.899 | 0.300 | -6.320 | 2.62E-10 | 1.90E-08 |
| Dmel_CR34573 | 15.764 | 5.560 | 0.880 | 6.317 | 2.66E-10 | 1.92E-08 |
| Dmel_CG14162 | 1349.763 | 1.374 | 0.219 | 6.270 | 3.61E-10 | 2.58E-08 |
| Dmel_CG4500 | 7.493 | 6.350 | 1.013 | 6.270 | 3.60E-10 | 2.58E-08 |
| Dmel_CG32204 | 31.301 | -2.860 | 0.456 | -6.268 | 3.66E-10 | 2.59E-08 |
| Dmel_CG11372 | 1894.097 | 1.758 | 0.281 | 6.264 | 3.75E-10 | 2.62E-08 |

| | | | | | | |
|---|---|---|---|---|---|---|
| Dmel_CG3134 | 31.621 | -3.101 | 0.495 | -6.265 | 3.73E-10 | 2.62E-08 |
| Dmel_CG11160 | 16.059 | -5.132 | 0.820 | -6.261 | 3.82E-10 | 2.66E-08 |
| Dmel_CG3653 | 339.374 | 2.166 | 0.346 | 6.258 | 3.89E-10 | 2.68E-08 |
| Dmel_CG3832 | 660.172 | 1.414 | 0.226 | 6.254 | 4.00E-10 | 2.74E-08 |
| Dmel_CR31853 | 15.583 | -6.169 | 0.987 | -6.248 | 4.17E-10 | 2.84E-08 |
| Dmel_CG46315 | 1065.167 | -0.931 | 0.149 | -6.227 | 4.75E-10 | 3.22E-08 |
| Dmel_CG10062 | 88.652 | -2.498 | 0.402 | -6.217 | 5.06E-10 | 3.41E-08 |
| Dmel_CG45019 | 491.689 | -1.639 | 0.264 | -6.213 | 5.19E-10 | 3.47E-08 |
| Dmel_CG18495 | 1721.586 | -1.927 | 0.311 | -6.193 | 5.92E-10 | 3.93E-08 |
| Dmel_CG14416 | 20.114 | 3.533 | 0.571 | 6.187 | 6.12E-10 | 4.04E-08 |
| Dmel_CG30401 | 80.682 | -2.540 | 0.412 | -6.167 | 6.96E-10 | 4.56E-08 |
| Dmel_CR44285 | 16.737 | -4.535 | 0.738 | -6.149 | 7.81E-10 | 5.09E-08 |
| Dmel_CR44842 | 51.141 | 2.339 | 0.381 | 6.139 | 8.31E-10 | 5.39E-08 |
| Dmel_CG6644 | 396.732 | -2.312 | 0.377 | -6.137 | 8.42E-10 | 5.41E-08 |
| Dmel_CR43957 | 57.161 | -2.135 | 0.348 | -6.136 | 8.44E-10 | 5.41E-08 |
| Dmel_CG30026 | 144.278 | 2.596 | 0.423 | 6.135 | 8.50E-10 | 5.41E-08 |
| Dmel_CG11263 | 812.988 | 1.589 | 0.260 | 6.120 | 9.33E-10 | 5.91E-08 |
| Dmel_CG17959 | 18.151 | 3.189 | 0.522 | 6.114 | 9.69E-10 | 6.10E-08 |
| Dmel_CG32191 | 18.062 | 3.536 | 0.579 | 6.104 | 1.03E-09 | 6.45E-08 |
| Dmel_CG15695 | 15.117 | -5.448 | 0.899 | -6.058 | 1.38E-09 | 8.58E-08 |
| Dmel_CG8964 | 30.624 | -2.560 | 0.424 | -6.041 | 1.53E-09 | 9.48E-08 |
| Dmel_CR46147 | 58.757 | -1.968 | 0.326 | -6.035 | 1.59E-09 | 9.77E-08 |
| Dmel_CG13426 | 9.974 | 6.113 | 1.016 | 6.018 | 1.77E-09 | 1.08E-07 |
| Dmel_CG6352 | 20.044 | -3.623 | 0.602 | -6.017 | 1.78E-09 | 1.08E-07 |
| Dmel_CG4486 | 56.498 | 2.700 | 0.449 | 6.015 | 1.80E-09 | 1.09E-07 |
| Dmel_CG6829 | 1109.723 | 1.216 | 0.204 | 5.975 | 2.30E-09 | 1.38E-07 |
| Dmel_CG18766 | 417.316 | 1.249 | 0.209 | 5.962 | 2.49E-09 | 1.49E-07 |
| Dmel_CG9475 | 65.197 | -1.919 | 0.322 | -5.959 | 2.53E-09 | 1.51E-07 |
| Dmel_CG4381 | 408.162 | -1.619 | 0.273 | -5.941 | 2.84E-09 | 1.68E-07 |
| Dmel_CG7863 | 307.615 | 1.102 | 0.186 | 5.935 | 2.94E-09 | 1.73E-07 |
| Dmel_CG13540 | 16.816 | -4.164 | 0.703 | -5.924 | 3.14E-09 | 1.84E-07 |

| | | | | | | |
|---|---|---|---|---|---|---|
| Dmel_CG11940 | 4570.884 | 1.039 | 0.176 | 5.892 | 3.81E-09 | 2.21E-07 |
| Dmel_CG45781 | 99.356 | 2.496 | 0.424 | 5.880 | 4.10E-09 | 2.37E-07 |
| Dmel_CG1544 | 56.811 | 3.186 | 0.543 | 5.869 | 4.38E-09 | 2.52E-07 |
| Dmel_CR41257 | 19.191 | 3.731 | 0.637 | 5.859 | 4.67E-09 | 2.67E-07 |
| Dmel_CG13375 | 172.764 | -1.742 | 0.298 | -5.852 | 4.85E-09 | 2.76E-07 |
| Dmel_CG7084 | 38.686 | -3.722 | 0.638 | -5.838 | 5.29E-09 | 2.99E-07 |
| Dmel_CG12370 | 119.708 | -2.318 | 0.398 | -5.825 | 5.71E-09 | 3.20E-07 |
| Dmel_CR43635 | 28.969 | -2.640 | 0.453 | -5.826 | 5.68E-09 | 3.20E-07 |
| Dmel_CG3036 | 780.679 | -1.187 | 0.204 | -5.821 | 5.86E-09 | 3.27E-07 |
| Dmel_CG13617 | 40.995 | 2.463 | 0.425 | 5.801 | 6.60E-09 | 3.66E-07 |
| Dmel_CG42639 | 16.055 | -4.825 | 0.832 | -5.796 | 6.77E-09 | 3.73E-07 |
| Dmel_CR44648 | 7.481 | 5.743 | 0.999 | 5.747 | 9.08E-09 | 4.98E-07 |
| Dmel_CG32187 | 11.172 | 6.232 | 1.085 | 5.745 | 9.19E-09 | 5.02E-07 |
| Dmel_CG10650 | 150.114 | -1.962 | 0.342 | -5.741 | 9.43E-09 | 5.12E-07 |
| Dmel_CG17836 | 1929.733 | -1.570 | 0.274 | -5.732 | 9.94E-09 | 5.37E-07 |
| Dmel_CR43605 | 66.687 | 2.061 | 0.360 | 5.719 | 1.07E-08 | 5.75E-07 |
| Dmel_CG3588 | 32.672 | 2.561 | 0.448 | 5.716 | 1.09E-08 | 5.83E-07 |
| Dmel_CG7433 | 4321.842 | -0.989 | 0.174 | -5.689 | 1.27E-08 | 6.78E-07 |
| Dmel_CG9701 | 11.604 | -4.237 | 0.747 | -5.676 | 1.38E-08 | 7.27E-07 |
| Dmel_CR32886 | 5281.077 | -1.355 | 0.239 | -5.676 | 1.38E-08 | 7.27E-07 |
| Dmel_CG44193 | 1265.153 | -1.120 | 0.198 | -5.661 | 1.50E-08 | 7.84E-07 |
| Dmel_CG9411 | 11.974 | -4.948 | 0.874 | -5.661 | 1.50E-08 | 7.84E-07 |
| Dmel_CG12986 | 9.202 | -5.950 | 1.054 | -5.645 | 1.66E-08 | 8.60E-07 |
| Dmel_CG42566 | 568.495 | -2.099 | 0.372 | -5.643 | 1.67E-08 | 8.62E-07 |
| Dmel_CG18789 | 575.054 | -1.270 | 0.226 | -5.613 | 1.99E-08 | 1.02E-06 |
| Dmel_CR42861 | 395.796 | -1.322 | 0.236 | -5.607 | 2.06E-08 | 1.06E-06 |
| Dmel_CG10246 | 507.936 | -1.118 | 0.200 | -5.587 | 2.31E-08 | 1.17E-06 |
| Dmel_CG2381 | 70.702 | 2.064 | 0.369 | 5.587 | 2.31E-08 | 1.17E-06 |
| Dmel_CG8165 | 279.924 | 0.802 | 0.144 | 5.583 | 2.37E-08 | 1.19E-06 |
| Dmel_CG42694 | 322.776 | 1.171 | 0.210 | 5.571 | 2.53E-08 | 1.27E-06 |
| Dmel_CR43836 | 1370.913 | 1.125 | 0.202 | 5.567 | 2.58E-08 | 1.29E-06 |

| | | | | | | |
|---|---|---|---|---|---|---|
| Dmel_CG11280 | 82.750 | 1.787 | 0.322 | 5.545 | 2.94E-08 | 1.46E-06 |
| Dmel_CG30046 | 65.427 | 1.783 | 0.322 | 5.539 | 3.05E-08 | 1.51E-06 |
| Dmel_CR34151 | 2597.926 | 1.281 | 0.232 | 5.528 | 3.24E-08 | 1.60E-06 |
| Dmel_CR44105 | 111.289 | 1.812 | 0.328 | 5.522 | 3.35E-08 | 1.65E-06 |
| Dmel_CG3926 | 50.891 | -2.071 | 0.375 | -5.518 | 3.43E-08 | 1.68E-06 |
| Dmel_CG15422 | 16.358 | -3.229 | 0.589 | -5.478 | 4.30E-08 | 2.08E-06 |
| Dmel_CG7970 | 1818.632 | -0.761 | 0.139 | -5.478 | 4.31E-08 | 2.08E-06 |
| Dmel_CR45600 | 123.977 | -1.441 | 0.264 | -5.466 | 4.62E-08 | 2.22E-06 |
| Dmel_CR42871 | 50.583 | -2.562 | 0.471 | -5.439 | 5.36E-08 | 2.57E-06 |
| Dmel_CG10962 | 178.967 | 2.521 | 0.464 | 5.433 | 5.55E-08 | 2.65E-06 |
| Dmel_CR44370 | 668.992 | 1.879 | 0.348 | 5.400 | 6.65E-08 | 3.16E-06 |
| Dmel_CG14591 | 15.962 | -3.474 | 0.643 | -5.399 | 6.71E-08 | 3.17E-06 |
| Dmel_CG45002 | 81.687 | 2.435 | 0.451 | 5.397 | 6.79E-08 | 3.20E-06 |
| Dmel_CR43887 | 48.400 | -2.054 | 0.381 | -5.391 | 6.99E-08 | 3.28E-06 |
| Dmel_CG8193 | 9.669 | -5.178 | 0.963 | -5.378 | 7.53E-08 | 3.52E-06 |
| Dmel_CG10693 | 63.837 | -2.304 | 0.429 | -5.370 | 7.89E-08 | 3.67E-06 |
| Dmel_CG17657 | 833.649 | 0.932 | 0.174 | 5.368 | 7.97E-08 | 3.69E-06 |
| Dmel_CG9922 | 1514.484 | -1.367 | 0.255 | -5.357 | 8.45E-08 | 3.89E-06 |
| Dmel_CG4998 | 35.482 | -2.912 | 0.545 | -5.344 | 9.07E-08 | 4.16E-06 |
| Dmel_CG14615 | 327.742 | 1.131 | 0.212 | 5.326 | 1.00E-07 | 4.58E-06 |
| Dmel_CG17669 | 10.096 | -4.037 | 0.759 | -5.317 | 1.06E-07 | 4.80E-06 |
| Dmel_CG16779 | 55.503 | -2.362 | 0.445 | -5.311 | 1.09E-07 | 4.93E-06 |
| Dmel_CG3548 | 367.754 | 1.676 | 0.319 | 5.258 | 1.46E-07 | 6.58E-06 |
| Dmel_CG1629 | 52.870 | -1.812 | 0.345 | -5.255 | 1.48E-07 | 6.66E-06 |
| Dmel_CG8666 | 1194.885 | -1.041 | 0.198 | -5.247 | 1.55E-07 | 6.94E-06 |
| Dmel_CG13780 | 102.811 | -3.674 | 0.701 | -5.241 | 1.60E-07 | 7.11E-06 |
| Dmel_CG30048 | 8.917 | -5.534 | 1.060 | -5.222 | 1.77E-07 | 7.84E-06 |
| Dmel_CG45544 | 25.811 | -2.041 | 0.391 | -5.219 | 1.80E-07 | 7.97E-06 |
| Dmel_CG10390 | 88.576 | -1.834 | 0.352 | -5.206 | 1.93E-07 | 8.49E-06 |
| Dmel_CR43461 | 15.553 | -2.544 | 0.489 | -5.205 | 1.94E-07 | 8.51E-06 |
| Dmel_CG12535 | 350.714 | -1.351 | 0.260 | -5.192 | 2.08E-07 | 9.08E-06 |

| | | | | | | |
|---|---|---|---|---|---|---|
| Dmel_CG14026 | 3581.522 | 0.517 | 0.100 | 5.176 | 2.26E-07 | 9.83E-06 |
| Dmel_CR43334 | 902.502 | 1.433 | 0.278 | 5.161 | 2.46E-07 | 1.07E-05 |
| Dmel_CG31956 | 7.679 | -5.030 | 0.975 | -5.159 | 2.48E-07 | 1.07E-05 |
| Dmel_CG8256 | 121.600 | -1.630 | 0.317 | -5.150 | 2.61E-07 | 1.12E-05 |
| Dmel_CG14424 | 11.673 | 4.159 | 0.809 | 5.138 | 2.77E-07 | 1.18E-05 |
| Dmel_CR45714 | 39.803 | 2.530 | 0.492 | 5.138 | 2.78E-07 | 1.18E-05 |
| Dmel_CG5927 | 130.550 | 3.518 | 0.685 | 5.134 | 2.83E-07 | 1.20E-05 |
| Dmel_CG42711 | 9.934 | -5.047 | 0.984 | -5.130 | 2.90E-07 | 1.23E-05 |
| Dmel_CG5022 | 23.972 | 2.232 | 0.436 | 5.117 | 3.10E-07 | 1.31E-05 |
| Dmel_CG10559 | 18.084 | -2.548 | 0.498 | -5.112 | 3.18E-07 | 1.33E-05 |
| Dmel_CG42362 | 214.695 | -1.187 | 0.232 | -5.107 | 3.28E-07 | 1.37E-05 |
| Dmel_CG42363 | 214.695 | -1.187 | 0.232 | -5.107 | 3.28E-07 | 1.37E-05 |
| Dmel_CG10245 | 245.057 | -1.401 | 0.275 | -5.096 | 3.47E-07 | 1.44E-05 |
| Dmel_CG12582 | 1823.184 | 0.794 | 0.156 | 5.095 | 3.48E-07 | 1.44E-05 |
| Dmel_CG3346 | 377.229 | 1.460 | 0.287 | 5.090 | 3.59E-07 | 1.48E-05 |
| Dmel_CG34313 | 123.277 | -1.216 | 0.239 | -5.082 | 3.73E-07 | 1.53E-05 |
| Dmel_CR46488 | 96.924 | 2.110 | 0.416 | 5.077 | 3.83E-07 | 1.56E-05 |
| Dmel_CR43283 | 44.621 | 4.945 | 0.975 | 5.074 | 3.90E-07 | 1.58E-05 |
| Dmel_CG15705 | 4.557 | 5.773 | 1.138 | 5.072 | 3.94E-07 | 1.60E-05 |
| Dmel_CG43798 | 14.265 | -2.669 | 0.527 | -5.068 | 4.02E-07 | 1.62E-05 |
| Dmel_CG1851 | 132.622 | -1.408 | 0.278 | -5.063 | 4.12E-07 | 1.66E-05 |
| Dmel_CG34329 | 9.708 | -5.152 | 1.018 | -5.060 | 4.20E-07 | 1.68E-05 |
| Dmel_CG11186 | 154.593 | 2.035 | 0.403 | 5.044 | 4.55E-07 | 1.82E-05 |
| Dmel_CG1631 | 6.321 | -5.205 | 1.033 | -5.039 | 4.69E-07 | 1.86E-05 |
| Dmel_CG2671 | 15581.812 | 1.147 | 0.228 | 5.039 | 4.67E-07 | 1.86E-05 |
| Dmel_CG10247 | 12.494 | -3.256 | 0.647 | -5.034 | 4.81E-07 | 1.90E-05 |
| Dmel_CG44102 | 63.130 | 2.047 | 0.407 | 5.024 | 5.07E-07 | 1.99E-05 |
| Dmel_CG44328 | 42.014 | -2.179 | 0.436 | -4.999 | 5.75E-07 | 2.25E-05 |
| Dmel_CG3546 | 24.816 | 1.961 | 0.392 | 4.996 | 5.84E-07 | 2.28E-05 |
| Dmel_CG3767 | 118.430 | -1.757 | 0.352 | -4.996 | 5.86E-07 | 2.28E-05 |
| Dmel_CG10391 | 295.990 | 1.311 | 0.263 | 4.985 | 6.19E-07 | 2.40E-05 |

| | | | | | | |
|---|---|---|---|---|---|---|
| Dmel_CG32595 | 13.102 | -3.220 | 0.647 | -4.979 | 6.38E-07 | 2.46E-05 |
| Dmel_CG31174 | 20.644 | -1.919 | 0.386 | -4.970 | 6.71E-07 | 2.58E-05 |
| Dmel_CG3598 | 12.213 | 4.080 | 0.823 | 4.958 | 7.10E-07 | 2.72E-05 |
| Dmel_CR34621 | 154.656 | 1.896 | 0.383 | 4.956 | 7.19E-07 | 2.75E-05 |
| Dmel_CG10160 | 13.374 | -3.030 | 0.613 | -4.944 | 7.64E-07 | 2.90E-05 |
| Dmel_CG10287 | 102.780 | -1.638 | 0.331 | -4.945 | 7.62E-07 | 2.90E-05 |
| Dmel_CG11450 | 84.360 | 1.490 | 0.302 | 4.942 | 7.75E-07 | 2.93E-05 |
| Dmel_CG31901 | 28.374 | -3.474 | 0.703 | -4.939 | 7.84E-07 | 2.95E-05 |
| Dmel_CG8051 | 10.915 | -2.609 | 0.529 | -4.930 | 8.24E-07 | 3.09E-05 |
| Dmel_CG45058 | 655.515 | 1.169 | 0.237 | 4.929 | 8.27E-07 | 3.09E-05 |
| Dmel_CG31693 | 14.293 | -3.563 | 0.724 | -4.921 | 8.63E-07 | 3.22E-05 |
| Dmel_CG3022 | 27.256 | -2.367 | 0.482 | -4.916 | 8.85E-07 | 3.29E-05 |
| Dmel_CG30383 | 49.644 | -1.689 | 0.344 | -4.913 | 8.98E-07 | 3.32E-05 |
| Dmel_CG13916 | 111.331 | 2.194 | 0.447 | 4.905 | 9.33E-07 | 3.44E-05 |
| Dmel_CG11205 | 397.986 | -1.798 | 0.368 | -4.892 | 1.00E-06 | 3.67E-05 |
| Dmel_CG32814 | 302.950 | -1.768 | 0.362 | -4.887 | 1.02E-06 | 3.75E-05 |
| Dmel_CG10695 | 3201.672 | -1.191 | 0.244 | -4.886 | 1.03E-06 | 3.76E-05 |
| Dmel_CG12763 | 27.517 | -6.787 | 1.391 | -4.878 | 1.07E-06 | 3.89E-05 |
| Dmel_CR46284 | 5.583 | -5.554 | 1.139 | -4.878 | 1.07E-06 | 3.89E-05 |
| Dmel_CG14173 | 20.836 | 2.596 | 0.533 | 4.872 | 1.11E-06 | 3.99E-05 |
| Dmel_CG15083 | 836.327 | -1.186 | 0.244 | -4.861 | 1.17E-06 | 4.21E-05 |
| Dmel_CG6502 | 1593.297 | 0.700 | 0.144 | 4.856 | 1.20E-06 | 4.31E-05 |
| Dmel_CG14502 | 161.282 | 1.211 | 0.250 | 4.843 | 1.28E-06 | 4.57E-05 |
| Dmel_CG11125 | 345.793 | 1.079 | 0.224 | 4.819 | 1.44E-06 | 5.14E-05 |
| Dmel_CG17523 | 9.395 | -2.831 | 0.588 | -4.816 | 1.46E-06 | 5.19E-05 |
| Dmel_CG31718 | 13.955 | -2.733 | 0.568 | -4.815 | 1.47E-06 | 5.21E-05 |
| Dmel_CG13401 | 706.768 | 0.814 | 0.169 | 4.813 | 1.49E-06 | 5.24E-05 |
| Dmel_CG8942 | 20.884 | -3.571 | 0.742 | -4.813 | 1.49E-06 | 5.24E-05 |
| Dmel_CG34002 | 25.029 | -2.108 | 0.439 | -4.807 | 1.53E-06 | 5.38E-05 |
| Dmel_CG34251 | 4.631 | -5.622 | 1.173 | -4.793 | 1.64E-06 | 5.74E-05 |
| Dmel_CG12473 | 121.417 | 1.006 | 0.210 | 4.788 | 1.69E-06 | 5.86E-05 |

| | | | | | | |
|---|---|---|---|---|---|---|
| Dmel_CG12500 | 121.417 | 1.006 | 0.210 | 4.788 | 1.69E-06 | 5.86E-05 |
| Dmel_CG30101 | 46.664 | 2.599 | 0.543 | 4.783 | 1.73E-06 | 5.99E-05 |
| Dmel_CG10794 | 23.361 | -6.086 | 1.273 | -4.782 | 1.74E-06 | 5.99E-05 |
| Dmel_CG32845 | 27.360 | -2.062 | 0.431 | -4.780 | 1.75E-06 | 6.02E-05 |
| Dmel_CG18095 | 35.302 | 4.346 | 0.910 | 4.775 | 1.80E-06 | 6.12E-05 |
| Dmel_CG6798 | 39.400 | 2.450 | 0.513 | 4.774 | 1.80E-06 | 6.12E-05 |
| Dmel_CG8561 | 48.662 | -2.040 | 0.427 | -4.774 | 1.80E-06 | 6.12E-05 |
| Dmel_CR45215 | 6.526 | 3.592 | 0.752 | 4.774 | 1.80E-06 | 6.12E-05 |
| Dmel_CG13091 | 80.925 | 2.136 | 0.447 | 4.773 | 1.81E-06 | 6.14E-05 |
| Dmel_CG31619 | 134.708 | -1.970 | 0.413 | -4.773 | 1.82E-06 | 6.14E-05 |
| Dmel_CG17604 | 2294.672 | 1.027 | 0.215 | 4.769 | 1.85E-06 | 6.23E-05 |
| Dmel_CG10089 | 69.275 | 1.575 | 0.331 | 4.758 | 1.96E-06 | 6.56E-05 |
| Dmel_CG12730 | 69.324 | 1.436 | 0.303 | 4.742 | 2.11E-06 | 7.06E-05 |
| Dmel_CG17885 | 29.019 | -1.638 | 0.346 | -4.739 | 2.15E-06 | 7.15E-05 |
| Dmel_CG1343 | 8.774 | -4.947 | 1.046 | -4.730 | 2.24E-06 | 7.42E-05 |
| Dmel_CG14420 | 15.867 | 2.736 | 0.578 | 4.731 | 2.24E-06 | 7.42E-05 |
| Dmel_CG14489 | 4874.516 | 0.630 | 0.134 | 4.717 | 2.39E-06 | 7.89E-05 |
| Dmel_CG4476 | 2247.740 | 1.223 | 0.259 | 4.714 | 2.43E-06 | 7.98E-05 |
| Dmel_CG5976 | 1222.502 | -1.007 | 0.214 | -4.712 | 2.46E-06 | 8.06E-05 |
| Dmel_CR34645 | 9.943 | 4.238 | 0.900 | 4.708 | 2.51E-06 | 8.20E-05 |
| Dmel_CG33531 | 10.591 | 3.263 | 0.696 | 4.688 | 2.76E-06 | 9.02E-05 |
| Dmel_CG2505 | 44.932 | 3.869 | 0.826 | 4.683 | 2.83E-06 | 9.21E-05 |
| Dmel_CG9921 | 462.165 | -1.269 | 0.271 | -4.676 | 2.92E-06 | 9.46E-05 |
| Dmel_CG4780 | 256.282 | -1.154 | 0.247 | -4.675 | 2.94E-06 | 9.48E-05 |
| Dmel_CG32485 | 670.585 | 0.877 | 0.188 | 4.674 | 2.96E-06 | 9.51E-05 |
| Dmel_CG33494 | 73.278 | -2.457 | 0.526 | -4.674 | 2.96E-06 | 9.51E-05 |
| Dmel_CG14584 | 13.563 | -2.618 | 0.561 | -4.667 | 3.06E-06 | 9.80E-05 |
| Dmel_CR46075 | 89.606 | 1.330 | 0.286 | 4.660 | 3.17E-06 | 1.01E-04 |
| Dmel_CG4563 | 4.531 | -5.693 | 1.222 | -4.658 | 3.19E-06 | 1.02E-04 |
| Dmel_CG11700 | 93.380 | -1.222 | 0.263 | -4.655 | 3.24E-06 | 1.03E-04 |
| Dmel_CR32773 | 602.990 | 1.163 | 0.251 | 4.639 | 3.50E-06 | 1.11E-04 |

| | | | | | | |
|---|---|---|---|---|---|---|
| Dmel_CG42338 | 533.899 | -1.121 | 0.242 | -4.637 | 3.53E-06 | 1.11E-04 |
| Dmel_CG1483 | 10242.605 | 1.798 | 0.389 | 4.626 | 3.73E-06 | 1.17E-04 |
| Dmel_CG43867 | 1609.935 | 1.111 | 0.241 | 4.615 | 3.93E-06 | 1.23E-04 |
| Dmel_CR46231 | 7.957 | -5.437 | 1.179 | -4.613 | 3.97E-06 | 1.24E-04 |
| Dmel_CG32459 | 8.115 | -3.873 | 0.840 | -4.610 | 4.02E-06 | 1.25E-04 |
| Dmel_CG17610 | 1299.127 | 0.816 | 0.177 | 4.604 | 4.14E-06 | 1.29E-04 |
| Dmel_CG10816 | 140.617 | -3.652 | 0.794 | -4.601 | 4.21E-06 | 1.30E-04 |
| Dmel_CG1464 | 109.018 | 1.495 | 0.325 | 4.601 | 4.20E-06 | 1.30E-04 |
| Dmel_CR44833 | 413.121 | -0.980 | 0.213 | -4.601 | 4.21E-06 | 1.30E-04 |
| Dmel_CG1922 | 81.028 | 2.023 | 0.440 | 4.596 | 4.32E-06 | 1.33E-04 |
| Dmel_CG14356 | 31.409 | 2.251 | 0.490 | 4.594 | 4.36E-06 | 1.33E-04 |
| Dmel_CG6986 | 148.456 | -1.258 | 0.274 | -4.594 | 4.35E-06 | 1.33E-04 |
| Dmel_CG13636 | 2287.981 | 3.194 | 0.696 | 4.587 | 4.50E-06 | 1.37E-04 |
| Dmel_CG8279 | 98.613 | 1.543 | 0.336 | 4.587 | 4.49E-06 | 1.37E-04 |
| Dmel_CR34558 | 120.892 | 1.745 | 0.381 | 4.575 | 4.77E-06 | 1.45E-04 |
| Dmel_CG2657 | 34.331 | -2.251 | 0.492 | -4.573 | 4.81E-06 | 1.45E-04 |
| Dmel_CG46313 | 39.798 | 4.693 | 1.027 | 4.568 | 4.93E-06 | 1.48E-04 |
| Dmel_CG30382 | 3253.403 | -1.858 | 0.408 | -4.552 | 5.31E-06 | 1.59E-04 |
| Dmel_CG14686 | 5.867 | -5.206 | 1.144 | -4.550 | 5.35E-06 | 1.60E-04 |
| Dmel_CG18568 | 26.606 | 1.474 | 0.324 | 4.549 | 5.39E-06 | 1.61E-04 |
| Dmel_CG8023 | 8.117 | -4.055 | 0.891 | -4.549 | 5.39E-06 | 1.61E-04 |
| Dmel_CG15347 | 570.453 | -1.578 | 0.348 | -4.531 | 5.88E-06 | 1.75E-04 |
| Dmel_CG14948 | 111.422 | -1.570 | 0.347 | -4.521 | 6.17E-06 | 1.83E-04 |
| Dmel_CG8825 | 1165.276 | -1.305 | 0.289 | -4.511 | 6.45E-06 | 1.91E-04 |
| Dmel_CG3397 | 15.052 | -2.800 | 0.621 | -4.507 | 6.58E-06 | 1.94E-04 |
| Dmel_CG11094 | 251.870 | -1.553 | 0.345 | -4.502 | 6.73E-06 | 1.98E-04 |
| Dmel_CG9169 | 13.677 | 2.700 | 0.600 | 4.502 | 6.74E-06 | 1.98E-04 |
| Dmel_CG11390 | 112.637 | -2.243 | 0.498 | -4.500 | 6.78E-06 | 1.98E-04 |
| Dmel_CG43326 | 10.654 | -2.837 | 0.631 | -4.497 | 6.89E-06 | 2.01E-04 |
| Dmel_CG10151 | 33.822 | -2.032 | 0.452 | -4.492 | 7.05E-06 | 2.05E-04 |
| Dmel_CG9093 | 881.441 | -0.757 | 0.169 | -4.491 | 7.07E-06 | 2.05E-04 |

| | | | | | | |
|---|---|---|---|---|---|---|
| Dmel_CG2297 | 30.219 | -2.497 | 0.556 | -4.487 | 7.24E-06 | 2.09E-04 |
| Dmel_CG14931 | 1536.983 | 1.612 | 0.359 | 4.485 | 7.29E-06 | 2.10E-04 |
| Dmel_CG2528 | 8.511 | -4.082 | 0.910 | -4.485 | 7.30E-06 | 2.10E-04 |
| Dmel_CG1092 | 40.559 | -2.093 | 0.467 | -4.481 | 7.44E-06 | 2.13E-04 |
| Dmel_CG14688 | 934.841 | -0.685 | 0.153 | -4.476 | 7.60E-06 | 2.17E-04 |
| Dmel_CG15820 | 1082.505 | 0.912 | 0.204 | 4.477 | 7.59E-06 | 2.17E-04 |
| Dmel_CG43088 | 13.320 | -3.076 | 0.689 | -4.464 | 8.04E-06 | 2.29E-04 |
| Dmel_CG4950 | 13.824 | -3.830 | 0.858 | -4.461 | 8.16E-06 | 2.31E-04 |
| Dmel_CG3323 | 48.498 | 1.884 | 0.423 | 4.457 | 8.31E-06 | 2.35E-04 |
| Dmel_CG33481 | 45.632 | -1.894 | 0.425 | -4.455 | 8.39E-06 | 2.36E-04 |
| Dmel_CG8145 | 489.756 | -0.916 | 0.206 | -4.455 | 8.39E-06 | 2.36E-04 |
| Dmel_CR43264 | 294.406 | -1.474 | 0.331 | -4.453 | 8.45E-06 | 2.37E-04 |
| Dmel_CG11052 | 8.299 | 4.274 | 0.962 | 4.442 | 8.91E-06 | 2.50E-04 |
| Dmel_CG1960 | 4602.892 | 0.621 | 0.140 | 4.441 | 8.96E-06 | 2.50E-04 |
| Dmel_CG7727 | 134.263 | 1.172 | 0.265 | 4.429 | 9.48E-06 | 2.64E-04 |
| Dmel_CG32165 | 1153.177 | -1.572 | 0.355 | -4.427 | 9.54E-06 | 2.65E-04 |
| Dmel_CG9742 | 1047.738 | -1.639 | 0.371 | -4.423 | 9.75E-06 | 2.70E-04 |
| Dmel_CG43366 | 29.454 | -2.231 | 0.505 | -4.420 | 9.89E-06 | 2.73E-04 |
| Dmel_CG12470 | 17.654 | -2.391 | 0.542 | -4.408 | 1.04E-05 | 2.87E-04 |
| Dmel_CG6128 | 36.042 | -2.440 | 0.554 | -4.403 | 1.07E-05 | 2.94E-04 |
| Dmel_CG6698 | 17.300 | -3.573 | 0.812 | -4.401 | 1.08E-05 | 2.96E-04 |
| Dmel_CG1443 | 162.341 | 1.837 | 0.418 | 4.393 | 1.12E-05 | 3.05E-04 |
| Dmel_CG41520 | 200.878 | 1.460 | 0.332 | 4.392 | 1.12E-05 | 3.07E-04 |
| Dmel_CG18437 | 12.750 | -2.891 | 0.660 | -4.380 | 1.19E-05 | 3.24E-04 |
| Dmel_CR34575 | 10.820 | 2.843 | 0.650 | 4.377 | 1.20E-05 | 3.27E-04 |
| Dmel_CG7391 | 327.485 | 1.327 | 0.304 | 4.365 | 1.27E-05 | 3.44E-04 |
| Dmel_CG5080 | 19.507 | -2.449 | 0.562 | -4.355 | 1.33E-05 | 3.60E-04 |
| Dmel_CG14907 | 262.745 | -1.083 | 0.249 | -4.348 | 1.37E-05 | 3.70E-04 |
| Dmel_CG10638 | 1166.946 | -0.953 | 0.219 | -4.343 | 1.40E-05 | 3.77E-04 |
| Dmel_CG42829 | 4.622 | -5.282 | 1.218 | -4.337 | 1.45E-05 | 3.88E-04 |
| Dmel_CG7918 | 20.384 | 2.296 | 0.530 | 4.334 | 1.46E-05 | 3.91E-04 |

| | | | | | | |
|---|---|---|---|---|---|---|
| Dmel_CR32914 | 35.179 | -1.441 | 0.333 | -4.331 | 1.49E-05 | 3.97E-04 |
| Dmel_CG42368 | 30.697 | 1.790 | 0.414 | 4.319 | 1.57E-05 | 4.17E-04 |
| Dmel_CG10936 | 307.998 | 1.283 | 0.297 | 4.313 | 1.61E-05 | 4.28E-04 |
| Dmel_CG14423 | 8.974 | 3.751 | 0.872 | 4.299 | 1.71E-05 | 4.54E-04 |
| Dmel_CR43989 | 55.193 | -1.304 | 0.304 | -4.297 | 1.73E-05 | 4.57E-04 |
| Dmel_CR44183 | 4.078 | 4.873 | 1.134 | 4.296 | 1.74E-05 | 4.58E-04 |
| Dmel_CG6044 | 14.793 | 2.702 | 0.630 | 4.289 | 1.79E-05 | 4.71E-04 |
| Dmel_CG2812 | 61.353 | -1.769 | 0.413 | -4.287 | 1.81E-05 | 4.75E-04 |
| Dmel_CG1506 | 66.867 | -1.574 | 0.368 | -4.273 | 1.93E-05 | 5.04E-04 |
| Dmel_CG4620 | 4335.134 | 0.799 | 0.187 | 4.268 | 1.97E-05 | 5.14E-04 |
| Dmel_CG7549 | 52.479 | -1.357 | 0.318 | -4.265 | 2.00E-05 | 5.20E-04 |
| Dmel_CG31519 | 27.150 | -2.340 | 0.550 | -4.252 | 2.12E-05 | 5.50E-04 |
| Dmel_CG42281 | 10048.584 | 0.592 | 0.139 | 4.251 | 2.13E-05 | 5.50E-04 |
| Dmel_CR43302 | 6.039 | 3.219 | 0.758 | 4.248 | 2.16E-05 | 5.57E-04 |
| Dmel_CG17077 | 839.671 | 0.783 | 0.186 | 4.204 | 2.63E-05 | 6.77E-04 |
| Dmel_CG11961 | 335.734 | -0.664 | 0.158 | -4.201 | 2.66E-05 | 6.84E-04 |
| Dmel_CG1969 | 470.276 | -0.499 | 0.119 | -4.186 | 2.83E-05 | 7.27E-04 |
| Dmel_CG5399 | 18.394 | -2.327 | 0.557 | -4.180 | 2.92E-05 | 7.47E-04 |
| Dmel_CG1873 | 30.188 | -2.071 | 0.496 | -4.176 | 2.97E-05 | 7.57E-04 |
| Dmel_CG32017 | 173.619 | 1.439 | 0.345 | 4.174 | 3.00E-05 | 7.63E-04 |
| Dmel_CG12749 | 4132.742 | -0.630 | 0.151 | -4.169 | 3.07E-05 | 7.79E-04 |
| Dmel_CG7152 | 41.498 | 1.793 | 0.430 | 4.168 | 3.08E-05 | 7.80E-04 |
| Dmel_CG31601 | 12.167 | -3.060 | 0.735 | -4.163 | 3.13E-05 | 7.92E-04 |
| Dmel_CR18854 | 838.629 | 0.990 | 0.238 | 4.163 | 3.14E-05 | 7.92E-04 |
| Dmel_CG9580 | 35.037 | -1.839 | 0.442 | -4.162 | 3.15E-05 | 7.93E-04 |
| Dmel_CG31618 | 2229.642 | -0.953 | 0.229 | -4.154 | 3.26E-05 | 8.07E-04 |
| Dmel_CG33814 | 2229.642 | -0.953 | 0.229 | -4.154 | 3.26E-05 | 8.07E-04 |
| Dmel_CG33817 | 2229.642 | -0.953 | 0.229 | -4.154 | 3.26E-05 | 8.07E-04 |
| Dmel_CG33820 | 2229.642 | -0.953 | 0.229 | -4.154 | 3.26E-05 | 8.07E-04 |
| Dmel_CG33823 | 2229.642 | -0.953 | 0.229 | -4.154 | 3.26E-05 | 8.07E-04 |
| Dmel_CG33826 | 2229.642 | -0.953 | 0.229 | -4.154 | 3.26E-05 | 8.07E-04 |

| | | | | | | |
|---|---|---|---|---|---|---|
| Dmel_CG33829 | 2229.642 | -0.953 | 0.229 | -4.154 | 3.26E-05 | 8.07E-04 |
| Dmel_CG5792 | 3509.804 | 0.710 | 0.171 | 4.152 | 3.29E-05 | 8.12E-04 |
| Dmel_CR45923 | 329.714 | 1.048 | 0.252 | 4.152 | 3.30E-05 | 8.12E-04 |
| Dmel_CG11714 | 5.974 | -3.867 | 0.934 | -4.141 | 3.46E-05 | 8.50E-04 |
| Dmel_CG8638 | 7.909 | -2.815 | 0.680 | -4.139 | 3.49E-05 | 8.56E-04 |
| Dmel_CR43609 | 35.121 | -2.292 | 0.554 | -4.138 | 3.51E-05 | 8.58E-04 |
| Dmel_CG10723 | 53.062 | -1.426 | 0.345 | -4.134 | 3.57E-05 | 8.69E-04 |
| Dmel_CG5322 | 43.751 | -2.396 | 0.580 | -4.134 | 3.56E-05 | 8.69E-04 |
| Dmel_CG9380 | 914.006 | 2.463 | 0.596 | 4.130 | 3.62E-05 | 8.80E-04 |
| Dmel_CG34360 | 2556.488 | 0.835 | 0.202 | 4.129 | 3.64E-05 | 8.84E-04 |
| Dmel_CR44366 | 13.310 | -1.815 | 0.440 | -4.122 | 3.76E-05 | 9.11E-04 |
| Dmel_CG7395 | 11.002 | 2.961 | 0.719 | 4.120 | 3.79E-05 | 9.16E-04 |
| Dmel_CR43898 | 11.698 | 2.122 | 0.515 | 4.116 | 3.85E-05 | 9.27E-04 |
| Dmel_CG1004 | 121.543 | 1.633 | 0.397 | 4.112 | 3.93E-05 | 9.44E-04 |
| Dmel_CG31288 | 22.525 | -2.495 | 0.609 | -4.095 | 4.21E-05 | 1.01E-03 |
| Dmel_CG10877 | 88.197 | -1.683 | 0.411 | -4.093 | 4.26E-05 | 1.02E-03 |
| Dmel_CR32896 | 10.122 | -3.330 | 0.814 | -4.093 | 4.26E-05 | 1.02E-03 |
| Dmel_CG17681 | 14.570 | -2.303 | 0.563 | -4.090 | 4.31E-05 | 1.03E-03 |
| Dmel_CG5338 | 70.237 | -2.034 | 0.498 | -4.086 | 4.39E-05 | 1.04E-03 |
| Dmel_CG15143 | 100.498 | 1.533 | 0.375 | 4.083 | 4.44E-05 | 1.05E-03 |
| Dmel_CG6067 | 10.429 | -3.312 | 0.811 | -4.084 | 4.43E-05 | 1.05E-03 |
| Dmel_CG7882 | 16.945 | -4.855 | 1.190 | -4.080 | 4.51E-05 | 1.07E-03 |
| Dmel_CG14640 | 9.690 | -2.996 | 0.735 | -4.075 | 4.60E-05 | 1.08E-03 |
| Dmel_CG30170 | 20.719 | -2.277 | 0.560 | -4.065 | 4.80E-05 | 1.13E-03 |
| Dmel_CG14606 | 111.001 | 4.228 | 1.042 | 4.059 | 4.92E-05 | 1.15E-03 |
| Dmel_CG42486 | 7.794 | -2.105 | 0.520 | -4.053 | 5.06E-05 | 1.18E-03 |
| Dmel_CR44914 | 34.591 | 2.238 | 0.552 | 4.051 | 5.09E-05 | 1.19E-03 |
| Dmel_CG15599 | 55.457 | -2.241 | 0.553 | -4.050 | 5.13E-05 | 1.20E-03 |
| Dmel_CG17572 | 7.733 | 2.997 | 0.743 | 4.032 | 5.52E-05 | 1.28E-03 |
| Dmel_CG33779 | 7.876 | 2.716 | 0.674 | 4.032 | 5.54E-05 | 1.28E-03 |
| Dmel_CG9907 | 59.516 | -1.325 | 0.329 | -4.031 | 5.56E-05 | 1.29E-03 |

| | | | | | | |
|---|---|---|---|---|---|---|
| Dmel_CG32313 | 7.303 | 3.037 | 0.754 | 4.029 | 5.60E-05 | 1.29E-03 |
| Dmel_CG3008 | 1125.630 | -0.785 | 0.195 | -4.025 | 5.69E-05 | 1.31E-03 |
| Dmel_CG32082 | 143.362 | -1.201 | 0.299 | -4.012 | 6.03E-05 | 1.38E-03 |
| Dmel_CG32381 | 6.858 | -3.414 | 0.851 | -4.012 | 6.02E-05 | 1.38E-03 |
| Dmel_CR45199 | 15.610 | 2.262 | 0.564 | 4.011 | 6.05E-05 | 1.38E-03 |
| Dmel_CR45580 | 8.928 | -1.996 | 0.499 | -4.003 | 6.26E-05 | 1.43E-03 |
| Dmel_CG3857 | 518.727 | 0.838 | 0.210 | 3.997 | 6.43E-05 | 1.47E-03 |
| Dmel_CG15034 | 80.952 | 1.369 | 0.343 | 3.991 | 6.59E-05 | 1.50E-03 |
| Dmel_CG1683 | 61.082 | -1.424 | 0.357 | -3.988 | 6.67E-05 | 1.51E-03 |
| Dmel_CG15598 | 8.615 | 2.751 | 0.690 | 3.985 | 6.75E-05 | 1.53E-03 |
| Dmel_CG43389 | 9.534 | -2.635 | 0.664 | -3.969 | 7.21E-05 | 1.63E-03 |
| Dmel_CG7149 | 511.711 | 0.872 | 0.220 | 3.958 | 7.55E-05 | 1.70E-03 |
| Dmel_CG7383 | 5.086 | -4.596 | 1.162 | -3.956 | 7.62E-05 | 1.72E-03 |
| Dmel_CG12477 | 8.616 | -4.050 | 1.027 | -3.944 | 8.01E-05 | 1.80E-03 |
| Dmel_CG6125 | 90.955 | -1.593 | 0.404 | -3.944 | 8.03E-05 | 1.80E-03 |
| Dmel_CR45029 | 87.357 | 1.013 | 0.257 | 3.944 | 8.02E-05 | 1.80E-03 |
| Dmel_CG32365 | 2522.157 | 0.876 | 0.222 | 3.939 | 8.17E-05 | 1.82E-03 |
| Dmel_CG43395 | 5.634 | -3.479 | 0.884 | -3.937 | 8.24E-05 | 1.84E-03 |
| Dmel_CG32793 | 17.470 | 2.676 | 0.680 | 3.933 | 8.38E-05 | 1.86E-03 |
| Dmel_CG9738 | 2394.475 | -0.619 | 0.157 | -3.933 | 8.39E-05 | 1.86E-03 |
| Dmel_CG17570 | 6.795 | 2.553 | 0.650 | 3.931 | 8.47E-05 | 1.87E-03 |
| Dmel_CG31262 | 1828.361 | 0.790 | 0.201 | 3.930 | 8.49E-05 | 1.88E-03 |
| Dmel_CG11155 | 1048.739 | -1.266 | 0.323 | -3.917 | 8.97E-05 | 1.98E-03 |
| Dmel_CG34431 | 161.743 | -1.020 | 0.261 | -3.910 | 9.22E-05 | 2.03E-03 |
| Dmel_CG34031 | 23.601 | -2.008 | 0.514 | -3.905 | 9.43E-05 | 2.07E-03 |
| Dmel_CG9780 | 15.302 | -2.148 | 0.550 | -3.902 | 9.54E-05 | 2.09E-03 |
| Dmel_CR46268 | 170.251 | -1.026 | 0.263 | -3.901 | 9.58E-05 | 2.10E-03 |
| Dmel_CG14643 | 29.737 | -2.214 | 0.568 | -3.897 | 9.75E-05 | 2.13E-03 |
| Dmel_CG7542 | 10.141 | 2.826 | 0.726 | 3.892 | 9.92E-05 | 2.16E-03 |
| Dmel_CG42598 | 119.404 | 3.509 | 0.902 | 3.891 | 1.00E-04 | 2.17E-03 |
| Dmel_CG14616 | 5403.289 | 0.544 | 0.140 | 3.890 | 1.00E-04 | 2.17E-03 |

| | | | | | | |
|---|---|---|---|---|---|---|
| Dmel_CG17907 | 42.096 | -1.423 | 0.366 | -3.887 | 1.01E-04 | 2.19E-03 |
| Dmel_CG3448 | 358.908 | -0.949 | 0.244 | -3.888 | 1.01E-04 | 2.19E-03 |
| Dmel_CG42335 | 49.366 | 4.755 | 1.224 | 3.884 | 1.03E-04 | 2.21E-03 |
| Dmel_CG15150 | 6.838 | -3.981 | 1.025 | -3.884 | 1.03E-04 | 2.21E-03 |
| Dmel_CR42254 | 371.532 | 0.984 | 0.254 | 3.881 | 1.04E-04 | 2.24E-03 |
| Dmel_CG10091 | 216.140 | -1.617 | 0.417 | -3.876 | 1.06E-04 | 2.27E-03 |
| Dmel_CG3796 | 20.569 | -1.603 | 0.414 | -3.873 | 1.08E-04 | 2.30E-03 |
| Dmel_CR45334 | 5.283 | -3.658 | 0.946 | -3.869 | 1.09E-04 | 2.34E-03 |
| Dmel_CR42910 | 65.924 | 1.696 | 0.439 | 3.866 | 1.11E-04 | 2.36E-03 |
| Dmel_CG14639 | 7.664 | -3.024 | 0.783 | -3.863 | 1.12E-04 | 2.37E-03 |
| Dmel_CR42743 | 4.749 | -4.446 | 1.151 | -3.863 | 1.12E-04 | 2.37E-03 |
| Dmel_CR44348 | 10.906 | 2.269 | 0.588 | 3.862 | 1.13E-04 | 2.39E-03 |
| Dmel_CG33658 | 6.283 | 4.239 | 1.100 | 3.855 | 1.16E-04 | 2.45E-03 |
| Dmel_CG5392 | 15.491 | -1.899 | 0.493 | -3.850 | 1.18E-04 | 2.49E-03 |
| Dmel_CR44953 | 22.110 | -1.841 | 0.479 | -3.845 | 1.21E-04 | 2.54E-03 |
| Dmel_CG33111 | 598.892 | 0.612 | 0.159 | 3.841 | 1.23E-04 | 2.58E-03 |
| Dmel_CG14933 | 32.287 | -2.019 | 0.526 | -3.839 | 1.24E-04 | 2.59E-03 |
| Dmel_CG31801 | 8.478 | -2.242 | 0.585 | -3.833 | 1.27E-04 | 2.65E-03 |
| Dmel_CR45124 | 51.575 | 1.671 | 0.436 | 3.832 | 1.27E-04 | 2.65E-03 |
| Dmel_CG12910 | 6.836 | 3.245 | 0.847 | 3.832 | 1.27E-04 | 2.65E-03 |
| Dmel_CG31015 | 538.599 | 1.076 | 0.281 | 3.831 | 1.28E-04 | 2.66E-03 |
| Dmel_CG1743 | 114.738 | -1.366 | 0.357 | -3.827 | 1.29E-04 | 2.69E-03 |
| Dmel_CG8003 | 1339.184 | 0.620 | 0.162 | 3.825 | 1.31E-04 | 2.71E-03 |
| Dmel_CG6701 | 6459.136 | 0.625 | 0.163 | 3.823 | 1.32E-04 | 2.72E-03 |
| Dmel_CG4608 | 113.986 | -1.184 | 0.310 | -3.821 | 1.33E-04 | 2.75E-03 |
| Dmel_CG12410 | 544.961 | 0.593 | 0.155 | 3.817 | 1.35E-04 | 2.77E-03 |
| Dmel_CG44835 | 2748.099 | 0.677 | 0.177 | 3.818 | 1.35E-04 | 2.77E-03 |
| Dmel_CG6324 | 34.299 | -1.676 | 0.439 | -3.818 | 1.34E-04 | 2.77E-03 |
| Dmel_CG9472 | 10.918 | 2.066 | 0.542 | 3.812 | 1.38E-04 | 2.82E-03 |
| Dmel_CG13793 | 53.229 | 1.552 | 0.408 | 3.808 | 1.40E-04 | 2.86E-03 |
| Dmel_CG14757 | 107.605 | 0.895 | 0.235 | 3.802 | 1.44E-04 | 2.93E-03 |

| | | | | | | |
|---|---|---|---|---|---|---|
| Dmel_CG18408 | 587.102 | 0.741 | 0.195 | 3.799 | 1.45E-04 | 2.94E-03 |
| Dmel_CG34357 | 51.034 | 1.339 | 0.352 | 3.800 | 1.45E-04 | 2.94E-03 |
| Dmel_CG5455 | 372.518 | -1.046 | 0.275 | -3.800 | 1.45E-04 | 2.94E-03 |
| Dmel_CG11356 | 13.659 | -2.148 | 0.566 | -3.794 | 1.48E-04 | 2.99E-03 |
| Dmel_CR45577 | 14.166 | 2.232 | 0.588 | 3.795 | 1.48E-04 | 2.99E-03 |
| Dmel_CG5644 | 28.092 | 1.858 | 0.491 | 3.787 | 1.52E-04 | 3.08E-03 |
| Dmel_CG42813 | 353.216 | 1.405 | 0.371 | 3.787 | 1.53E-04 | 3.08E-03 |
| Dmel_CR44230 | 13.634 | -2.024 | 0.535 | -3.784 | 1.54E-04 | 3.10E-03 |
| Dmel_CG16778 | 54.657 | 2.348 | 0.621 | 3.779 | 1.57E-04 | 3.15E-03 |
| Dmel_CG8588 | 597.074 | 1.012 | 0.268 | 3.774 | 1.61E-04 | 3.22E-03 |
| Dmel_CR43242 | 104.596 | 1.334 | 0.354 | 3.772 | 1.62E-04 | 3.23E-03 |
| Dmel_CR34626 | 1126.535 | 1.548 | 0.411 | 3.768 | 1.64E-04 | 3.28E-03 |
| Dmel_CG41434 | 7.270 | -3.616 | 0.961 | -3.762 | 1.68E-04 | 3.35E-03 |
| Dmel_CG8274 | 6007.375 | -0.557 | 0.148 | -3.762 | 1.69E-04 | 3.35E-03 |
| Dmel_CG3104 | 56.006 | -1.293 | 0.344 | -3.758 | 1.71E-04 | 3.40E-03 |
| Dmel_CG14062 | 3.626 | -4.929 | 1.313 | -3.754 | 1.74E-04 | 3.44E-03 |
| Dmel_CG42294 | 6.836 | -3.208 | 0.855 | -3.752 | 1.75E-04 | 3.47E-03 |
| Dmel_CG13604 | 682.846 | 0.784 | 0.209 | 3.749 | 1.78E-04 | 3.50E-03 |
| Dmel_CG13970 | 20.195 | -2.312 | 0.617 | -3.748 | 1.78E-04 | 3.51E-03 |
| Dmel_CG11049 | 56.984 | -1.611 | 0.430 | -3.743 | 1.82E-04 | 3.57E-03 |
| Dmel_CG1894 | 11.141 | -2.082 | 0.557 | -3.735 | 1.88E-04 | 3.68E-03 |
| Dmel_CG10469 | 24.417 | -1.430 | 0.383 | -3.730 | 1.92E-04 | 3.76E-03 |
| Dmel_CG4260 | 7777.907 | 0.530 | 0.142 | 3.727 | 1.93E-04 | 3.78E-03 |
| Dmel_CG10868 | 20270.917 | 0.732 | 0.197 | 3.726 | 1.95E-04 | 3.80E-03 |
| Dmel_CG9968 | 1175.027 | -0.709 | 0.191 | -3.710 | 2.07E-04 | 4.03E-03 |
| Dmel_CG12800 | 317.337 | -1.068 | 0.288 | -3.705 | 2.11E-04 | 4.11E-03 |
| Dmel_CG10776 | 280.046 | 0.841 | 0.228 | 3.687 | 2.26E-04 | 4.40E-03 |
| Dmel_CG31477 | 8.939 | -2.314 | 0.631 | -3.669 | 2.43E-04 | 4.71E-03 |
| Dmel_CG31721 | 43.300 | -1.874 | 0.511 | -3.669 | 2.43E-04 | 4.71E-03 |
| Dmel_CG30000 | 890.667 | -1.008 | 0.275 | -3.665 | 2.48E-04 | 4.78E-03 |
| Dmel_CG10440 | 6.298 | 2.592 | 0.708 | 3.662 | 2.50E-04 | 4.81E-03 |

| | | | | | | |
|---|---|---|---|---|---|---|
| Dmel_CG18522 | 301.389 | -1.437 | 0.392 | -3.662 | 2.50E-04 | 4.81E-03 |
| Dmel_CG4209 | 16.847 | -1.945 | 0.531 | -3.661 | 2.51E-04 | 4.82E-03 |
| Dmel_CG42316 | 192.031 | -1.525 | 0.417 | -3.658 | 2.54E-04 | 4.88E-03 |
| Dmel_CG13772 | 15.539 | -2.247 | 0.614 | -3.657 | 2.55E-04 | 4.88E-03 |
| Dmel_CG9339 | 7193.178 | 0.491 | 0.134 | 3.657 | 2.55E-04 | 4.88E-03 |
| Dmel_CG12662 | 86.071 | 1.509 | 0.413 | 3.656 | 2.56E-04 | 4.89E-03 |
| Dmel_CG3359 | 1017.402 | -0.873 | 0.239 | -3.654 | 2.58E-04 | 4.91E-03 |
| Dmel_CG2679 | 77.051 | 0.973 | 0.266 | 3.654 | 2.59E-04 | 4.91E-03 |
| Dmel_CG12190 | 1070.714 | 0.942 | 0.258 | 3.652 | 2.60E-04 | 4.92E-03 |
| Dmel_CG15879 | 82.055 | 0.980 | 0.268 | 3.652 | 2.60E-04 | 4.92E-03 |
| Dmel_CG17352 | 58.907 | 2.070 | 0.567 | 3.650 | 2.62E-04 | 4.96E-03 |
| Dmel_CG9165 | 1988.485 | -1.065 | 0.292 | -3.647 | 2.65E-04 | 5.01E-03 |
| Dmel_CR45510 | 7.744 | -3.035 | 0.833 | -3.645 | 2.67E-04 | 5.03E-03 |
| Dmel_CG31436 | 22.798 | -1.887 | 0.519 | -3.639 | 2.74E-04 | 5.14E-03 |
| Dmel_CG40305 | 50.491 | 1.110 | 0.305 | 3.639 | 2.74E-04 | 5.14E-03 |
| Dmel_CG2641 | 219.749 | -0.803 | 0.221 | -3.637 | 2.76E-04 | 5.17E-03 |
| Dmel_CG33474 | 26.760 | -1.370 | 0.377 | -3.634 | 2.79E-04 | 5.22E-03 |
| Dmel_CG34120 | 382.219 | -0.954 | 0.263 | -3.631 | 2.82E-04 | 5.26E-03 |
| Dmel_CG8606 | 869.507 | 0.588 | 0.162 | 3.624 | 2.90E-04 | 5.40E-03 |
| Dmel_CG6232 | 43.952 | -1.009 | 0.279 | -3.622 | 2.92E-04 | 5.43E-03 |
| Dmel_CG34411 | 5.697 | -3.139 | 0.868 | -3.617 | 2.98E-04 | 5.53E-03 |
| Dmel_CG2985 | 15846.085 | -1.395 | 0.386 | -3.613 | 3.03E-04 | 5.60E-03 |
| Dmel_CG33855 | 661.466 | 1.268 | 0.351 | 3.613 | 3.03E-04 | 5.60E-03 |
| Dmel_CG33858 | 661.466 | 1.268 | 0.351 | 3.613 | 3.03E-04 | 5.60E-03 |
| Dmel_CG18313 | 6.700 | 2.594 | 0.719 | 3.609 | 3.07E-04 | 5.67E-03 |
| Dmel_CG34325 | 218.050 | -1.043 | 0.289 | -3.608 | 3.08E-04 | 5.67E-03 |
| Dmel_CG43749 | 68.898 | -1.747 | 0.484 | -3.608 | 3.08E-04 | 5.67E-03 |
| Dmel_CR45822 | 59.722 | 1.856 | 0.515 | 3.606 | 3.11E-04 | 5.71E-03 |
| Dmel_CG31300 | 10.149 | -2.756 | 0.765 | -3.603 | 3.15E-04 | 5.77E-03 |
| Dmel_CG15828 | 114.025 | -2.543 | 0.706 | -3.601 | 3.17E-04 | 5.79E-03 |
| Dmel_CG7565 | 530.624 | 0.618 | 0.172 | 3.588 | 3.33E-04 | 6.08E-03 |

| | | | | | | |
|---|---|---|---|---|---|---|
| Dmel_CG13624 | 4837.368 | 0.626 | 0.175 | 3.587 | 3.35E-04 | 6.10E-03 |
| Dmel_CR45472 | 12.276 | 2.189 | 0.611 | 3.584 | 3.38E-04 | 6.15E-03 |
| Dmel_CG1532 | 1363.500 | -0.736 | 0.205 | -3.581 | 3.43E-04 | 6.22E-03 |
| Dmel_CG1971 | 113.016 | -0.924 | 0.258 | -3.580 | 3.44E-04 | 6.23E-03 |
| Dmel_CG31687 | 689.422 | -0.814 | 0.228 | -3.577 | 3.47E-04 | 6.28E-03 |
| Dmel_CG7737 | 406.711 | -0.897 | 0.251 | -3.576 | 3.49E-04 | 6.31E-03 |
| Dmel_CG3837 | 293.345 | 0.924 | 0.258 | 3.574 | 3.51E-04 | 6.34E-03 |
| Dmel_CG13659 | 14.223 | -1.963 | 0.550 | -3.566 | 3.62E-04 | 6.52E-03 |
| Dmel_CG8819 | 502.091 | -0.591 | 0.166 | -3.566 | 3.63E-04 | 6.52E-03 |
| Dmel_CG12099 | 2772.921 | -0.642 | 0.180 | -3.563 | 3.67E-04 | 6.57E-03 |
| Dmel_CG9068 | 9.058 | 3.563 | 1.000 | 3.563 | 3.67E-04 | 6.57E-03 |
| Dmel_CG10481 | 27.823 | -1.465 | 0.412 | -3.559 | 3.72E-04 | 6.66E-03 |
| Dmel_CG14307 | 1087.564 | 0.958 | 0.269 | 3.557 | 3.75E-04 | 6.70E-03 |
| Dmel_CG31918 | 958.032 | -0.742 | 0.209 | -3.551 | 3.83E-04 | 6.83E-03 |
| Dmel_CG32354 | 36.624 | -1.592 | 0.448 | -3.551 | 3.84E-04 | 6.83E-03 |
| Dmel_CR46216 | 14.209 | -2.633 | 0.741 | -3.551 | 3.84E-04 | 6.83E-03 |
| Dmel_CG14472 | 25993.883 | 0.315 | 0.089 | 3.549 | 3.87E-04 | 6.86E-03 |
| Dmel_CG7466 | 1646.183 | -0.502 | 0.142 | -3.539 | 4.02E-04 | 7.10E-03 |
| Dmel_CR43303 | 10.440 | 2.706 | 0.765 | 3.539 | 4.02E-04 | 7.10E-03 |
| Dmel_CG18102 | 3933.133 | 0.395 | 0.112 | 3.538 | 4.03E-04 | 7.11E-03 |
| Dmel_CG9652 | 17.219 | 1.772 | 0.501 | 3.537 | 4.05E-04 | 7.13E-03 |
| Dmel_CG12066 | 170.368 | -0.917 | 0.259 | -3.535 | 4.07E-04 | 7.16E-03 |
| Dmel_CG45057 | 104.242 | -1.329 | 0.376 | -3.532 | 4.12E-04 | 7.23E-03 |
| Dmel_CG9925 | 8017.743 | 0.506 | 0.143 | 3.530 | 4.15E-04 | 7.28E-03 |
| Dmel_CG2759 | 71.072 | -1.197 | 0.339 | -3.528 | 4.18E-04 | 7.33E-03 |
| Dmel_CG3159 | 17.200 | -1.981 | 0.562 | -3.527 | 4.21E-04 | 7.35E-03 |
| Dmel_CG40006 | 2376.258 | 0.767 | 0.217 | 3.526 | 4.21E-04 | 7.35E-03 |
| Dmel_CG3329 | 5252.860 | -0.707 | 0.201 | -3.522 | 4.28E-04 | 7.45E-03 |
| Dmel_CR45260 | 5.707 | 4.054 | 1.153 | 3.517 | 4.37E-04 | 7.60E-03 |
| Dmel_CR44330 | 9.131 | 2.344 | 0.667 | 3.516 | 4.38E-04 | 7.60E-03 |
| Dmel_CG15784 | 293.016 | -1.201 | 0.342 | -3.513 | 4.43E-04 | 7.67E-03 |

| | | | | | | |
|---|---|---|---|---|---|---|
| Dmel_CG9390 | 2169.006 | -0.876 | 0.249 | -3.513 | 4.43E-04 | 7.67E-03 |
| Dmel_CG43921 | 462.043 | 0.947 | 0.270 | 3.511 | 4.47E-04 | 7.72E-03 |
| Dmel_CG43052 | 15.089 | 1.909 | 0.544 | 3.508 | 4.52E-04 | 7.79E-03 |
| Dmel_CG17292 | 1474.083 | 0.696 | 0.199 | 3.503 | 4.59E-04 | 7.91E-03 |
| Dmel_CG13384 | 2664.424 | 0.482 | 0.138 | 3.500 | 4.65E-04 | 8.00E-03 |
| Dmel_CG3565 | 6.277 | -3.306 | 0.945 | -3.499 | 4.67E-04 | 8.01E-03 |
| Dmel_CG10833 | 21.891 | -4.331 | 1.239 | -3.496 | 4.72E-04 | 8.10E-03 |
| Dmel_CG13539 | 9.683 | 2.242 | 0.642 | 3.495 | 4.74E-04 | 8.12E-03 |
| Dmel_CG3879 | 12.492 | -2.007 | 0.575 | -3.491 | 4.81E-04 | 8.21E-03 |
| Dmel_CG13140 | 9.747 | -1.950 | 0.559 | -3.485 | 4.93E-04 | 8.37E-03 |
| Dmel_CG3209 | 1529.199 | 0.411 | 0.118 | 3.485 | 4.93E-04 | 8.37E-03 |
| Dmel_CG4472 | 34.227 | -1.331 | 0.382 | -3.485 | 4.93E-04 | 8.37E-03 |
| Dmel_CG7874 | 21.781 | -4.907 | 1.408 | -3.486 | 4.91E-04 | 8.37E-03 |
| Dmel_CG34141 | 119.156 | 0.971 | 0.279 | 3.477 | 5.08E-04 | 8.60E-03 |
| Dmel_CG31866 | 314.021 | 0.646 | 0.186 | 3.475 | 5.10E-04 | 8.63E-03 |
| Dmel_CR44793 | 9.708 | -2.928 | 0.843 | -3.474 | 5.12E-04 | 8.65E-03 |
| Dmel_CG7997 | 780.900 | 0.530 | 0.153 | 3.474 | 5.13E-04 | 8.65E-03 |
| Dmel_CR34635 | 8.396 | 2.244 | 0.646 | 3.473 | 5.15E-04 | 8.67E-03 |
| Dmel_CG16956 | 123.954 | 1.533 | 0.442 | 3.466 | 5.28E-04 | 8.87E-03 |
| Dmel_CG3929 | 5193.590 | -0.960 | 0.277 | -3.463 | 5.35E-04 | 8.98E-03 |
| Dmel_CG9901 | 6113.781 | 0.361 | 0.104 | 3.462 | 5.36E-04 | 8.98E-03 |
| Dmel_CG6618 | 40.165 | 1.385 | 0.401 | 3.456 | 5.48E-04 | 9.17E-03 |
| Dmel_CR43607 | 38.001 | 1.008 | 0.292 | 3.454 | 5.52E-04 | 9.22E-03 |
| Dmel_CG17527 | 10.179 | -2.272 | 0.658 | -3.453 | 5.55E-04 | 9.27E-03 |
| Dmel_CR44024 | 22.499 | 1.792 | 0.520 | 3.447 | 5.67E-04 | 9.45E-03 |
| Dmel_CG3091 | 43.997 | -1.541 | 0.447 | -3.445 | 5.70E-04 | 9.49E-03 |
| Dmel_CG7644 | 30.473 | -1.781 | 0.517 | -3.445 | 5.71E-04 | 9.49E-03 |
| Dmel_CR45789 | 14.109 | 1.924 | 0.560 | 3.438 | 5.86E-04 | 9.72E-03 |
| Dmel_CG30385 | 5.769 | -2.845 | 0.829 | -3.433 | 5.96E-04 | 9.84E-03 |
| Dmel_CG4995 | 6.334 | -2.402 | 0.700 | -3.433 | 5.97E-04 | 9.84E-03 |
| Dmel_CG6704 | 461.899 | 1.126 | 0.328 | 3.433 | 5.96E-04 | 9.84E-03 |

| | | | | | | |
|---|---|---|---|---|---|---|
| Dmel_CR43949 | 41.829 | 1.450 | 0.422 | 3.433 | 5.97E-04 | 9.84E-03 |
| Dmel_CG10241 | 394.726 | 1.066 | 0.311 | 3.430 | 6.03E-04 | 9.87E-03 |
| Dmel_CG10683 | 1174.214 | 0.730 | 0.213 | 3.431 | 6.02E-04 | 9.87E-03 |
| Dmel_CG13516 | 62.853 | -1.387 | 0.404 | -3.430 | 6.03E-04 | 9.87E-03 |
| Dmel_CG32350 | 8070.134 | 0.783 | 0.228 | 3.430 | 6.03E-04 | 9.87E-03 |
| Dmel_CR32875 | 15.889 | 2.368 | 0.690 | 3.431 | 6.02E-04 | 9.87E-03 |
| Dmel_CG10334 | 1524.281 | 0.510 | 0.149 | 3.430 | 6.04E-04 | 9.87E-03 |
| Dmel_CG8595 | 16.500 | 1.613 | 0.471 | 3.428 | 6.08E-04 | 9.91E-03 |
| Dmel_CG14898 | 511.421 | 0.874 | 0.255 | 3.425 | 6.15E-04 | 1.00E-02 |
| Dmel_CG44246 | 646.353 | 0.760 | 0.222 | 3.423 | 6.20E-04 | 1.01E-02 |
| Dmel_CG7224 | 1716.811 | 0.836 | 0.244 | 3.421 | 6.24E-04 | 1.01E-02 |
| Dmel_CG15269 | 8.248 | -3.161 | 0.924 | -3.420 | 6.26E-04 | 1.01E-02 |
| Dmel_CG40160 | 3039.300 | 0.735 | 0.215 | 3.418 | 6.31E-04 | 1.02E-02 |
| Dmel_CR42452 | 1770.822 | 1.031 | 0.302 | 3.417 | 6.34E-04 | 1.02E-02 |
| Dmel_CG9919 | 9.509 | -2.393 | 0.701 | -3.415 | 6.38E-04 | 1.03E-02 |
| Dmel_CG16896 | 1701.150 | 0.433 | 0.127 | 3.412 | 6.46E-04 | 1.04E-02 |
| Dmel_CG6604 | 39.186 | 1.492 | 0.437 | 3.412 | 6.46E-04 | 1.04E-02 |
| Dmel_CG3314 | 19145.955 | -1.176 | 0.345 | -3.411 | 6.48E-04 | 1.04E-02 |
| Dmel_CG14218 | 17.215 | 2.344 | 0.688 | 3.407 | 6.56E-04 | 1.05E-02 |
| Dmel_CG13982 | 106.094 | -1.105 | 0.324 | -3.407 | 6.58E-04 | 1.05E-02 |
| Dmel_CG10514 | 44.024 | -3.242 | 0.952 | -3.404 | 6.64E-04 | 1.06E-02 |
| Dmel_CG4859 | 83.161 | -1.139 | 0.335 | -3.404 | 6.65E-04 | 1.06E-02 |
| Dmel_CG18507 | 93.141 | 1.294 | 0.381 | 3.401 | 6.71E-04 | 1.07E-02 |
| Dmel_CG8550 | 20.847 | -2.053 | 0.604 | -3.402 | 6.69E-04 | 1.07E-02 |
| Dmel_CG9353 | 452.173 | -1.013 | 0.298 | -3.401 | 6.70E-04 | 1.07E-02 |
| Dmel_CR43589 | 107.488 | 1.689 | 0.497 | 3.400 | 6.75E-04 | 1.07E-02 |
| Dmel_CG14995 | 758.069 | 0.822 | 0.242 | 3.397 | 6.81E-04 | 1.08E-02 |
| Dmel_CG42606 | 8.504 | 2.328 | 0.685 | 3.397 | 6.80E-04 | 1.08E-02 |
| Dmel_CG11284 | 4635.783 | 0.581 | 0.171 | 3.390 | 7.00E-04 | 1.11E-02 |
| Dmel_CG12342 | 308.203 | 1.065 | 0.314 | 3.389 | 7.01E-04 | 1.11E-02 |
| Dmel_CG42365 | 650.988 | -0.730 | 0.215 | -3.387 | 7.05E-04 | 1.11E-02 |

| | | | | | | |
|---|---|---|---|---|---|---|
| Dmel_CG5096 | 38.054 | -1.443 | 0.427 | -3.382 | 7.18E-04 | 1.13E-02 |
| Dmel_CG3984 | 9.224 | 3.722 | 1.101 | 3.382 | 7.20E-04 | 1.13E-02 |
| Dmel_CR44756 | 369.555 | 0.923 | 0.273 | 3.378 | 7.30E-04 | 1.15E-02 |
| Dmel_CG4700 | 8378.432 | 0.563 | 0.167 | 3.376 | 7.37E-04 | 1.15E-02 |
| Dmel_CG9177 | 13081.628 | 0.723 | 0.214 | 3.375 | 7.38E-04 | 1.15E-02 |
| Dmel_CG6718 | 2868.486 | 0.799 | 0.237 | 3.374 | 7.40E-04 | 1.16E-02 |
| Dmel_CG46440 | 920.684 | 0.589 | 0.175 | 3.374 | 7.42E-04 | 1.16E-02 |
| Dmel_CG18131 | 15.976 | -1.772 | 0.525 | -3.372 | 7.45E-04 | 1.16E-02 |
| Dmel_CR44472 | 23.272 | 1.695 | 0.503 | 3.373 | 7.44E-04 | 1.16E-02 |
| Dmel_CG8891 | 326.888 | -1.078 | 0.320 | -3.369 | 7.55E-04 | 1.17E-02 |
| Dmel_CG43346 | 278.496 | -1.055 | 0.313 | -3.368 | 7.58E-04 | 1.17E-02 |
| Dmel_CG14053 | 287.028 | 0.654 | 0.194 | 3.366 | 7.64E-04 | 1.18E-02 |
| Dmel_CG4484 | 163.176 | 1.219 | 0.362 | 3.363 | 7.72E-04 | 1.19E-02 |
| Dmel_CG7449 | 30.833 | 1.278 | 0.380 | 3.361 | 7.75E-04 | 1.20E-02 |
| Dmel_CR45171 | 45.239 | -1.316 | 0.392 | -3.359 | 7.82E-04 | 1.20E-02 |
| Dmel_CG31075 | 302.346 | -1.178 | 0.351 | -3.358 | 7.85E-04 | 1.21E-02 |
| Dmel_CG31495 | 548.374 | -0.665 | 0.198 | -3.358 | 7.85E-04 | 1.21E-02 |
| Dmel_CG2239 | 26.138 | -1.554 | 0.463 | -3.354 | 7.96E-04 | 1.22E-02 |
| Dmel_CR34555 | 357.248 | 1.901 | 0.567 | 3.352 | 8.03E-04 | 1.23E-02 |
| Dmel_CG30361 | 21.616 | 1.814 | 0.541 | 3.351 | 8.04E-04 | 1.23E-02 |
| Dmel_CG5370 | 1911.098 | -0.532 | 0.159 | -3.349 | 8.11E-04 | 1.24E-02 |
| Dmel_CG9331 | 286.658 | -0.670 | 0.200 | -3.347 | 8.17E-04 | 1.25E-02 |
| Dmel_CG9173 | 16.362 | 1.790 | 0.535 | 3.346 | 8.19E-04 | 1.25E-02 |
| Dmel_CG12789 | 42.944 | -1.243 | 0.371 | -3.345 | 8.22E-04 | 1.25E-02 |
| Dmel_CG9586 | 483.605 | -0.903 | 0.270 | -3.344 | 8.25E-04 | 1.25E-02 |
| Dmel_CR34531 | 17.468 | -2.141 | 0.641 | -3.343 | 8.30E-04 | 1.26E-02 |
| Dmel_CG34235 | 13.980 | 1.569 | 0.470 | 3.341 | 8.34E-04 | 1.26E-02 |
| Dmel_CG13855 | 5.891 | -3.022 | 0.905 | -3.339 | 8.41E-04 | 1.27E-02 |
| Dmel_CG10315 | 393.534 | -0.984 | 0.295 | -3.333 | 8.58E-04 | 1.30E-02 |
| Dmel_CG13833 | 4.924 | -3.643 | 1.093 | -3.332 | 8.61E-04 | 1.30E-02 |
| Dmel_CG8389 | 831.413 | 0.536 | 0.161 | 3.331 | 8.64E-04 | 1.30E-02 |

| | | | | | | |
|---|---|---|---|---|---|---|
| Dmel_CR45121 | 50.125 | 1.516 | 0.455 | 3.331 | 8.64E-04 | 1.30E-02 |
| Dmel_CG12344 | 38.021 | -1.307 | 0.392 | -3.330 | 8.69E-04 | 1.31E-02 |
| Dmel_CG31666 | 208.145 | -0.975 | 0.293 | -3.327 | 8.76E-04 | 1.31E-02 |
| Dmel_CG15362 | 501.697 | -0.726 | 0.218 | -3.327 | 8.79E-04 | 1.32E-02 |
| Dmel_CG6282 | 29.876 | -1.687 | 0.508 | -3.322 | 8.93E-04 | 1.34E-02 |
| Dmel_CG18155 | 80.770 | 1.100 | 0.331 | 3.321 | 8.97E-04 | 1.34E-02 |
| Dmel_CG18550 | 29.990 | -1.155 | 0.348 | -3.321 | 8.97E-04 | 1.34E-02 |
| Dmel_CG14644 | 19.112 | -1.835 | 0.553 | -3.319 | 9.03E-04 | 1.34E-02 |
| Dmel_CG33978 | 236.502 | -1.110 | 0.335 | -3.316 | 9.12E-04 | 1.36E-02 |
| Dmel_CG6953 | 7.399 | -2.783 | 0.841 | -3.308 | 9.38E-04 | 1.39E-02 |
| Dmel_CG9155 | 396.786 | -0.873 | 0.264 | -3.308 | 9.39E-04 | 1.39E-02 |
| Dmel_CR44841 | 147.190 | 2.036 | 0.616 | 3.304 | 9.53E-04 | 1.41E-02 |
| Dmel_CR43278 | 17.357 | 1.816 | 0.550 | 3.303 | 9.56E-04 | 1.41E-02 |
| Dmel_CG16727 | 9.877 | -2.937 | 0.890 | -3.302 | 9.59E-04 | 1.42E-02 |
| Dmel_CG43772 | 8.841 | 2.289 | 0.694 | 3.299 | 9.70E-04 | 1.43E-02 |
| Dmel_CG30058 | 14.018 | 1.569 | 0.476 | 3.295 | 9.85E-04 | 1.45E-02 |
| Dmel_CG9503 | 341.519 | -0.705 | 0.214 | -3.294 | 9.86E-04 | 1.45E-02 |
| Dmel_CG8453 | 48.480 | -2.074 | 0.630 | -3.292 | 9.95E-04 | 1.46E-02 |
| Dmel_CG6202 | 3774.453 | 0.413 | 0.125 | 3.290 | 1.00E-03 | 1.47E-02 |
| Dmel_CG9203 | 680.819 | 0.559 | 0.170 | 3.284 | 1.02E-03 | 1.50E-02 |
| Dmel_CG10424 | 667.728 | -0.482 | 0.147 | -3.282 | 1.03E-03 | 1.51E-02 |
| Dmel_CR34535 | 12.981 | -2.039 | 0.621 | -3.283 | 1.03E-03 | 1.51E-02 |
| Dmel_CR45335 | 4.273 | -3.693 | 1.126 | -3.280 | 1.04E-03 | 1.51E-02 |
| Dmel_CR46485 | 143.767 | 0.819 | 0.250 | 3.280 | 1.04E-03 | 1.51E-02 |
| Dmel_CG5381 | 714.190 | -0.894 | 0.273 | -3.277 | 1.05E-03 | 1.53E-02 |
| Dmel_CG42292 | 4.012 | -3.931 | 1.201 | -3.274 | 1.06E-03 | 1.54E-02 |
| Dmel_CR45225 | 15.865 | -1.435 | 0.438 | -3.274 | 1.06E-03 | 1.54E-02 |
| Dmel_CG11500 | 360.013 | -0.970 | 0.297 | -3.270 | 1.07E-03 | 1.55E-02 |
| Dmel_CG32639 | 35.139 | 1.443 | 0.441 | 3.271 | 1.07E-03 | 1.55E-02 |
| Dmel_CG30428 | 377.270 | 0.990 | 0.303 | 3.269 | 1.08E-03 | 1.56E-02 |
| Dmel_CG5905 | 30.547 | -1.520 | 0.465 | -3.268 | 1.08E-03 | 1.57E-02 |

| | | | | | | |
|---|---|---|---|---|---|---|
| Dmel_CR34570 | 18.906 | 1.670 | 0.511 | 3.266 | 1.09E-03 | 1.57E-02 |
| Dmel_CG12484 | 15.035 | 1.633 | 0.500 | 3.264 | 1.10E-03 | 1.58E-02 |
| Dmel_CG9509 | 63.869 | -1.305 | 0.400 | -3.264 | 1.10E-03 | 1.58E-02 |
| Dmel_CG12493 | 49.069 | -1.106 | 0.339 | -3.262 | 1.11E-03 | 1.59E-02 |
| Dmel_CR46048 | 204.191 | 1.157 | 0.355 | 3.262 | 1.11E-03 | 1.59E-02 |
| Dmel_CG12242 | 24.371 | -1.881 | 0.577 | -3.258 | 1.12E-03 | 1.61E-02 |
| Dmel_CR33987 | 5.856 | -3.176 | 0.976 | -3.253 | 1.14E-03 | 1.63E-02 |
| Dmel_CG11254 | 2543.570 | 0.494 | 0.152 | 3.253 | 1.14E-03 | 1.63E-02 |
| Dmel_CR45052 | 9.726 | 1.810 | 0.557 | 3.252 | 1.15E-03 | 1.63E-02 |
| Dmel_CG3180 | 5430.417 | 0.472 | 0.145 | 3.251 | 1.15E-03 | 1.64E-02 |
| Dmel_CG13162 | 4958.261 | 0.650 | 0.200 | 3.248 | 1.16E-03 | 1.65E-02 |
| Dmel_CG12355 | 88.195 | 1.418 | 0.438 | 3.239 | 1.20E-03 | 1.70E-02 |
| Dmel_CG31661 | 694.811 | 1.283 | 0.396 | 3.239 | 1.20E-03 | 1.70E-02 |
| Dmel_CG3533 | 709.897 | 0.595 | 0.184 | 3.240 | 1.20E-03 | 1.70E-02 |
| Dmel_CG31760 | 6.061 | -2.663 | 0.823 | -3.235 | 1.22E-03 | 1.72E-02 |
| Dmel_CG3526 | 23.555 | 1.393 | 0.431 | 3.234 | 1.22E-03 | 1.72E-02 |
| Dmel_CG30345 | 500.338 | -0.774 | 0.240 | -3.227 | 1.25E-03 | 1.76E-02 |
| Dmel_CG11937 | 394.841 | 1.254 | 0.389 | 3.226 | 1.26E-03 | 1.77E-02 |
| Dmel_CG12692 | 6.274 | -2.248 | 0.697 | -3.226 | 1.26E-03 | 1.77E-02 |
| Dmel_CG7938 | 124.712 | -1.225 | 0.380 | -3.225 | 1.26E-03 | 1.77E-02 |
| Dmel_CG9281 | 6711.799 | 0.780 | 0.242 | 3.223 | 1.27E-03 | 1.78E-02 |
| Dmel_CG14620 | 142.968 | 0.811 | 0.252 | 3.220 | 1.28E-03 | 1.79E-02 |
| Dmel_CG43161 | 11.588 | -1.841 | 0.572 | -3.220 | 1.28E-03 | 1.79E-02 |
| Dmel_CG12295 | 35.781 | 1.357 | 0.422 | 3.217 | 1.30E-03 | 1.81E-02 |
| Dmel_CG32572 | 6.465 | 2.419 | 0.752 | 3.215 | 1.30E-03 | 1.82E-02 |
| Dmel_CG3964 | 83.092 | -1.019 | 0.317 | -3.215 | 1.30E-03 | 1.82E-02 |
| Dmel_CG8532 | 1616.996 | -0.506 | 0.157 | -3.216 | 1.30E-03 | 1.82E-02 |
| Dmel_CG1618 | 539.997 | 0.960 | 0.299 | 3.214 | 1.31E-03 | 1.82E-02 |
| Dmel_CG7672 | 11.702 | 1.904 | 0.593 | 3.211 | 1.32E-03 | 1.84E-02 |
| Dmel_CG14419 | 12.691 | 1.942 | 0.605 | 3.209 | 1.33E-03 | 1.85E-02 |
| Dmel_CG3239 | 15.410 | -3.234 | 1.008 | -3.208 | 1.34E-03 | 1.85E-02 |

| | | | | | | |
|---|---|---|---|---|---|---|
| Dmel_CG4821 | 225.884 | -0.959 | 0.299 | -3.206 | 1.35E-03 | 1.86E-02 |
| Dmel_CG12194 | 17.091 | -1.938 | 0.605 | -3.204 | 1.36E-03 | 1.88E-02 |
| Dmel_CG3252 | 14.207 | -2.242 | 0.701 | -3.199 | 1.38E-03 | 1.90E-02 |
| Dmel_CG10207 | 23.822 | -2.141 | 0.670 | -3.197 | 1.39E-03 | 1.91E-02 |
| Dmel_CG6449 | 103.030 | -0.934 | 0.292 | -3.197 | 1.39E-03 | 1.91E-02 |
| Dmel_CG7221 | 353.424 | 0.505 | 0.158 | 3.192 | 1.41E-03 | 1.94E-02 |
| Dmel_CR43866 | 19.789 | 1.269 | 0.397 | 3.192 | 1.41E-03 | 1.94E-02 |
| Dmel_CR46451 | 4.256 | -2.936 | 0.920 | -3.192 | 1.42E-03 | 1.94E-02 |
| Dmel_CG10630 | 13.277 | -2.498 | 0.783 | -3.191 | 1.42E-03 | 1.94E-02 |
| Dmel_CG43968 | 34.300 | -1.639 | 0.514 | -3.190 | 1.42E-03 | 1.94E-02 |
| Dmel_CG42326 | 4.987 | -2.601 | 0.816 | -3.189 | 1.43E-03 | 1.95E-02 |
| Dmel_CG7083 | 1256.892 | 0.491 | 0.154 | 3.187 | 1.44E-03 | 1.96E-02 |
| Dmel_CG11951 | 16.133 | -1.770 | 0.556 | -3.181 | 1.47E-03 | 2.00E-02 |
| Dmel_CG33988 | 8.586 | -1.871 | 0.588 | -3.181 | 1.47E-03 | 2.00E-02 |
| Dmel_CG42732 | 1224.002 | 0.612 | 0.193 | 3.180 | 1.47E-03 | 2.00E-02 |
| Dmel_CG11255 | 998.369 | -0.899 | 0.283 | -3.179 | 1.48E-03 | 2.01E-02 |
| Dmel_CG3918 | 1166.254 | -0.917 | 0.289 | -3.177 | 1.49E-03 | 2.02E-02 |
| Dmel_CG10444 | 1982.603 | -0.834 | 0.263 | -3.176 | 1.49E-03 | 2.02E-02 |
| Dmel_CG11387 | 867.178 | 0.889 | 0.280 | 3.172 | 1.51E-03 | 2.05E-02 |
| Dmel_CG2857 | 23.133 | -1.681 | 0.530 | -3.172 | 1.51E-03 | 2.05E-02 |
| Dmel_CG4496 | 194.538 | 0.797 | 0.252 | 3.166 | 1.55E-03 | 2.08E-02 |
| Dmel_CG4715 | 932.204 | 1.110 | 0.351 | 3.165 | 1.55E-03 | 2.08E-02 |
| Dmel_CG7054 | 954.374 | 0.660 | 0.209 | 3.165 | 1.55E-03 | 2.08E-02 |
| Dmel_CR44987 | 153.397 | 0.870 | 0.275 | 3.166 | 1.55E-03 | 2.08E-02 |
| Dmel_CG13871 | 15.439 | 2.691 | 0.851 | 3.163 | 1.56E-03 | 2.10E-02 |
| Dmel_CG7497 | 146.135 | -0.872 | 0.276 | -3.162 | 1.57E-03 | 2.10E-02 |
| Dmel_CG3176 | 41.451 | -1.512 | 0.478 | -3.161 | 1.57E-03 | 2.10E-02 |
| Dmel_CR45471 | 11.922 | 1.836 | 0.581 | 3.161 | 1.57E-03 | 2.10E-02 |
| Dmel_CG15102 | 2050.109 | -0.525 | 0.166 | -3.160 | 1.58E-03 | 2.11E-02 |
| Dmel_CG44880 | 229.239 | 0.758 | 0.240 | 3.160 | 1.58E-03 | 2.11E-02 |
| Dmel_CG30106 | 16.974 | -2.048 | 0.649 | -3.154 | 1.61E-03 | 2.14E-02 |

| | | | | | | |
|---|---|---|---|---|---|---|
| Dmel_CG5059 | 2186.489 | 0.800 | 0.254 | 3.152 | 1.62E-03 | 2.16E-02 |
| Dmel_CG32373 | 792.364 | 1.121 | 0.356 | 3.148 | 1.65E-03 | 2.18E-02 |
| Dmel_CR46029 | 29.522 | -1.437 | 0.457 | -3.148 | 1.64E-03 | 2.18E-02 |
| Dmel_CG33181 | 2963.425 | 0.751 | 0.239 | 3.146 | 1.65E-03 | 2.19E-02 |
| Dmel_CG5041 | 570.781 | 0.613 | 0.195 | 3.144 | 1.67E-03 | 2.20E-02 |
| Dmel_CG11892 | 208.524 | -2.690 | 0.856 | -3.142 | 1.68E-03 | 2.22E-02 |
| Dmel_CG8864 | 7.182 | -3.023 | 0.962 | -3.142 | 1.68E-03 | 2.22E-02 |
| Dmel_CG5391 | 5.261 | 2.931 | 0.933 | 3.140 | 1.69E-03 | 2.23E-02 |
| Dmel_CG33126 | 197.114 | -1.147 | 0.366 | -3.137 | 1.71E-03 | 2.24E-02 |
| Dmel_CG9460 | 9.877 | -1.909 | 0.608 | -3.137 | 1.70E-03 | 2.24E-02 |
| Dmel_CG33140 | 6.012 | -2.506 | 0.800 | -3.134 | 1.73E-03 | 2.27E-02 |
| Dmel_CG5835 | 43.348 | -0.978 | 0.312 | -3.133 | 1.73E-03 | 2.27E-02 |
| Dmel_CG3254 | 25.889 | -1.471 | 0.470 | -3.132 | 1.74E-03 | 2.28E-02 |
| Dmel_CG34161 | 8.362 | -2.108 | 0.674 | -3.129 | 1.75E-03 | 2.29E-02 |
| Dmel_CR34335 | 36494.636 | -1.023 | 0.327 | -3.127 | 1.76E-03 | 2.31E-02 |
| Dmel_CG18870 | 2820.357 | 0.447 | 0.143 | 3.126 | 1.77E-03 | 2.32E-02 |
| Dmel_CG10702 | 1150.706 | 0.814 | 0.261 | 3.124 | 1.78E-03 | 2.33E-02 |
| Dmel_CG10734 | 120.828 | 0.968 | 0.310 | 3.121 | 1.80E-03 | 2.34E-02 |
| Dmel_CG13784 | 2273.112 | 0.439 | 0.141 | 3.120 | 1.81E-03 | 2.35E-02 |
| Dmel_CR32957 | 2001.447 | 0.688 | 0.221 | 3.120 | 1.81E-03 | 2.35E-02 |
| Dmel_CG6503 | 26.425 | -3.794 | 1.216 | -3.119 | 1.81E-03 | 2.35E-02 |
| Dmel_CG30334 | 16.024 | 1.724 | 0.553 | 3.119 | 1.82E-03 | 2.35E-02 |
| Dmel_CG34331 | 7.516 | -2.857 | 0.917 | -3.114 | 1.85E-03 | 2.39E-02 |
| Dmel_CG6584 | 2008.800 | 0.434 | 0.139 | 3.113 | 1.85E-03 | 2.40E-02 |
| Dmel_CG4733 | 177.034 | 0.838 | 0.269 | 3.111 | 1.86E-03 | 2.41E-02 |
| Dmel_CG9456 | 35.892 | -1.475 | 0.475 | -3.108 | 1.89E-03 | 2.43E-02 |
| Dmel_CG9650 | 454.888 | -0.791 | 0.255 | -3.106 | 1.90E-03 | 2.45E-02 |
| Dmel_CG15309 | 1390.190 | 0.451 | 0.145 | 3.105 | 1.90E-03 | 2.45E-02 |
| Dmel_CR43962 | 8.553 | -2.440 | 0.786 | -3.103 | 1.92E-03 | 2.47E-02 |
| Dmel_CG2060 | 1551.905 | 0.661 | 0.213 | 3.102 | 1.92E-03 | 2.47E-02 |
| Dmel_CG41284 | 85.265 | 1.643 | 0.531 | 3.096 | 1.96E-03 | 2.51E-02 |

| | | | | | | |
|---|---|---|---|---|---|---|
| Dmel_CG8380 | 8.756 | 1.965 | 0.635 | 3.095 | 1.97E-03 | 2.52E-02 |
| Dmel_CG31365 | 1258.248 | 0.778 | 0.251 | 3.094 | 1.98E-03 | 2.53E-02 |
| Dmel_CG5612 | 169.169 | -0.991 | 0.321 | -3.091 | 2.00E-03 | 2.55E-02 |
| Dmel_CG8083 | 13.191 | -1.741 | 0.563 | -3.091 | 2.00E-03 | 2.55E-02 |
| Dmel_CG3705 | 1508.729 | -0.635 | 0.205 | -3.090 | 2.00E-03 | 2.55E-02 |
| Dmel_CG13282 | 99.633 | -1.054 | 0.341 | -3.088 | 2.01E-03 | 2.56E-02 |
| Dmel_CG16799 | 9.929 | -2.275 | 0.737 | -3.087 | 2.02E-03 | 2.57E-02 |
| Dmel_CG12346 | 351.844 | -0.783 | 0.254 | -3.084 | 2.04E-03 | 2.59E-02 |
| Dmel_CG10005 | 59.957 | -1.055 | 0.342 | -3.084 | 2.04E-03 | 2.59E-02 |
| Dmel_CG12002 | 217.218 | -1.329 | 0.432 | -3.079 | 2.08E-03 | 2.63E-02 |
| Dmel_CG12684 | 9.256 | 2.071 | 0.673 | 3.079 | 2.08E-03 | 2.63E-02 |
| Dmel_CG17795 | 17.109 | -1.962 | 0.638 | -3.077 | 2.09E-03 | 2.64E-02 |
| Dmel_CG17217 | 8.090 | -2.152 | 0.700 | -3.075 | 2.10E-03 | 2.65E-02 |
| Dmel_CG6071 | 45.004 | 1.312 | 0.427 | 3.076 | 2.10E-03 | 2.65E-02 |
| Dmel_CG18548 | 12.434 | -1.665 | 0.542 | -3.074 | 2.11E-03 | 2.66E-02 |
| Dmel_CG7595 | 4521.269 | 0.400 | 0.130 | 3.072 | 2.13E-03 | 2.67E-02 |
| Dmel_CG11634 | 4.279 | -3.755 | 1.222 | -3.071 | 2.13E-03 | 2.68E-02 |
| Dmel_CG9207 | 300.491 | -0.757 | 0.247 | -3.069 | 2.15E-03 | 2.69E-02 |
| Dmel_CR33686 | 13262.350 | 0.888 | 0.289 | 3.068 | 2.15E-03 | 2.70E-02 |
| Dmel_CG17716 | 19.533 | -1.772 | 0.578 | -3.066 | 2.17E-03 | 2.71E-02 |
| Dmel_CG40298 | 76.791 | 1.427 | 0.465 | 3.065 | 2.17E-03 | 2.72E-02 |
| Dmel_CG17970 | 165.747 | 0.974 | 0.318 | 3.063 | 2.19E-03 | 2.74E-02 |
| Dmel_CG7041 | 1004.052 | 0.604 | 0.197 | 3.063 | 2.19E-03 | 2.74E-02 |
| Dmel_CG8339 | 1557.479 | 0.337 | 0.110 | 3.063 | 2.19E-03 | 2.74E-02 |
| Dmel_CG32506 | 9.534 | -1.944 | 0.635 | -3.062 | 2.20E-03 | 2.74E-02 |
| Dmel_CG9772 | 2989.847 | 0.486 | 0.159 | 3.060 | 2.21E-03 | 2.75E-02 |
| Dmel_CG30047 | 10.082 | -2.525 | 0.827 | -3.055 | 2.25E-03 | 2.80E-02 |
| Dmel_CG4257 | 9979.136 | 0.364 | 0.119 | 3.054 | 2.26E-03 | 2.80E-02 |
| Dmel_CG5778 | 6.358 | -1.972 | 0.646 | -3.052 | 2.27E-03 | 2.81E-02 |
| Dmel_CG6677 | 880.216 | 0.618 | 0.203 | 3.052 | 2.27E-03 | 2.81E-02 |
| Dmel_CR45534 | 7.894 | 1.872 | 0.614 | 3.048 | 2.30E-03 | 2.85E-02 |

| | | | | | | |
|---|---|---|---|---|---|---|
| Dmel_CG1634 | 2307.073 | -0.690 | 0.227 | -3.047 | 2.31E-03 | 2.85E-02 |
| Dmel_CG1806 | 80.643 | 0.842 | 0.276 | 3.046 | 2.32E-03 | 2.86E-02 |
| Dmel_CG15576 | 10.175 | -2.048 | 0.672 | -3.045 | 2.32E-03 | 2.86E-02 |
| Dmel_CG33093 | 9.378 | 2.029 | 0.667 | 3.041 | 2.36E-03 | 2.90E-02 |
| Dmel_CG11899 | 1333.295 | 0.723 | 0.238 | 3.039 | 2.37E-03 | 2.92E-02 |
| Dmel_CG34166 | 13.640 | -3.258 | 1.073 | -3.036 | 2.39E-03 | 2.94E-02 |
| Dmel_CG6303 | 18054.014 | 0.312 | 0.103 | 3.035 | 2.40E-03 | 2.95E-02 |
| Dmel_CG12120 | 10.092 | -1.932 | 0.637 | -3.034 | 2.41E-03 | 2.95E-02 |
| Dmel_CG17524 | 127.121 | -0.831 | 0.274 | -3.033 | 2.42E-03 | 2.96E-02 |
| Dmel_CG6575 | 11723.986 | 0.711 | 0.234 | 3.033 | 2.42E-03 | 2.96E-02 |
| Dmel_CG3905 | 717.826 | 0.548 | 0.181 | 3.031 | 2.44E-03 | 2.98E-02 |
| Dmel_CG9677 | 7820.096 | 0.424 | 0.140 | 3.030 | 2.45E-03 | 2.99E-02 |
| Dmel_CG5904 | 373.382 | 0.587 | 0.194 | 3.028 | 2.46E-03 | 2.99E-02 |
| Dmel_CG7913 | 6017.463 | 0.432 | 0.143 | 3.028 | 2.46E-03 | 2.99E-02 |
| Dmel_CG3971 | 4591.617 | 0.532 | 0.176 | 3.027 | 2.47E-03 | 3.01E-02 |
| Dmel_CG13315 | 109.740 | -1.502 | 0.497 | -3.026 | 2.48E-03 | 3.01E-02 |
| Dmel_CR44115 | 21.000 | -2.527 | 0.836 | -3.023 | 2.50E-03 | 3.04E-02 |
| Dmel_CG31673 | 855.982 | -0.521 | 0.173 | -3.022 | 2.51E-03 | 3.04E-02 |
| Dmel_CR43018 | 20.119 | -5.429 | 1.797 | -3.022 | 2.51E-03 | 3.04E-02 |
| Dmel_CG32656 | 29.905 | -1.495 | 0.495 | -3.021 | 2.52E-03 | 3.05E-02 |
| Dmel_CG13663 | 409.022 | -0.692 | 0.229 | -3.019 | 2.53E-03 | 3.06E-02 |
| Dmel_CG15040 | 6.619 | -2.235 | 0.741 | -3.018 | 2.54E-03 | 3.07E-02 |
| Dmel_CG14275 | 101.798 | 0.985 | 0.326 | 3.017 | 2.55E-03 | 3.07E-02 |
| Dmel_CR33674 | 23.624 | 1.375 | 0.456 | 3.016 | 2.56E-03 | 3.08E-02 |
| Dmel_CG30195 | 12.984 | -1.876 | 0.623 | -3.014 | 2.58E-03 | 3.09E-02 |
| Dmel_CG6511 | 1515.921 | 0.437 | 0.145 | 3.014 | 2.57E-03 | 3.09E-02 |
| Dmel_CG8808 | 3822.949 | 0.618 | 0.205 | 3.012 | 2.59E-03 | 3.11E-02 |
| Dmel_CG11064 | 2076.511 | -1.666 | 0.554 | -3.007 | 2.64E-03 | 3.16E-02 |
| Dmel_CG6495 | 14.977 | 2.389 | 0.795 | 3.004 | 2.67E-03 | 3.19E-02 |
| Dmel_CG33960 | 90.640 | 1.186 | 0.395 | 3.002 | 2.69E-03 | 3.21E-02 |
| Dmel_CG42249 | 22.299 | -1.121 | 0.374 | -3.000 | 2.70E-03 | 3.22E-02 |

| | | | | | | |
|---|---|---|---|---|---|---|
| Dmel_CG9242 | 7363.625 | 0.548 | 0.183 | 3.000 | 2.70E-03 | 3.22E-02 |
| Dmel_CG31898 | 1376.311 | 0.923 | 0.308 | 2.996 | 2.74E-03 | 3.26E-02 |
| Dmel_CR42491 | 542.160 | -0.920 | 0.307 | -2.995 | 2.74E-03 | 3.26E-02 |
| Dmel_CG2102 | 31.808 | -1.577 | 0.527 | -2.995 | 2.75E-03 | 3.26E-02 |
| Dmel_CG31445 | 45.239 | -1.471 | 0.491 | -2.993 | 2.76E-03 | 3.27E-02 |
| Dmel_CG43295 | 68.057 | -0.972 | 0.325 | -2.993 | 2.76E-03 | 3.27E-02 |
| Dmel_CG5337 | 106.836 | -0.633 | 0.212 | -2.993 | 2.76E-03 | 3.27E-02 |
| Dmel_CG10466 | 102.353 | -1.024 | 0.342 | -2.992 | 2.77E-03 | 3.28E-02 |
| Dmel_CG11783 | 659.129 | -0.599 | 0.200 | -2.989 | 2.80E-03 | 3.30E-02 |
| Dmel_CG3504 | 37.047 | -1.060 | 0.355 | -2.989 | 2.80E-03 | 3.30E-02 |
| Dmel_CG4579 | 7553.916 | 0.337 | 0.113 | 2.989 | 2.80E-03 | 3.30E-02 |
| Dmel_CG32146 | 3320.000 | 0.565 | 0.189 | 2.988 | 2.81E-03 | 3.31E-02 |
| Dmel_CG6217 | 22.325 | -1.437 | 0.481 | -2.986 | 2.83E-03 | 3.32E-02 |
| Dmel_CG12179 | 1309.413 | 0.444 | 0.149 | 2.983 | 2.85E-03 | 3.35E-02 |
| Dmel_CG34323 | 38.530 | -1.299 | 0.436 | -2.982 | 2.86E-03 | 3.36E-02 |
| Dmel_CG33926 | 47.226 | -1.752 | 0.588 | -2.981 | 2.87E-03 | 3.37E-02 |
| Dmel_CG7447 | 113.824 | 1.091 | 0.366 | 2.979 | 2.89E-03 | 3.39E-02 |
| Dmel_CR46083 | 34.947 | 1.067 | 0.360 | 2.967 | 3.01E-03 | 3.52E-02 |
| Dmel_CG2706 | 50.078 | -1.306 | 0.441 | -2.966 | 3.02E-03 | 3.53E-02 |
| Dmel_CG9098 | 194.284 | 0.586 | 0.198 | 2.963 | 3.05E-03 | 3.56E-02 |
| Dmel_CG12708 | 73.374 | 1.280 | 0.432 | 2.961 | 3.07E-03 | 3.58E-02 |
| Dmel_CG13827 | 209.725 | 0.751 | 0.254 | 2.958 | 3.09E-03 | 3.60E-02 |
| Dmel_CG2736 | 76.651 | -0.862 | 0.291 | -2.959 | 3.09E-03 | 3.60E-02 |
| Dmel_CR45179 | 43.006 | 1.321 | 0.447 | 2.958 | 3.10E-03 | 3.60E-02 |
| Dmel_CG1950 | 10.505 | -2.311 | 0.783 | -2.950 | 3.18E-03 | 3.69E-02 |
| Dmel_CG9611 | 723.808 | 0.458 | 0.155 | 2.949 | 3.19E-03 | 3.70E-02 |
| Dmel_CR33753 | 103.198 | 1.049 | 0.356 | 2.947 | 3.21E-03 | 3.71E-02 |
| Dmel_CG15848 | 101.067 | 1.135 | 0.386 | 2.944 | 3.24E-03 | 3.75E-02 |
| Dmel_CG2194 | 15.037 | -1.685 | 0.573 | -2.939 | 3.29E-03 | 3.80E-02 |
| Dmel_CR43493 | 219.103 | 0.871 | 0.296 | 2.940 | 3.28E-03 | 3.80E-02 |
| Dmel_CG18372 | 32.607 | -2.602 | 0.885 | -2.938 | 3.30E-03 | 3.81E-02 |

| | | | | | | |
|---|---|---|---|---|---|---|
| Dmel_CG31764 | 398.620 | -0.674 | 0.230 | -2.938 | 3.31E-03 | 3.81E-02 |
| Dmel_CG34392 | 202.441 | 1.223 | 0.417 | 2.935 | 3.34E-03 | 3.85E-02 |
| Dmel_CG42600 | 8452.538 | 0.533 | 0.182 | 2.934 | 3.35E-03 | 3.85E-02 |
| Dmel_CG42739 | 692.104 | 0.780 | 0.266 | 2.933 | 3.36E-03 | 3.86E-02 |
| Dmel_CG46511 | 10.848 | 2.683 | 0.916 | 2.930 | 3.39E-03 | 3.89E-02 |
| Dmel_CG8930 | 31.973 | 1.092 | 0.373 | 2.929 | 3.40E-03 | 3.90E-02 |
| Dmel_CG11152 | 73.962 | 1.292 | 0.441 | 2.928 | 3.41E-03 | 3.91E-02 |
| Dmel_CG14142 | 22.763 | 1.082 | 0.370 | 2.927 | 3.42E-03 | 3.91E-02 |
| Dmel_CG10901 | 69851.455 | 0.493 | 0.169 | 2.927 | 3.43E-03 | 3.92E-02 |
| Dmel_CG9610 | 35.323 | -1.337 | 0.457 | -2.926 | 3.43E-03 | 3.92E-02 |
| Dmel_CG1112 | 34.251 | -1.273 | 0.435 | -2.924 | 3.45E-03 | 3.93E-02 |
| Dmel_CG15035 | 33.968 | 1.521 | 0.520 | 2.925 | 3.45E-03 | 3.93E-02 |
| Dmel_CG9414 | 984.747 | -0.576 | 0.197 | -2.924 | 3.45E-03 | 3.93E-02 |
| Dmel_CG3757 | 66.606 | -1.071 | 0.366 | -2.924 | 3.46E-03 | 3.93E-02 |
| Dmel_CG13309 | 45.049 | -5.610 | 1.920 | -2.922 | 3.47E-03 | 3.94E-02 |
| Dmel_CG3694 | 9.047 | 2.078 | 0.711 | 2.922 | 3.48E-03 | 3.94E-02 |
| Dmel_CG7660 | 20442.794 | 0.369 | 0.126 | 2.922 | 3.48E-03 | 3.94E-02 |
| Dmel_CG42626 | 9.841 | -1.383 | 0.474 | -2.921 | 3.49E-03 | 3.95E-02 |
| Dmel_CG43770 | 12411.707 | 0.490 | 0.168 | 2.921 | 3.49E-03 | 3.95E-02 |
| Dmel_CG7910 | 43.394 | 1.649 | 0.565 | 2.919 | 3.51E-03 | 3.97E-02 |
| Dmel_CG11852 | 53.225 | -1.131 | 0.388 | -2.918 | 3.52E-03 | 3.97E-02 |
| Dmel_CG8014 | 7309.156 | 0.345 | 0.118 | 2.918 | 3.52E-03 | 3.97E-02 |
| Dmel_CG8318 | 6370.629 | 0.435 | 0.149 | 2.915 | 3.55E-03 | 4.00E-02 |
| Dmel_CG30339 | 24.402 | -1.351 | 0.464 | -2.912 | 3.59E-03 | 4.03E-02 |
| Dmel_CG13113 | 571.943 | 1.134 | 0.390 | 2.912 | 3.59E-03 | 4.04E-02 |
| Dmel_CG2849 | 5829.301 | -0.601 | 0.207 | -2.909 | 3.62E-03 | 4.06E-02 |
| Dmel_CG40494 | 5617.941 | 0.382 | 0.131 | 2.909 | 3.62E-03 | 4.06E-02 |
| Dmel_CG2699 | 9132.501 | 0.534 | 0.184 | 2.908 | 3.64E-03 | 4.07E-02 |
| Dmel_CG43370 | 11.643 | -1.723 | 0.593 | -2.908 | 3.64E-03 | 4.07E-02 |
| Dmel_CG42356 | 20.310 | 1.454 | 0.500 | 2.906 | 3.66E-03 | 4.09E-02 |
| Dmel_CG3004 | 1347.285 | 0.378 | 0.130 | 2.904 | 3.69E-03 | 4.11E-02 |

| | | | | | | |
|---|---|---|---|---|---|---|
| Dmel_CG17294 | 289.599 | -0.625 | 0.215 | -2.903 | 3.69E-03 | 4.11E-02 |
| Dmel_CG42357 | 20.533 | 1.583 | 0.545 | 2.903 | 3.69E-03 | 4.11E-02 |
| Dmel_CG30156 | 45.744 | 0.920 | 0.317 | 2.900 | 3.74E-03 | 4.15E-02 |
| Dmel_CR32900 | 11.168 | 1.445 | 0.498 | 2.900 | 3.73E-03 | 4.15E-02 |
| Dmel_CG12605 | 8.184 | -2.030 | 0.701 | -2.894 | 3.80E-03 | 4.22E-02 |
| Dmel_CG45088 | 63.539 | 1.460 | 0.505 | 2.893 | 3.81E-03 | 4.22E-02 |
| Dmel_CG9256 | 405.857 | -0.703 | 0.243 | -2.893 | 3.81E-03 | 4.22E-02 |
| Dmel_CR46094 | 67.098 | 0.875 | 0.302 | 2.894 | 3.81E-03 | 4.22E-02 |
| Dmel_CR46090 | 25.206 | 0.995 | 0.344 | 2.893 | 3.82E-03 | 4.22E-02 |
| Dmel_CG14341 | 124.607 | -1.008 | 0.349 | -2.892 | 3.83E-03 | 4.23E-02 |
| Dmel_CG31038 | 332.270 | 0.559 | 0.193 | 2.892 | 3.83E-03 | 4.23E-02 |
| Dmel_CG34433 | 25.355 | 1.353 | 0.468 | 2.892 | 3.83E-03 | 4.23E-02 |
| Dmel_CG10777 | 10216.604 | 0.507 | 0.175 | 2.889 | 3.86E-03 | 4.25E-02 |
| Dmel_CG13888 | 26.473 | -0.993 | 0.344 | -2.889 | 3.86E-03 | 4.25E-02 |
| Dmel_CG4559 | 243.698 | -0.610 | 0.211 | -2.889 | 3.86E-03 | 4.25E-02 |
| Dmel_CG3259 | 25.044 | -1.221 | 0.423 | -2.888 | 3.87E-03 | 4.25E-02 |
| Dmel_CG15525 | 215.123 | -0.719 | 0.249 | -2.885 | 3.91E-03 | 4.29E-02 |
| Dmel_CG32077 | 13.884 | 1.639 | 0.568 | 2.885 | 3.92E-03 | 4.29E-02 |
| Dmel_CG17129 | 2764.223 | 0.763 | 0.264 | 2.884 | 3.93E-03 | 4.30E-02 |
| Dmel_CG44195 | 214.639 | 0.979 | 0.339 | 2.884 | 3.93E-03 | 4.30E-02 |
| Dmel_CG7607 | 91.260 | 0.863 | 0.299 | 2.883 | 3.93E-03 | 4.30E-02 |
| Dmel_CG4170 | 1895.737 | -0.667 | 0.231 | -2.882 | 3.95E-03 | 4.30E-02 |
| Dmel_CG8665 | 64.322 | -0.946 | 0.328 | -2.882 | 3.95E-03 | 4.30E-02 |
| Dmel_CG17060 | 4727.730 | 0.355 | 0.123 | 2.880 | 3.97E-03 | 4.32E-02 |
| Dmel_CG9492 | 5.030 | -2.490 | 0.864 | -2.880 | 3.97E-03 | 4.32E-02 |
| Dmel_CG13317 | 26.761 | 1.699 | 0.590 | 2.879 | 3.99E-03 | 4.33E-02 |
| Dmel_CG15739 | 27.147 | -1.222 | 0.424 | -2.879 | 3.99E-03 | 4.34E-02 |
| Dmel_CG8651 | 11703.032 | 0.342 | 0.119 | 2.876 | 4.03E-03 | 4.38E-02 |
| Dmel_CG6330 | 698.336 | -0.683 | 0.238 | -2.873 | 4.06E-03 | 4.40E-02 |
| Dmel_CG6542 | 4210.347 | 0.424 | 0.148 | 2.872 | 4.08E-03 | 4.42E-02 |
| Dmel_CG31743 | 245.562 | 0.807 | 0.281 | 2.871 | 4.09E-03 | 4.42E-02 |

| | | | | | | |
|---|---|---|---|---|---|---|
| Dmel_CG3424 | 18475.822 | 0.403 | 0.141 | 2.871 | 4.10E-03 | 4.42E-02 |
| Dmel_CG17717 | 17.339 | -1.491 | 0.520 | -2.866 | 4.15E-03 | 4.48E-02 |
| Dmel_CG44325 | 724.990 | -0.660 | 0.230 | -2.863 | 4.19E-03 | 4.52E-02 |
| Dmel_CG7635 | 8.520 | -2.027 | 0.708 | -2.863 | 4.20E-03 | 4.52E-02 |
| Dmel_CR45047 | 18.644 | 1.176 | 0.411 | 2.863 | 4.20E-03 | 4.52E-02 |
| Dmel_CG11000 | 88.807 | 1.094 | 0.382 | 2.860 | 4.24E-03 | 4.56E-02 |
| Dmel_CG10006 | 27.615 | -1.283 | 0.449 | -2.859 | 4.25E-03 | 4.56E-02 |
| Dmel_CG1774 | 146.285 | -0.977 | 0.342 | -2.857 | 4.28E-03 | 4.58E-02 |
| Dmel_CG31992 | 20913.729 | 0.405 | 0.142 | 2.856 | 4.29E-03 | 4.58E-02 |
| Dmel_CG4180 | 1002.902 | -0.407 | 0.143 | -2.856 | 4.29E-03 | 4.58E-02 |
| Dmel_CG7134 | 1948.993 | 0.544 | 0.190 | 2.857 | 4.28E-03 | 4.58E-02 |
| Dmel_CG9682 | 8.008 | -1.986 | 0.695 | -2.855 | 4.30E-03 | 4.60E-02 |
| Dmel_CG3544 | 5.634 | -2.084 | 0.730 | -2.854 | 4.31E-03 | 4.60E-02 |
| Dmel_CG8827 | 140.828 | -1.011 | 0.354 | -2.854 | 4.31E-03 | 4.60E-02 |
| Dmel_CG30343 | 277.129 | -0.634 | 0.222 | -2.853 | 4.34E-03 | 4.61E-02 |
| Dmel_CG46520 | 3.070 | -5.281 | 1.851 | -2.853 | 4.34E-03 | 4.61E-02 |
| Dmel_CG46522 | 3.070 | -5.281 | 1.851 | -2.853 | 4.34E-03 | 4.61E-02 |
| Dmel_CG10146 | 15.302 | -2.344 | 0.823 | -2.849 | 4.39E-03 | 4.66E-02 |
| Dmel_CG42854 | 30.574 | -1.603 | 0.563 | -2.849 | 4.39E-03 | 4.66E-02 |
| Dmel_CG12423 | 221.162 | -0.929 | 0.326 | -2.845 | 4.44E-03 | 4.71E-02 |
| Dmel_CG6490 | 2609.600 | 0.762 | 0.268 | 2.845 | 4.44E-03 | 4.71E-02 |
| Dmel_CG8805 | 2531.220 | 0.438 | 0.154 | 2.844 | 4.45E-03 | 4.71E-02 |
| Dmel_CG9372 | 9.420 | -1.894 | 0.667 | -2.840 | 4.52E-03 | 4.77E-02 |
| Dmel_CR34607 | 29.650 | 1.410 | 0.497 | 2.838 | 4.53E-03 | 4.79E-02 |
| Dmel_CG14193 | 12.461 | -1.601 | 0.565 | -2.837 | 4.56E-03 | 4.80E-02 |
| Dmel_CG18473 | 60.517 | 1.180 | 0.416 | 2.836 | 4.56E-03 | 4.80E-02 |
| Dmel_CG3039 | 3401.084 | 0.700 | 0.247 | 2.837 | 4.56E-03 | 4.80E-02 |
| Dmel_CG1772 | 8275.607 | 0.661 | 0.233 | 2.833 | 4.61E-03 | 4.84E-02 |
| Dmel_CG8205 | 2676.441 | 0.546 | 0.193 | 2.832 | 4.63E-03 | 4.86E-02 |
| Dmel_CG6207 | 2745.065 | 0.598 | 0.212 | 2.825 | 4.72E-03 | 4.96E-02 |
| Dmel_CG8316 | 213.397 | -0.793 | 0.281 | -2.825 | 4.73E-03 | 4.96E-02 |

| | | | | | | |
|---|---|---|---|---|---|---|
| Dmel_CG10479 | 75.907 | 1.041 | 0.369 | 2.821 | 4.79E-03 | 5.02E-02 |
| Dmel_CR45758 | 6.430 | 2.623 | 0.930 | 2.820 | 4.81E-03 | 5.03E-02 |
| Dmel_CG11128 | 2190.850 | 0.888 | 0.315 | 2.817 | 4.84E-03 | 5.06E-02 |
| Dmel_CR46037 | 8285.229 | -1.108 | 0.393 | -2.817 | 4.84E-03 | 5.06E-02 |
| Dmel_CG43079 | 631.857 | -1.001 | 0.355 | -2.816 | 4.86E-03 | 5.07E-02 |
| Dmel_CG7460 | 93.172 | 1.036 | 0.368 | 2.814 | 4.89E-03 | 5.10E-02 |
| Dmel_CG34220 | 261.387 | 1.093 | 0.388 | 2.813 | 4.91E-03 | 5.11E-02 |
| Dmel_CG14695 | 5.018 | -2.908 | 1.034 | -2.812 | 4.93E-03 | 5.13E-02 |
| Dmel_CG34455 | 1000.243 | 0.458 | 0.163 | 2.808 | 4.98E-03 | 5.18E-02 |
| Dmel_CG15822 | 271.011 | 0.876 | 0.312 | 2.806 | 5.02E-03 | 5.21E-02 |
| Dmel_CG14032 | 170.072 | 1.031 | 0.368 | 2.805 | 5.03E-03 | 5.21E-02 |
| Dmel_CG2902 | 12.707 | -1.599 | 0.570 | -2.802 | 5.08E-03 | 5.26E-02 |
| Dmel_CG3290 | 41.169 | -5.759 | 2.055 | -2.802 | 5.08E-03 | 5.26E-02 |
| Dmel_CG10553 | 15.343 | -2.517 | 0.898 | -2.801 | 5.09E-03 | 5.26E-02 |
| Dmel_CG7470 | 8.831 | -2.032 | 0.725 | -2.801 | 5.10E-03 | 5.27E-02 |
| Dmel_CG32086 | 4.755 | -2.790 | 0.998 | -2.797 | 5.16E-03 | 5.33E-02 |
| Dmel_CG7272 | 93.577 | -0.653 | 0.234 | -2.797 | 5.16E-03 | 5.33E-02 |
| Dmel_CG3241 | 198.254 | 0.608 | 0.217 | 2.795 | 5.19E-03 | 5.35E-02 |
| Dmel_CG11310 | 8.366 | 2.046 | 0.733 | 2.792 | 5.25E-03 | 5.40E-02 |
| Dmel_CR45036 | 19.168 | -1.346 | 0.483 | -2.790 | 5.27E-03 | 5.42E-02 |
| Dmel_CG4006 | 3810.911 | 0.486 | 0.174 | 2.789 | 5.29E-03 | 5.43E-02 |
| Dmel_CG7363 | 14.727 | -1.578 | 0.566 | -2.789 | 5.29E-03 | 5.43E-02 |
| Dmel_CG14064 | 16.145 | 1.903 | 0.683 | 2.788 | 5.31E-03 | 5.44E-02 |
| Dmel_CG1307 | 973.268 | -0.811 | 0.291 | -2.787 | 5.32E-03 | 5.44E-02 |
| Dmel_CG2767 | 1424.389 | 0.545 | 0.195 | 2.787 | 5.32E-03 | 5.44E-02 |

**table S17**. *D. melanogaster* genes Wald Test significant results for ~Genotype vs ~Genotype+Infection+Genotype*Infection

| gene_id | baseMean | log2FoldChange | lfcSE | stat | pvalue | padj |
|---|---|---|---|---|---|---|
| Dmel_CG43079 | 631.857 | -1.956 | 0.356 | -5.494 | 3.92E-08 | 2.56E-04 |
| Dmel_CG9871 | 50.560 | -3.292 | 0.610 | -5.399 | 6.69E-08 | 2.56E-04 |
| Dmel_CG7002 | 154.856 | -2.425 | 0.450 | -5.387 | 7.17E-08 | 2.56E-04 |
| Dmel_CG13772 | 15.539 | -3.435 | 0.646 | -5.318 | 1.05E-07 | 2.80E-04 |
| Dmel_CG10593 | 2009.122 | 2.010 | 0.385 | 5.220 | 1.79E-07 | 3.83E-04 |
| Dmel_CG15599 | 55.457 | -2.867 | 0.557 | -5.151 | 2.59E-07 | 4.61E-04 |
| Dmel_CG6533 | 2736.531 | 4.123 | 0.808 | 5.101 | 3.38E-07 | 5.16E-04 |
| Dmel_CG9073 | 36.796 | -2.448 | 0.487 | -5.031 | 4.89E-07 | 6.54E-04 |
| Dmel_CG6524 | 6268.890 | 3.938 | 0.791 | 4.976 | 6.48E-07 | 7.54E-04 |
| Dmel_CR46481 | 1090.588 | -1.267 | 0.256 | -4.947 | 7.54E-07 | 7.54E-04 |
| Dmel_CG2187 | 500.405 | 1.275 | 0.258 | 4.937 | 7.95E-07 | 7.54E-04 |
| Dmel_CG11941 | 16.019 | -3.213 | 0.653 | -4.924 | 8.46E-07 | 7.54E-04 |
| Dmel_CG8663 | 114.131 | -1.984 | 0.409 | -4.856 | 1.19E-06 | 9.83E-04 |
| Dmel_CG31973 | 444.390 | -1.252 | 0.260 | -4.814 | 1.48E-06 | 1.13E-03 |
| Dmel_CG42711 | 9.934 | -4.252 | 0.888 | -4.790 | 1.66E-06 | 1.19E-03 |
| Dmel_CG10287 | 102.780 | -1.574 | 0.330 | -4.776 | 1.79E-06 | 1.19E-03 |
| Dmel_CR32368 | 12.082 | -3.326 | 0.708 | -4.699 | 2.61E-06 | 1.64E-03 |
| Dmel_CG30046 | 65.427 | 1.508 | 0.322 | 4.685 | 2.80E-06 | 1.65E-03 |
| Dmel_CG8942 | 20.884 | -3.404 | 0.728 | -4.676 | 2.93E-06 | 1.65E-03 |
| Dmel_CG34323 | 38.530 | -2.054 | 0.441 | -4.661 | 3.14E-06 | 1.68E-03 |
| Dmel_CG5644 | 28.092 | 2.236 | 0.486 | 4.598 | 4.26E-06 | 2.17E-03 |
| Dmel_CG4373 | 307.928 | -2.564 | 0.564 | -4.547 | 5.44E-06 | 2.58E-03 |
| Dmel_CG13084 | 2952.416 | 3.172 | 0.700 | 4.532 | 5.83E-06 | 2.58E-03 |
| Dmel_CG4783 | 18.606 | -2.194 | 0.484 | -4.531 | 5.87E-06 | 2.58E-03 |
| Dmel_CG3157 | 1113.612 | 0.878 | 0.194 | 4.524 | 6.08E-06 | 2.58E-03 |
| Dmel_CG1091 | 2685.582 | 0.732 | 0.162 | 4.517 | 6.26E-06 | 2.58E-03 |
| Dmel_CG6517 | 5048.988 | 3.842 | 0.857 | 4.483 | 7.35E-06 | 2.91E-03 |
| Dmel_CG8388 | 689.979 | 0.544 | 0.121 | 4.474 | 7.66E-06 | 2.93E-03 |

| | | | | | | |
|---|---|---|---|---|---|---|
| Dmel_CG3635 | 29.922 | -2.392 | 0.543 | -4.406 | 1.05E-05 | 3.89E-03 |
| Dmel_CG32640 | 167.387 | -1.214 | 0.278 | -4.373 | 1.22E-05 | 4.36E-03 |
| Dmel_CG9663 | 323.692 | 1.203 | 0.276 | 4.355 | 1.33E-05 | 4.59E-03 |
| Dmel_CR46350 | 65.211 | -1.765 | 0.407 | -4.332 | 1.48E-05 | 4.90E-03 |
| Dmel_CG5937 | 33.106 | -2.091 | 0.483 | -4.327 | 1.51E-05 | 4.90E-03 |
| Dmel_CG3625 | 103.288 | -0.979 | 0.228 | -4.284 | 1.84E-05 | 5.60E-03 |
| Dmel_CG32082 | 143.362 | -1.278 | 0.299 | -4.279 | 1.87E-05 | 5.60E-03 |
| Dmel_CG8805 | 2531.220 | 0.658 | 0.154 | 4.278 | 1.89E-05 | 5.60E-03 |
| Dmel_CG6542 | 4210.347 | 0.628 | 0.148 | 4.254 | 2.10E-05 | 6.06E-03 |
| Dmel_CG34325 | 218.050 | -1.225 | 0.289 | -4.241 | 2.23E-05 | 6.16E-03 |
| Dmel_CG7050 | 174.496 | -1.607 | 0.379 | -4.239 | 2.25E-05 | 6.16E-03 |
| Dmel_CG7676 | 35.784 | -1.999 | 0.474 | -4.219 | 2.45E-05 | 6.28E-03 |
| Dmel_CG42854 | 30.574 | -2.400 | 0.569 | -4.218 | 2.47E-05 | 6.28E-03 |
| Dmel_CG13000 | 15.067 | -2.359 | 0.560 | -4.214 | 2.51E-05 | 6.28E-03 |
| Dmel_CG10726 | 2404.031 | 0.799 | 0.190 | 4.212 | 2.53E-05 | 6.28E-03 |
| Dmel_CG6178 | 2685.460 | 0.496 | 0.119 | 4.180 | 2.91E-05 | 6.96E-03 |
| Dmel_CG14193 | 12.461 | -2.423 | 0.580 | -4.179 | 2.93E-05 | 6.96E-03 |
| Dmel_CG43749 | 68.898 | -2.003 | 0.484 | -4.138 | 3.50E-05 | 8.10E-03 |
| Dmel_CG2191 | 17.249 | -2.812 | 0.680 | -4.134 | 3.56E-05 | 8.10E-03 |
| Dmel_CG8262 | 752.433 | 1.089 | 0.265 | 4.107 | 4.00E-05 | 8.92E-03 |
| Dmel_CG7178 | 1358.215 | -1.725 | 0.421 | -4.100 | 4.14E-05 | 9.03E-03 |
| Dmel_CG3297 | 3125.993 | 0.709 | 0.173 | 4.092 | 4.27E-05 | 9.14E-03 |
| Dmel_CG5252 | 4020.919 | 0.503 | 0.124 | 4.074 | 4.63E-05 | 9.53E-03 |
| Dmel_CG14808 | 573.015 | -1.527 | 0.375 | -4.070 | 4.69E-05 | 9.53E-03 |
| Dmel_CG6006 | 523.400 | -1.228 | 0.302 | -4.065 | 4.80E-05 | 9.53E-03 |
| Dmel_CG7660 | 20442.794 | 0.514 | 0.126 | 4.065 | 4.81E-05 | 9.53E-03 |
| Dmel_CG16995 | 24.742 | -1.770 | 0.436 | -4.056 | 5.00E-05 | 9.72E-03 |
| Dmel_CG2275 | 1215.191 | 0.896 | 0.222 | 4.036 | 5.43E-05 | 1.04E-02 |
| Dmel_CG18870 | 2820.357 | 0.576 | 0.143 | 4.030 | 5.57E-05 | 1.05E-02 |
| Dmel_CG32944 | 92.791 | -1.246 | 0.310 | -4.016 | 5.91E-05 | 1.09E-02 |

| | | | | | | |
|---|---|---|---|---|---|---|
| Dmel_CG3879 | 12.492 | -2.318 | 0.578 | -4.008 | 6.12E-05 | 1.09E-02 |
| Dmel_CG4476 | 2247.740 | 1.039 | 0.259 | 4.008 | 6.12E-05 | 1.09E-02 |
| Dmel_CG12582 | 1823.184 | 0.624 | 0.156 | 4.005 | 6.20E-05 | 1.09E-02 |
| Dmel_CG15814 | 1716.320 | 0.639 | 0.160 | 3.995 | 6.47E-05 | 1.11E-02 |
| Dmel_CG15173 | 91.496 | -1.131 | 0.284 | -3.985 | 6.73E-05 | 1.11E-02 |
| Dmel_CG9772 | 2989.847 | 0.632 | 0.159 | 3.980 | 6.90E-05 | 1.11E-02 |
| Dmel_CG18547 | 97.360 | -1.483 | 0.373 | -3.980 | 6.90E-05 | 1.11E-02 |
| Dmel_CG6698 | 17.300 | -3.148 | 0.792 | -3.976 | 7.00E-05 | 1.11E-02 |
| Dmel_CG18619 | 37.742 | -2.011 | 0.506 | -3.974 | 7.06E-05 | 1.11E-02 |
| Dmel_CG12002 | 217.218 | -1.718 | 0.432 | -3.974 | 7.08E-05 | 1.11E-02 |
| Dmel_CG3631 | 610.954 | 0.637 | 0.161 | 3.967 | 7.29E-05 | 1.13E-02 |
| Dmel_CG9650 | 454.888 | -1.001 | 0.255 | -3.933 | 8.40E-05 | 1.27E-02 |
| Dmel_CG9411 | 11.974 | -2.846 | 0.724 | -3.932 | 8.44E-05 | 1.27E-02 |
| Dmel_CG30101 | 46.664 | 2.134 | 0.544 | 3.923 | 8.75E-05 | 1.30E-02 |
| Dmel_CG30489 | 122.263 | -1.002 | 0.256 | -3.916 | 8.99E-05 | 1.32E-02 |
| Dmel_CG6477 | 1858.505 | 1.056 | 0.270 | 3.913 | 9.12E-05 | 1.32E-02 |
| Dmel_CG6324 | 34.299 | -1.701 | 0.436 | -3.903 | 9.51E-05 | 1.36E-02 |
| Dmel_CG31989 | 498.990 | 0.777 | 0.200 | 3.893 | 9.90E-05 | 1.39E-02 |
| Dmel_CG17078 | 3064.101 | 0.927 | 0.239 | 3.885 | 1.02E-04 | 1.41E-02 |
| Dmel_CG7084 | 38.686 | -2.379 | 0.613 | -3.884 | 1.03E-04 | 1.41E-02 |
| Dmel_CG17927 | 1300.359 | -1.575 | 0.406 | -3.877 | 1.06E-04 | 1.43E-02 |
| Dmel_CG18568 | 26.606 | 1.247 | 0.322 | 3.870 | 1.09E-04 | 1.44E-02 |
| Dmel_CG14680 | 22.113 | -1.898 | 0.491 | -3.867 | 1.10E-04 | 1.44E-02 |
| Dmel_CG8193 | 9.669 | -2.935 | 0.759 | -3.866 | 1.11E-04 | 1.44E-02 |
| Dmel_CG10706 | 358.379 | -1.147 | 0.298 | -3.848 | 1.19E-04 | 1.54E-02 |
| Dmel_CG42512 | 8.618 | -2.278 | 0.593 | -3.840 | 1.23E-04 | 1.57E-02 |
| Dmel_CG43088 | 13.320 | -2.541 | 0.663 | -3.834 | 1.26E-04 | 1.57E-02 |
| Dmel_CG8233 | 3636.327 | 0.540 | 0.141 | 3.834 | 1.26E-04 | 1.57E-02 |
| Dmel_CG10638 | 1166.946 | -0.838 | 0.219 | -3.826 | 1.30E-04 | 1.60E-02 |
| Dmel_CG17117 | 134.383 | -1.721 | 0.451 | -3.820 | 1.33E-04 | 1.62E-02 |
| Dmel_CR45530 | 16.389 | -1.740 | 0.456 | -3.813 | 1.37E-04 | 1.65E-02 |

| | | | | | | |
|---|---|---|---|---|---|---|
| Dmel_CG13954 | 10.767 | -1.844 | 0.484 | -3.808 | 1.40E-04 | 1.66E-02 |
| Dmel_CG16762 | 7.124 | -3.388 | 0.893 | -3.796 | 1.47E-04 | 1.71E-02 |
| Dmel_CG14796 | 11568.819 | 1.541 | 0.406 | 3.795 | 1.48E-04 | 1.71E-02 |
| Dmel_CG13928 | 13.709 | -1.897 | 0.500 | -3.793 | 1.49E-04 | 1.71E-02 |
| Dmel_CG18522 | 301.389 | -1.485 | 0.392 | -3.787 | 1.52E-04 | 1.73E-02 |
| Dmel_CG10521 | 195.366 | -1.332 | 0.352 | -3.784 | 1.54E-04 | 1.74E-02 |
| Dmel_CG12283 | 681.554 | -0.997 | 0.265 | -3.759 | 1.70E-04 | 1.89E-02 |
| Dmel_CG2893 | 2410.997 | 0.588 | 0.156 | 3.758 | 1.71E-04 | 1.89E-02 |
| Dmel_CG6479 | 2633.182 | 0.946 | 0.252 | 3.749 | 1.78E-04 | 1.92E-02 |
| Dmel_CR43461 | 15.553 | -1.690 | 0.451 | -3.748 | 1.78E-04 | 1.92E-02 |
| Dmel_CG33472 | 105.352 | -1.291 | 0.345 | -3.745 | 1.80E-04 | 1.93E-02 |
| Dmel_CG12449 | 143.489 | -1.377 | 0.369 | -3.730 | 1.91E-04 | 1.98E-02 |
| Dmel_CG9338 | 181.285 | -1.597 | 0.429 | -3.727 | 1.94E-04 | 1.98E-02 |
| Dmel_CG6958 | 2318.760 | 0.411 | 0.110 | 3.727 | 1.94E-04 | 1.98E-02 |
| Dmel_CG10553 | 15.343 | -3.397 | 0.912 | -3.726 | 1.95E-04 | 1.98E-02 |
| Dmel_CG4099 | 50.953 | -1.505 | 0.404 | -3.726 | 1.95E-04 | 1.98E-02 |
| Dmel_CG3812 | 1006.282 | 0.566 | 0.152 | 3.724 | 1.96E-04 | 1.98E-02 |
| Dmel_CG42486 | 7.794 | -1.777 | 0.478 | -3.716 | 2.02E-04 | 2.01E-02 |
| Dmel_CG32642 | 1743.930 | 2.684 | 0.723 | 3.713 | 2.05E-04 | 2.01E-02 |
| Dmel_CG42492 | 197.428 | -1.605 | 0.432 | -3.712 | 2.06E-04 | 2.01E-02 |
| Dmel_CG15324 | 366.537 | 1.612 | 0.434 | 3.711 | 2.06E-04 | 2.01E-02 |
| Dmel_CG9610 | 35.323 | -1.695 | 0.458 | -3.704 | 2.12E-04 | 2.03E-02 |
| Dmel_CG10795 | 653.543 | 1.082 | 0.292 | 3.703 | 2.13E-04 | 2.03E-02 |
| Dmel_CG5338 | 70.237 | -1.831 | 0.495 | -3.699 | 2.16E-04 | 2.04E-02 |
| Dmel_CG14615 | 327.742 | 0.786 | 0.213 | 3.696 | 2.19E-04 | 2.04E-02 |
| Dmel_CG6422 | 4864.193 | 0.631 | 0.171 | 3.692 | 2.22E-04 | 2.04E-02 |
| Dmel_CG10550 | 16.054 | -2.006 | 0.543 | -3.692 | 2.23E-04 | 2.04E-02 |
| Dmel_CG32364 | 249.152 | -1.255 | 0.340 | -3.690 | 2.24E-04 | 2.04E-02 |
| Dmel_CG2857 | 23.133 | -1.953 | 0.530 | -3.688 | 2.26E-04 | 2.04E-02 |
| Dmel_CG12375 | 831.121 | 0.541 | 0.147 | 3.684 | 2.29E-04 | 2.04E-02 |

| | | | | | | |
|---|---|---|---|---|---|---|
| Dmel_CG11125 | 345.793 | 0.825 | 0.224 | 3.683 | 2.30E-04 | 2.04E-02 |
| Dmel_CG32577 | 132.606 | -1.219 | 0.331 | -3.683 | 2.30E-04 | 2.04E-02 |
| Dmel_CR44472 | 23.272 | 1.829 | 0.497 | 3.679 | 2.34E-04 | 2.05E-02 |
| Dmel_CR43144 | 234.567 | -1.219 | 0.332 | -3.674 | 2.39E-04 | 2.08E-02 |
| Dmel_CG31901 | 28.374 | -2.490 | 0.680 | -3.664 | 2.49E-04 | 2.15E-02 |
| Dmel_CG2246 | 2660.559 | 0.526 | 0.144 | 3.658 | 2.54E-04 | 2.15E-02 |
| Dmel_CG41106 | 15.890 | -1.904 | 0.521 | -3.657 | 2.55E-04 | 2.15E-02 |
| Dmel_CG9707 | 1943.581 | 0.589 | 0.161 | 3.649 | 2.63E-04 | 2.15E-02 |
| Dmel_CG32019 | 11590.256 | -1.412 | 0.387 | -3.649 | 2.64E-04 | 2.15E-02 |
| Dmel_CG7510 | 2088.312 | 0.558 | 0.153 | 3.648 | 2.64E-04 | 2.15E-02 |
| Dmel_CG5041 | 570.781 | 0.710 | 0.195 | 3.648 | 2.64E-04 | 2.15E-02 |
| Dmel_CG12423 | 221.162 | -1.190 | 0.326 | -3.647 | 2.65E-04 | 2.15E-02 |
| Dmel_CG10336 | 664.150 | 0.885 | 0.243 | 3.646 | 2.66E-04 | 2.15E-02 |
| Dmel_CG18039 | 57.006 | -1.759 | 0.483 | -3.644 | 2.69E-04 | 2.15E-02 |
| Dmel_CG7564 | 14794.411 | -1.158 | 0.318 | -3.643 | 2.70E-04 | 2.15E-02 |
| Dmel_CG42639 | 16.055 | -2.624 | 0.721 | -3.640 | 2.73E-04 | 2.16E-02 |
| Dmel_CG15432 | 193.546 | -1.233 | 0.339 | -3.635 | 2.77E-04 | 2.18E-02 |
| Dmel_CG45077 | 281.510 | -1.509 | 0.416 | -3.627 | 2.86E-04 | 2.23E-02 |
| Dmel_CG13364 | 597.882 | -1.546 | 0.427 | -3.624 | 2.90E-04 | 2.23E-02 |
| Dmel_CR34084 | 11.680 | -2.032 | 0.561 | -3.621 | 2.93E-04 | 2.23E-02 |
| Dmel_CG46462 | 29.749 | -1.810 | 0.500 | -3.621 | 2.94E-04 | 2.23E-02 |
| Dmel_CG10794 | 23.361 | -4.266 | 1.179 | -3.617 | 2.98E-04 | 2.23E-02 |
| Dmel_CG33474 | 26.760 | -1.326 | 0.367 | -3.616 | 2.99E-04 | 2.23E-02 |
| Dmel_CR45187 | 109.321 | -1.218 | 0.337 | -3.616 | 2.99E-04 | 2.23E-02 |
| Dmel_CG32677 | 64.186 | -1.706 | 0.472 | -3.615 | 3.01E-04 | 2.23E-02 |
| Dmel_CG15155 | 16.725 | -1.854 | 0.513 | -3.612 | 3.03E-04 | 2.24E-02 |
| Dmel_CG31198 | 57.349 | -5.117 | 1.419 | -3.605 | 3.12E-04 | 2.29E-02 |
| Dmel_CG13604 | 682.846 | 0.752 | 0.209 | 3.601 | 3.17E-04 | 2.29E-02 |
| Dmel_CG31897 | 145.916 | -0.865 | 0.241 | -3.596 | 3.24E-04 | 2.29E-02 |

| | | | | | | |
|---|---|---|---|---|---|---|
| Dmel_CG32474 | 120.332 | -1.621 | 0.451 | -3.592 | 3.29E-04 | 2.29E-02 |
| Dmel_CG4909 | 2833.384 | 0.845 | 0.235 | 3.590 | 3.30E-04 | 2.29E-02 |
| Dmel_CG5596 | 653.286 | -1.482 | 0.413 | -3.590 | 3.30E-04 | 2.29E-02 |
| Dmel_CG17716 | 19.533 | -2.076 | 0.578 | -3.590 | 3.31E-04 | 2.29E-02 |
| Dmel_CG32017 | 173.619 | -1.268 | 0.353 | -3.589 | 3.32E-04 | 2.29E-02 |
| Dmel_CG5867 | 86.914 | -1.403 | 0.391 | -3.588 | 3.33E-04 | 2.29E-02 |
| Dmel_CG42309 | 229.779 | -1.694 | 0.472 | -3.587 | 3.35E-04 | 2.29E-02 |
| Dmel_CG7128 | 682.992 | 1.249 | 0.348 | 3.586 | 3.35E-04 | 2.29E-02 |
| Dmel_CG43758 | 309.775 | -1.157 | 0.323 | -3.586 | 3.36E-04 | 2.29E-02 |
| Dmel_CG5445 | 1662.056 | 0.563 | 0.157 | 3.579 | 3.46E-04 | 2.33E-02 |
| Dmel_CG3613 | 1120.960 | 0.860 | 0.240 | 3.577 | 3.47E-04 | 2.33E-02 |
| Dmel_CG10364 | 1000.208 | 1.068 | 0.299 | 3.575 | 3.50E-04 | 2.33E-02 |
| Dmel_CG1024 | 621.198 | 0.818 | 0.229 | 3.574 | 3.51E-04 | 2.33E-02 |
| Dmel_CG8165 | 279.924 | 0.513 | 0.144 | 3.572 | 3.54E-04 | 2.34E-02 |
| Dmel_CG8585 | 391.292 | -1.143 | 0.320 | -3.569 | 3.58E-04 | 2.35E-02 |
| Dmel_CG3171 | 1422.762 | 0.794 | 0.223 | 3.561 | 3.69E-04 | 2.41E-02 |
| Dmel_CG18536 | 49.200 | -1.180 | 0.332 | -3.556 | 3.77E-04 | 2.44E-02 |
| Dmel_CG5038 | 341.144 | 0.614 | 0.173 | 3.554 | 3.80E-04 | 2.45E-02 |
| Dmel_CG7083 | 1256.892 | 0.546 | 0.154 | 3.550 | 3.85E-04 | 2.46E-02 |
| Dmel_CG34445 | 67.998 | -1.639 | 0.462 | -3.549 | 3.87E-04 | 2.46E-02 |
| Dmel_CG9297 | 38.481 | -1.620 | 0.458 | -3.539 | 4.01E-04 | 2.54E-02 |
| Dmel_CG15443 | 379.439 | 0.676 | 0.191 | 3.537 | 4.05E-04 | 2.55E-02 |
| Dmel_CG1825 | 2353.709 | 1.329 | 0.376 | 3.532 | 4.13E-04 | 2.56E-02 |
| Dmel_CG43664 | 825.006 | 0.783 | 0.222 | 3.531 | 4.14E-04 | 2.56E-02 |
| Dmel_CR46482 | 19457.346 | -1.042 | 0.295 | -3.531 | 4.14E-04 | 2.56E-02 |
| Dmel_CG13977 | 13.266 | -2.838 | 0.804 | -3.529 | 4.16E-04 | 2.56E-02 |
| Dmel_CG4608 | 113.986 | -1.087 | 0.308 | -3.525 | 4.24E-04 | 2.59E-02 |
| Dmel_CG30195 | 12.984 | -2.196 | 0.623 | -3.523 | 4.26E-04 | 2.59E-02 |
| Dmel_CG14026 | 3581.522 | 0.351 | 0.100 | 3.521 | 4.30E-04 | 2.60E-02 |
| Dmel_CG12106 | 309.494 | 0.865 | 0.246 | 3.515 | 4.40E-04 | 2.61E-02 |

| | | | | | | |
|---|---|---|---|---|---|---|
| Dmel_CG2184 | 1690.958 | -1.347 | 0.383 | -3.514 | 4.41E-04 | 2.61E-02 |
| Dmel_CR33628 | 2075.290 | -2.268 | 0.646 | -3.512 | 4.45E-04 | 2.61E-02 |
| Dmel_CR33921 | 2075.290 | -2.268 | 0.646 | -3.512 | 4.45E-04 | 2.61E-02 |
| Dmel_CG7997 | 780.900 | 0.535 | 0.152 | 3.511 | 4.46E-04 | 2.61E-02 |
| Dmel_CG1455 | 14.264 | -1.917 | 0.546 | -3.511 | 4.46E-04 | 2.61E-02 |
| Dmel_CG9796 | 5516.729 | 0.623 | 0.178 | 3.503 | 4.60E-04 | 2.67E-02 |
| Dmel_CG9057 | 16761.326 | 1.152 | 0.329 | 3.502 | 4.62E-04 | 2.67E-02 |
| Dmel_CG15138 | 159.816 | -1.071 | 0.306 | -3.498 | 4.68E-04 | 2.68E-02 |
| Dmel_CG18102 | 3933.133 | 0.390 | 0.112 | 3.498 | 4.69E-04 | 2.68E-02 |
| Dmel_CG44007 | 82.476 | -1.583 | 0.453 | -3.494 | 4.76E-04 | 2.71E-02 |
| Dmel_CG9423 | 10906.330 | 0.593 | 0.170 | 3.489 | 4.85E-04 | 2.73E-02 |
| Dmel_CG14711 | 1052.838 | 0.629 | 0.180 | 3.488 | 4.86E-04 | 2.73E-02 |
| Dmel_CG6957 | 112.117 | -1.274 | 0.366 | -3.485 | 4.92E-04 | 2.75E-02 |
| Dmel_CG6202 | 3774.453 | 0.437 | 0.125 | 3.484 | 4.95E-04 | 2.75E-02 |
| Dmel_CG12099 | 2772.921 | 0.627 | 0.180 | 3.483 | 4.96E-04 | 2.75E-02 |
| Dmel_CG6927 | 4422.478 | 0.656 | 0.189 | 3.479 | 5.04E-04 | 2.76E-02 |
| Dmel_CG1894 | 11.141 | -1.881 | 0.541 | -3.479 | 5.04E-04 | 2.76E-02 |
| Dmel_CG5439 | 666.759 | 0.731 | 0.210 | 3.475 | 5.11E-04 | 2.78E-02 |
| Dmel_CG3407 | 971.487 | 0.648 | 0.187 | 3.473 | 5.15E-04 | 2.78E-02 |
| Dmel_CG12220 | 492.880 | -1.064 | 0.306 | -3.473 | 5.16E-04 | 2.78E-02 |
| Dmel_CG4622 | 759.225 | 0.889 | 0.256 | 3.471 | 5.18E-04 | 2.78E-02 |
| Dmel_CG6658 | 157.093 | -1.473 | 0.425 | -3.464 | 5.32E-04 | 2.85E-02 |
| Dmel_CG18549 | 1584.276 | 0.459 | 0.133 | 3.462 | 5.35E-04 | 2.85E-02 |
| Dmel_CG6930 | 714.992 | -0.856 | 0.247 | -3.459 | 5.41E-04 | 2.87E-02 |
| Dmel_CG2072 | 1813.889 | 0.512 | 0.148 | 3.457 | 5.47E-04 | 2.87E-02 |
| Dmel_CG8023 | 8.117 | -2.756 | 0.798 | -3.454 | 5.51E-04 | 2.87E-02 |
| Dmel_CG3163 | 236.299 | 0.760 | 0.220 | 3.454 | 5.52E-04 | 2.87E-02 |
| Dmel_CG12110 | 4136.182 | 0.530 | 0.154 | 3.451 | 5.58E-04 | 2.87E-02 |
| Dmel_CG32641 | 631.055 | -0.993 | 0.288 | -3.450 | 5.60E-04 | 2.87E-02 |

| | | | | | | |
|---|---|---|---|---|---|---|
| Dmel_CG6357 | 882.290 | -0.763 | 0.221 | -3.450 | 5.60E-04 | 2.87E-02 |
| Dmel_CG9155 | 396.786 | -0.909 | 0.264 | -3.450 | 5.61E-04 | 2.87E-02 |
| Dmel_CG33344 | 7.747 | -2.076 | 0.602 | -3.449 | 5.63E-04 | 2.87E-02 |
| Dmel_CG8316 | 213.397 | -0.967 | 0.281 | -3.447 | 5.67E-04 | 2.87E-02 |
| Dmel_CG6665 | 468.601 | 0.565 | 0.164 | 3.444 | 5.74E-04 | 2.89E-02 |
| Dmel_CG6040 | 548.589 | -0.925 | 0.269 | -3.441 | 5.81E-04 | 2.92E-02 |
| Dmel_CG4274 | 2581.259 | 1.038 | 0.302 | 3.438 | 5.86E-04 | 2.93E-02 |
| Dmel_CG7957 | 723.957 | 0.726 | 0.211 | 3.436 | 5.90E-04 | 2.93E-02 |
| Dmel_CG4395 | 13.227 | 2.039 | 0.594 | 3.432 | 5.99E-04 | 2.93E-02 |
| Dmel_CG13941 | 124.885 | -0.800 | 0.233 | -3.431 | 6.02E-04 | 2.93E-02 |
| Dmel_CG6121 | 978.359 | 0.918 | 0.268 | 3.430 | 6.04E-04 | 2.93E-02 |
| Dmel_CG4951 | 1445.689 | 0.530 | 0.155 | 3.428 | 6.09E-04 | 2.93E-02 |
| Dmel_CG34200 | 556.763 | -1.353 | 0.395 | -3.426 | 6.12E-04 | 2.93E-02 |
| Dmel_CR45132 | 40.764 | -0.987 | 0.288 | -3.426 | 6.14E-04 | 2.93E-02 |
| Dmel_CG7289 | 1206.822 | 0.653 | 0.191 | 3.425 | 6.15E-04 | 2.93E-02 |
| Dmel_CG7157 | 83.139 | -1.164 | 0.340 | -3.424 | 6.17E-04 | 2.93E-02 |
| Dmel_CG11737 | 998.601 | 0.956 | 0.279 | 3.423 | 6.18E-04 | 2.93E-02 |
| Dmel_CG6398 | 1931.984 | 0.944 | 0.276 | 3.423 | 6.18E-04 | 2.93E-02 |
| Dmel_CG4686 | 727.346 | 0.684 | 0.200 | 3.423 | 6.19E-04 | 2.93E-02 |
| Dmel_CG18642 | 441.573 | 0.918 | 0.268 | 3.421 | 6.25E-04 | 2.94E-02 |
| Dmel_CG15201 | 9.880 | -2.006 | 0.587 | -3.420 | 6.27E-04 | 2.94E-02 |
| Dmel_CG11210 | 1947.088 | 0.651 | 0.191 | 3.417 | 6.33E-04 | 2.94E-02 |
| Dmel_CG1664 | 3508.885 | 0.707 | 0.207 | 3.417 | 6.34E-04 | 2.94E-02 |
| Dmel_CG6962 | 1700.725 | 0.814 | 0.239 | 3.414 | 6.40E-04 | 2.94E-02 |
| Dmel_CG10825 | 890.498 | 0.655 | 0.192 | 3.414 | 6.41E-04 | 2.94E-02 |
| Dmel_CG13506 | 177.409 | -1.083 | 0.317 | -3.413 | 6.41E-04 | 2.94E-02 |
| Dmel_CG34392 | 202.441 | 1.421 | 0.416 | 3.413 | 6.43E-04 | 2.94E-02 |
| Dmel_CG4620 | 4335.134 | 0.639 | 0.187 | 3.411 | 6.47E-04 | 2.94E-02 |
| Dmel_CG44325 | 724.990 | -0.784 | 0.230 | -3.405 | 6.62E-04 | 2.99E-02 |
| Dmel_CG14946 | 22.832 | -1.745 | 0.513 | -3.403 | 6.66E-04 | 2.99E-02 |
| Dmel_CG17084 | 133.181 | -1.112 | 0.327 | -3.403 | 6.66E-04 | 2.99E-02 |

| | | | | | | |
|---|---|---|---|---|---|---|
| Dmel_CG41265 | 266.240 | -1.196 | 0.352 | -3.398 | 6.78E-04 | 3.03E-02 |
| Dmel_CG17149 | 1566.784 | 0.898 | 0.264 | 3.396 | 6.83E-04 | 3.04E-02 |
| Dmel_CG7772 | 674.473 | 0.756 | 0.223 | 3.393 | 6.91E-04 | 3.04E-02 |
| Dmel_CG4260 | 7777.907 | 0.482 | 0.142 | 3.393 | 6.91E-04 | 3.04E-02 |
| Dmel_CG32115 | 14.590 | -1.894 | 0.558 | -3.393 | 6.91E-04 | 3.04E-02 |
| Dmel_CR42451 | 792.575 | -1.170 | 0.345 | -3.391 | 6.96E-04 | 3.04E-02 |
| Dmel_CG30170 | 20.719 | -1.855 | 0.547 | -3.391 | 6.96E-04 | 3.04E-02 |
| Dmel_CG4262 | 252.039 | -0.841 | 0.248 | -3.387 | 7.07E-04 | 3.07E-02 |
| Dmel_CG34133 | 4321.118 | 0.566 | 0.167 | 3.383 | 7.18E-04 | 3.11E-02 |
| Dmel_CG12942 | 1863.699 | 0.587 | 0.174 | 3.381 | 7.23E-04 | 3.12E-02 |
| Dmel_CG5725 | 1904.398 | 0.734 | 0.217 | 3.377 | 7.33E-04 | 3.14E-02 |
| Dmel_CG1915 | 6242.146 | -1.045 | 0.310 | -3.376 | 7.34E-04 | 3.14E-02 |
| Dmel_CG33147 | 9.630 | -2.113 | 0.626 | -3.374 | 7.40E-04 | 3.15E-02 |
| Dmel_CG32506 | 9.534 | -2.114 | 0.627 | -3.372 | 7.46E-04 | 3.16E-02 |
| Dmel_CG4267 | 559.666 | 0.880 | 0.261 | 3.371 | 7.48E-04 | 3.16E-02 |
| Dmel_CG12391 | 1629.747 | 0.738 | 0.219 | 3.367 | 7.61E-04 | 3.20E-02 |
| Dmel_CG10309 | 2135.421 | 1.285 | 0.382 | 3.366 | 7.62E-04 | 3.20E-02 |
| Dmel_CG15848 | 101.067 | -1.331 | 0.396 | -3.364 | 7.68E-04 | 3.21E-02 |
| Dmel_CG1911 | 1431.817 | 0.860 | 0.256 | 3.359 | 7.81E-04 | 3.25E-02 |
| Dmel_CG8051 | 10.915 | -1.571 | 0.468 | -3.354 | 7.96E-04 | 3.30E-02 |
| Dmel_CG33103 | 1003.050 | -1.419 | 0.424 | -3.351 | 8.04E-04 | 3.32E-02 |
| Dmel_CG6207 | 2745.065 | 0.708 | 0.211 | 3.350 | 8.07E-04 | 3.32E-02 |
| Dmel_CG32452 | 1457.804 | 0.813 | 0.243 | 3.349 | 8.12E-04 | 3.33E-02 |
| Dmel_CG10240 | 935.082 | 0.709 | 0.212 | 3.347 | 8.18E-04 | 3.34E-02 |
| Dmel_CR43626 | 142.895 | -1.241 | 0.371 | -3.346 | 8.20E-04 | 3.34E-02 |
| Dmel_CG5222 | 595.645 | 0.896 | 0.268 | 3.344 | 8.27E-04 | 3.35E-02 |
| Dmel_CG17462 | 1947.213 | 0.930 | 0.279 | 3.337 | 8.46E-04 | 3.39E-02 |
| Dmel_CG2765 | 1659.825 | 0.624 | 0.187 | 3.337 | 8.48E-04 | 3.39E-02 |
| Dmel_CG4029 | 5627.255 | 0.658 | 0.197 | 3.336 | 8.50E-04 | 3.39E-02 |
| Dmel_CG12118 | 916.548 | 0.632 | 0.189 | 3.336 | 8.51E-04 | 3.39E-02 |
| Dmel_CG4978 | 3421.524 | 0.862 | 0.259 | 3.334 | 8.55E-04 | 3.39E-02 |

| | | | | | | |
|---|---|---|---|---|---|---|
| Dmel_CG4821 | 225.884 | -0.994 | 0.298 | -3.332 | 8.62E-04 | 3.39E-02 |
| Dmel_CG3041 | 913.876 | 0.915 | 0.275 | 3.332 | 8.63E-04 | 3.39E-02 |
| Dmel_CG9712 | 1406.203 | 0.729 | 0.219 | 3.331 | 8.64E-04 | 3.39E-02 |
| Dmel_CG11975 | 684.605 | 0.676 | 0.203 | 3.331 | 8.66E-04 | 3.39E-02 |
| Dmel_CG44008 | 16.277 | -1.804 | 0.542 | -3.329 | 8.70E-04 | 3.40E-02 |
| Dmel_CG6536 | 115.126 | -1.090 | 0.328 | -3.325 | 8.84E-04 | 3.43E-02 |
| Dmel_CG11367 | 593.483 | 1.063 | 0.320 | 3.325 | 8.86E-04 | 3.43E-02 |
| Dmel_CG17839 | 101.907 | -1.084 | 0.326 | -3.324 | 8.88E-04 | 3.43E-02 |
| Dmel_CG8282 | 2013.871 | 0.529 | 0.159 | 3.322 | 8.94E-04 | 3.44E-02 |
| Dmel_CG11804 | 8263.425 | 0.502 | 0.151 | 3.320 | 9.00E-04 | 3.44E-02 |
| Dmel_CG8389 | 831.413 | 0.533 | 0.161 | 3.318 | 9.07E-04 | 3.44E-02 |
| Dmel_CG9470 | 1461.555 | -1.099 | 0.331 | -3.317 | 9.09E-04 | 3.44E-02 |
| Dmel_CG42253 | 66.402 | -1.475 | 0.445 | -3.316 | 9.14E-04 | 3.44E-02 |
| Dmel_CG1674 | 540.658 | -1.334 | 0.402 | -3.316 | 9.14E-04 | 3.44E-02 |
| Dmel_CG8411 | 7127.715 | 0.438 | 0.132 | 3.315 | 9.16E-04 | 3.44E-02 |
| Dmel_CG5177 | 88.185 | -1.824 | 0.550 | -3.315 | 9.18E-04 | 3.44E-02 |
| Dmel_CG34438 | 3017.459 | 0.722 | 0.218 | 3.311 | 9.29E-04 | 3.47E-02 |
| Dmel_CG5602 | 1070.389 | 0.892 | 0.269 | 3.310 | 9.34E-04 | 3.48E-02 |
| Dmel_CG5807 | 2244.113 | 0.476 | 0.144 | 3.308 | 9.39E-04 | 3.49E-02 |
| Dmel_CG7107 | 460.078 | -1.355 | 0.410 | -3.303 | 9.56E-04 | 3.54E-02 |
| Dmel_CG10895 | 11736.045 | 0.416 | 0.126 | 3.302 | 9.61E-04 | 3.54E-02 |
| Dmel_CG7404 | 1099.831 | 0.354 | 0.107 | 3.299 | 9.69E-04 | 3.56E-02 |
| Dmel_CG8994 | 15778.334 | 0.610 | 0.185 | 3.297 | 9.79E-04 | 3.58E-02 |
| Dmel_CR43314 | 390.005 | -1.144 | 0.347 | -3.294 | 9.88E-04 | 3.61E-02 |
| Dmel_CG10126 | 206.811 | -1.209 | 0.367 | -3.293 | 9.91E-04 | 3.61E-02 |
| Dmel_CG34431 | 161.743 | -0.853 | 0.259 | -3.291 | 9.99E-04 | 3.62E-02 |
| Dmel_CG9203 | 680.819 | 0.559 | 0.170 | 3.290 | 1.00E-03 | 3.62E-02 |
| Dmel_CG9739 | 548.694 | -1.009 | 0.307 | -3.288 | 1.01E-03 | 3.63E-02 |
| Dmel_CG31469 | 13.182 | -1.602 | 0.487 | -3.287 | 1.01E-03 | 3.63E-02 |

| | | | | | | |
|---|---|---|---|---|---|---|
| Dmel_CG9594 | 2785.003 | 0.659 | 0.201 | 3.285 | 1.02E-03 | 3.65E-02 |
| Dmel_CG12004 | 4534.896 | 0.695 | 0.212 | 3.284 | 1.03E-03 | 3.65E-02 |
| Dmel_CG9432 | 408.363 | -1.247 | 0.380 | -3.283 | 1.03E-03 | 3.65E-02 |
| Dmel_CG4405 | 155.934 | -1.333 | 0.406 | -3.281 | 1.03E-03 | 3.66E-02 |
| Dmel_CG13403 | 9.694 | -2.267 | 0.691 | -3.279 | 1.04E-03 | 3.68E-02 |
| Dmel_CG4788 | 695.972 | 0.809 | 0.247 | 3.276 | 1.05E-03 | 3.70E-02 |
| Dmel_CG42259 | 266.650 | -1.368 | 0.418 | -3.273 | 1.06E-03 | 3.72E-02 |
| Dmel_CG1019 | 287.029 | -0.997 | 0.305 | -3.273 | 1.07E-03 | 3.72E-02 |
| Dmel_CG4615 | 903.993 | 0.844 | 0.258 | 3.268 | 1.08E-03 | 3.78E-02 |
| Dmel_CG3651 | 2573.552 | 0.462 | 0.141 | 3.266 | 1.09E-03 | 3.79E-02 |
| Dmel_CG2990 | 1142.823 | 0.655 | 0.201 | 3.265 | 1.10E-03 | 3.79E-02 |
| Dmel_CG5180 | 943.481 | 1.088 | 0.333 | 3.264 | 1.10E-03 | 3.79E-02 |
| Dmel_CG8440 | 4400.552 | 0.398 | 0.122 | 3.263 | 1.10E-03 | 3.80E-02 |
| Dmel_CR46254 | 14.144 | -1.647 | 0.505 | -3.261 | 1.11E-03 | 3.81E-02 |
| Dmel_CG9123 | 285.710 | 0.970 | 0.298 | 3.259 | 1.12E-03 | 3.82E-02 |
| Dmel_CG32758 | 1103.252 | -0.563 | 0.173 | -3.254 | 1.14E-03 | 3.88E-02 |
| Dmel_CG30418 | 100.457 | -1.346 | 0.414 | -3.247 | 1.16E-03 | 3.94E-02 |
| Dmel_CG31926 | 1397.870 | 2.466 | 0.759 | 3.247 | 1.17E-03 | 3.94E-02 |
| Dmel_CG4139 | 67.549 | -1.192 | 0.368 | -3.243 | 1.18E-03 | 3.96E-02 |
| Dmel_CG1109 | 2597.079 | 0.493 | 0.152 | 3.243 | 1.18E-03 | 3.96E-02 |
| Dmel_CR45941 | 8.758 | -2.186 | 0.674 | -3.243 | 1.18E-03 | 3.96E-02 |
| Dmel_CG11979 | 478.991 | -1.044 | 0.322 | -3.240 | 1.19E-03 | 3.96E-02 |
| Dmel_CG33556 | 69.075 | -1.477 | 0.456 | -3.239 | 1.20E-03 | 3.96E-02 |
| Dmel_CG8400 | 3643.172 | 0.672 | 0.207 | 3.239 | 1.20E-03 | 3.96E-02 |
| Dmel_CG33720 | 267.591 | -1.014 | 0.313 | -3.237 | 1.21E-03 | 3.96E-02 |
| Dmel_CG34250 | 185.219 | -1.312 | 0.405 | -3.237 | 1.21E-03 | 3.96E-02 |
| Dmel_CG10387 | 2509.834 | 0.619 | 0.191 | 3.237 | 1.21E-03 | 3.96E-02 |
| Dmel_CR44953 | 22.110 | -1.516 | 0.468 | -3.237 | 1.21E-03 | 3.96E-02 |
| Dmel_CG6711 | 1884.750 | 0.427 | 0.132 | 3.236 | 1.21E-03 | 3.96E-02 |
| Dmel_CG5938 | 1070.366 | 0.735 | 0.227 | 3.234 | 1.22E-03 | 3.98E-02 |
| Dmel_CG12275 | 39.817 | -1.912 | 0.592 | -3.230 | 1.24E-03 | 4.03E-02 |

| | | | | | | |
|---|---|---|---|---|---|---|
| Dmel_CG16947 | 157.907 | 1.077 | 0.333 | 3.229 | 1.24E-03 | 4.03E-02 |
| Dmel_CG17292 | 1474.083 | 0.641 | 0.199 | 3.224 | 1.26E-03 | 4.06E-02 |
| Dmel_CG43273 | 105.193 | -0.861 | 0.267 | -3.224 | 1.27E-03 | 4.06E-02 |
| Dmel_CG32096 | 237.586 | -0.892 | 0.277 | -3.223 | 1.27E-03 | 4.06E-02 |
| Dmel_CG6329 | 64.231 | -1.629 | 0.505 | -3.223 | 1.27E-03 | 4.06E-02 |
| Dmel_CG31753 | 71.034 | -1.431 | 0.445 | -3.216 | 1.30E-03 | 4.16E-02 |
| Dmel_CG4433 | 1942.289 | 0.732 | 0.228 | 3.213 | 1.31E-03 | 4.18E-02 |
| Dmel_CG42584 | 108.987 | -1.531 | 0.477 | -3.212 | 1.32E-03 | 4.18E-02 |
| Dmel_CG4145 | 1028.805 | -1.141 | 0.355 | -3.212 | 1.32E-03 | 4.18E-02 |
| Dmel_CG10063 | 16.237 | -1.568 | 0.488 | -3.209 | 1.33E-03 | 4.19E-02 |
| Dmel_CG11098 | 1467.654 | -0.416 | 0.130 | -3.209 | 1.33E-03 | 4.19E-02 |
| Dmel_CG32795 | 1510.068 | 0.441 | 0.137 | 3.208 | 1.34E-03 | 4.19E-02 |
| Dmel_CG15435 | 899.900 | 0.762 | 0.238 | 3.207 | 1.34E-03 | 4.20E-02 |
| Dmel_CG10570 | 76.217 | -1.661 | 0.518 | -3.203 | 1.36E-03 | 4.23E-02 |
| Dmel_CG42502 | 76.217 | -1.661 | 0.518 | -3.203 | 1.36E-03 | 4.23E-02 |
| Dmel_CR44042 | 410.787 | -0.827 | 0.259 | -3.196 | 1.40E-03 | 4.33E-02 |
| Dmel_CG17360 | 1450.158 | 0.628 | 0.197 | 3.193 | 1.41E-03 | 4.36E-02 |
| Dmel_CG3348 | 150.597 | -1.169 | 0.366 | -3.191 | 1.42E-03 | 4.36E-02 |
| Dmel_CG11007 | 1087.406 | 0.493 | 0.154 | 3.190 | 1.42E-03 | 4.36E-02 |
| Dmel_CG5905 | 30.547 | -1.470 | 0.461 | -3.190 | 1.43E-03 | 4.36E-02 |
| Dmel_CG5181 | 242.756 | 1.037 | 0.325 | 3.189 | 1.43E-03 | 4.36E-02 |
| Dmel_CG9046 | 18333.540 | 1.340 | 0.420 | 3.189 | 1.43E-03 | 4.36E-02 |
| Dmel_CG9836 | 704.353 | -0.686 | 0.215 | -3.183 | 1.46E-03 | 4.42E-02 |
| Dmel_CG12093 | 455.447 | 0.752 | 0.236 | 3.182 | 1.46E-03 | 4.42E-02 |
| Dmel_CG2993 | 277.855 | 0.974 | 0.306 | 3.181 | 1.47E-03 | 4.42E-02 |
| Dmel_CG10396 | 15.080 | -1.532 | 0.482 | -3.181 | 1.47E-03 | 4.42E-02 |
| Dmel_CG9984 | 1897.020 | 0.676 | 0.213 | 3.180 | 1.47E-03 | 4.42E-02 |
| Dmel_CG3403 | 2318.502 | 0.939 | 0.296 | 3.177 | 1.49E-03 | 4.45E-02 |
| Dmel_CG31807 | 141.606 | -0.932 | 0.293 | -3.176 | 1.49E-03 | 4.45E-02 |
| Dmel_CG46339 | 356.917 | -0.968 | 0.305 | -3.175 | 1.50E-03 | 4.45E-02 |

| | | | | | | |
|---|---|---|---|---|---|---|
| Dmel_CG7272 | 93.577 | -0.737 | 0.232 | -3.174 | 1.50E-03 | 4.45E-02 |
| Dmel_CG5939 | 354.903 | -1.070 | 0.337 | -3.174 | 1.50E-03 | 4.45E-02 |
| Dmel_CG10630 | 13.277 | -2.459 | 0.775 | -3.173 | 1.51E-03 | 4.45E-02 |
| Dmel_CG11723 | 1410.373 | 0.585 | 0.184 | 3.172 | 1.51E-03 | 4.45E-02 |
| Dmel_CR45897 | 10.683 | -1.749 | 0.551 | -3.171 | 1.52E-03 | 4.45E-02 |
| Dmel_CG7930 | 574.112 | -1.313 | 0.414 | -3.171 | 1.52E-03 | 4.45E-02 |
| Dmel_CG31365 | 1258.248 | 0.797 | 0.251 | 3.170 | 1.52E-03 | 4.45E-02 |
| Dmel_CG13434 | 94.002 | -0.939 | 0.297 | -3.166 | 1.55E-03 | 4.50E-02 |
| Dmel_CG42599 | 266.112 | -1.010 | 0.319 | -3.165 | 1.55E-03 | 4.50E-02 |
| Dmel_CG7837 | 1051.188 | 0.740 | 0.234 | 3.164 | 1.56E-03 | 4.52E-02 |
| Dmel_CG8811 | 8990.635 | 0.483 | 0.153 | 3.161 | 1.57E-03 | 4.53E-02 |
| Dmel_CG3836 | 2301.302 | 0.722 | 0.228 | 3.161 | 1.57E-03 | 4.53E-02 |
| Dmel_CG2330 | 77.759 | -1.493 | 0.472 | -3.160 | 1.58E-03 | 4.54E-02 |
| Dmel_CG44246 | 646.353 | 0.700 | 0.222 | 3.158 | 1.59E-03 | 4.55E-02 |
| Dmel_CG15721 | 6051.069 | 1.209 | 0.383 | 3.155 | 1.61E-03 | 4.59E-02 |
| Dmel_CG18507 | 93.141 | 1.199 | 0.380 | 3.154 | 1.61E-03 | 4.59E-02 |
| Dmel_CG32212 | 22.799 | -1.623 | 0.515 | -3.152 | 1.62E-03 | 4.60E-02 |
| Dmel_CG32320 | 96.491 | -1.370 | 0.435 | -3.152 | 1.62E-03 | 4.60E-02 |
| Dmel_CR44291 | 23.840 | -1.339 | 0.425 | -3.150 | 1.63E-03 | 4.62E-02 |
| Dmel_CG8933 | 5203.559 | 0.513 | 0.163 | 3.145 | 1.66E-03 | 4.68E-02 |
| Dmel_CG9379 | 238.648 | -0.827 | 0.263 | -3.145 | 1.66E-03 | 4.68E-02 |
| Dmel_CG3065 | 790.124 | 1.218 | 0.387 | 3.144 | 1.67E-03 | 4.68E-02 |
| Dmel_CG4214 | 1980.475 | 0.474 | 0.151 | 3.139 | 1.70E-03 | 4.75E-02 |
| Dmel_CG8066 | 345.072 | -0.741 | 0.236 | -3.134 | 1.72E-03 | 4.81E-02 |
| Dmel_CG1448 | 1370.609 | -0.865 | 0.277 | -3.128 | 1.76E-03 | 4.89E-02 |
| Dmel_CG1745 | 4690.633 | 0.672 | 0.215 | 3.128 | 1.76E-03 | 4.89E-02 |
| Dmel_CG7999 | 1414.777 | 0.475 | 0.152 | 3.126 | 1.77E-03 | 4.89E-02 |
| Dmel_CG8114 | 6061.248 | 0.364 | 0.116 | 3.126 | 1.78E-03 | 4.89E-02 |
| Dmel_CG12763 | 27.517 | -3.900 | 1.248 | -3.125 | 1.78E-03 | 4.89E-02 |
| Dmel_CG5083 | 2062.330 | 0.757 | 0.242 | 3.123 | 1.79E-03 | 4.92E-02 |
| Dmel_CG1487 | 3370.356 | 0.413 | 0.132 | 3.121 | 1.80E-03 | 4.93E-02 |

| | | | | | | |
|---|---|---|---|---|---|---|
| Dmel_CG5907 | 136.047 | -1.175 | 0.377 | -3.121 | 1.80E-03 | 4.93E-02 |
| Dmel_CG14162 | 1349.763 | 0.684 | 0.219 | 3.120 | 1.81E-03 | 4.93E-02 |
| Dmel_CG30118 | 7132.753 | 0.703 | 0.226 | 3.118 | 1.82E-03 | 4.96E-02 |
| Dmel_CG9078 | 3970.323 | 0.435 | 0.140 | 3.115 | 1.84E-03 | 4.98E-02 |
| Dmel_CG2249 | 2474.631 | -0.901 | 0.289 | -3.114 | 1.84E-03 | 4.98E-02 |
| Dmel_CG9220 | 125.885 | -0.824 | 0.265 | -3.113 | 1.85E-03 | 4.98E-02 |
| Dmel_CG8548 | 6621.909 | 0.439 | 0.141 | 3.113 | 1.85E-03 | 4.98E-02 |
| Dmel_CG8338 | 594.660 | -1.008 | 0.324 | -3.112 | 1.86E-03 | 4.98E-02 |
| Dmel_CG4407 | 1040.887 | 0.405 | 0.130 | 3.111 | 1.86E-03 | 4.98E-02 |
| Dmel_CG3671 | 2152.230 | 0.679 | 0.218 | 3.111 | 1.86E-03 | 4.98E-02 |
| Dmel_CR46037 | 8285.229 | -1.223 | 0.393 | -3.111 | 1.87E-03 | 4.98E-02 |
| Dmel_CG9342 | 904.312 | 0.552 | 0.178 | 3.106 | 1.90E-03 | 5.03E-02 |
| Dmel_CG33722 | 1440.994 | 0.563 | 0.181 | 3.106 | 1.90E-03 | 5.03E-02 |
| Dmel_CG12242 | 24.371 | -1.777 | 0.572 | -3.105 | 1.90E-03 | 5.03E-02 |
| Dmel_CG34333 | 20400.941 | 1.614 | 0.520 | 3.104 | 1.91E-03 | 5.04E-02 |
| Dmel_CG15431 | 62.781 | -0.872 | 0.281 | -3.104 | 1.91E-03 | 5.04E-02 |
| Dmel_CG32850 | 1595.721 | -0.790 | 0.255 | -3.100 | 1.93E-03 | 5.08E-02 |
| Dmel_CG1311 | 1395.571 | 0.650 | 0.210 | 3.097 | 1.96E-03 | 5.13E-02 |
| Dmel_CG12404 | 1283.354 | 0.419 | 0.136 | 3.095 | 1.97E-03 | 5.14E-02 |
| Dmel_CR45146 | 7.571 | -1.851 | 0.598 | -3.094 | 1.98E-03 | 5.15E-02 |
| Dmel_CG2934 | 5007.021 | 0.586 | 0.190 | 3.092 | 1.99E-03 | 5.17E-02 |
| Dmel_CG14621 | 2331.371 | 0.753 | 0.244 | 3.087 | 2.02E-03 | 5.24E-02 |
| Dmel_CG15094 | 334.092 | 0.800 | 0.259 | 3.084 | 2.04E-03 | 5.30E-02 |
| Dmel_CG11263 | 812.988 | -0.804 | 0.261 | -3.078 | 2.09E-03 | 5.38E-02 |
| Dmel_CG5848 | 5306.631 | 0.850 | 0.276 | 3.077 | 2.09E-03 | 5.38E-02 |
| Dmel_CG3227 | 1353.712 | 1.111 | 0.361 | 3.077 | 2.09E-03 | 5.38E-02 |
| Dmel_CG1569 | 2450.503 | 0.625 | 0.203 | 3.075 | 2.10E-03 | 5.39E-02 |
| Dmel_CG3961 | 102.114 | -0.958 | 0.312 | -3.075 | 2.11E-03 | 5.39E-02 |
| Dmel_CG7692 | 1472.997 | 0.524 | 0.170 | 3.074 | 2.11E-03 | 5.39E-02 |
| Dmel_CG16840 | 80.024 | -1.618 | 0.526 | -3.073 | 2.12E-03 | 5.40E-02 |

| | | | | | | |
|---|---|---|---|---|---|---|
| Dmel_CG5857 | 1494.849 | 0.656 | 0.214 | 3.071 | 2.13E-03 | 5.42E-02 |
| Dmel_CG34163 | 207.187 | -1.039 | 0.338 | -3.069 | 2.15E-03 | 5.44E-02 |

**table S18**. *D. melanogaster* genes Wald Test significant results for ~Infection vs ~Genotype+Infection+Genotype*Infection

| gene_id | baseMean | log2FoldChange | lfcSE | stat | pvalue | padj |
|---|---|---|---|---|---|---|
| Dmel_CG32834 | 13.460 | -21.521 | 2.735 | -7.868 | 3.61E-15 | 3.64E-11 |
| Dmel_CG2187 | 500.405 | -2.857 | 0.369 | -7.751 | 9.12E-15 | 4.59E-11 |
| Dmel_CG3104 | 56.006 | 2.402 | 0.476 | 5.043 | 4.59E-07 | 1.54E-03 |
| Dmel_CG10045 | 10067.445 | -4.248 | 0.867 | -4.901 | 9.52E-07 | 2.40E-03 |
| Dmel_CG7404 | 1099.831 | -0.694 | 0.152 | -4.557 | 5.20E-06 | 1.05E-02 |
| Dmel_CG5937 | 33.106 | 2.974 | 0.677 | 4.394 | 1.11E-05 | 1.86E-02 |
| Dmel_CG9650 | 454.888 | 1.566 | 0.360 | 4.350 | 1.36E-05 | 1.96E-02 |
| Dmel_CG8805 | 2531.220 | -0.929 | 0.218 | -4.270 | 1.95E-05 | 2.46E-02 |
| Dmel_CG15599 | 55.457 | 3.314 | 0.789 | 4.200 | 2.67E-05 | 2.61E-02 |
| Dmel_CG9155 | 396.786 | 1.554 | 0.373 | 4.164 | 3.12E-05 | 2.61E-02 |
| Dmel_CG6178 | 2685.460 | -0.699 | 0.168 | -4.163 | 3.14E-05 | 2.61E-02 |
| Dmel_CG17167 | 307.196 | -1.705 | 0.411 | -4.151 | 3.31E-05 | 2.61E-02 |
| Dmel_CG13772 | 15.539 | 3.746 | 0.905 | 4.139 | 3.48E-05 | 2.61E-02 |
| Dmel_CG8388 | 689.979 | -0.713 | 0.173 | -4.121 | 3.78E-05 | 2.61E-02 |
| Dmel_CG3157 | 1113.612 | -1.129 | 0.275 | -4.114 | 3.89E-05 | 2.61E-02 |
| Dmel_CG43079 | 631.857 | 2.061 | 0.503 | 4.096 | 4.21E-05 | 2.65E-02 |
| Dmel_CG18870 | 2820.357 | -0.823 | 0.202 | -4.070 | 4.70E-05 | 2.78E-02 |
| Dmel_CG18522 | 301.389 | 2.247 | 0.555 | 4.048 | 5.16E-05 | 2.89E-02 |
| Dmel_CG12002 | 217.218 | 2.463 | 0.611 | 4.029 | 5.60E-05 | 2.93E-02 |
| Dmel_CG31901 | 28.374 | 3.961 | 0.985 | 4.020 | 5.82E-05 | 2.93E-02 |
| Dmel_CG10593 | 2009.122 | -2.174 | 0.544 | -3.993 | 6.52E-05 | 3.09E-02 |
| Dmel_CG10573 | 175.250 | 1.302 | 0.327 | 3.985 | 6.75E-05 | 3.09E-02 |
| Dmel_CR43461 | 15.553 | 2.695 | 0.680 | 3.965 | 7.34E-05 | 3.12E-02 |
| Dmel_CG16762 | 7.124 | 4.634 | 1.172 | 3.955 | 7.66E-05 | NA |
| Dmel_CG13871 | 15.439 | -5.015 | 1.269 | -3.952 | 7.75E-05 | 3.12E-02 |
| Dmel_CG8663 | 114.131 | 2.257 | 0.573 | 3.941 | 8.11E-05 | 3.12E-02 |
| Dmel_CG30046 | 65.427 | -1.745 | 0.444 | -3.929 | 8.54E-05 | 3.12E-02 |
| Dmel_CR44115 | 21.000 | 4.551 | 1.163 | 3.913 | 9.12E-05 | 3.12E-02 |

| | | | | | | |
|---|---|---|---|---|---|---|
| Dmel_CG31973 | 444.390 | 1.434 | 0.367 | 3.906 | 9.40E-05 | 3.12E-02 |
| Dmel_CG6542 | 4210.347 | -0.815 | 0.209 | -3.903 | 9.51E-05 | 3.12E-02 |
| Dmel_CG5870 | 1313.514 | 1.321 | 0.339 | 3.894 | 9.84E-05 | 3.12E-02 |
| Dmel_CG9610 | 35.323 | 2.525 | 0.648 | 3.893 | 9.90E-05 | 3.12E-02 |
| Dmel_CG10726 | 2404.031 | -1.042 | 0.268 | -3.886 | 1.02E-04 | 3.12E-02 |
| Dmel_CG15814 | 1716.320 | -0.877 | 0.227 | -3.871 | 1.09E-04 | 3.12E-02 |
| Dmel_CG8256 | 121.600 | 1.720 | 0.445 | 3.865 | 1.11E-04 | 3.12E-02 |
| Dmel_CG7660 | 20442.794 | -0.690 | 0.179 | -3.861 | 1.13E-04 | 3.12E-02 |
| Dmel_CG9073 | 36.796 | 2.621 | 0.679 | 3.858 | 1.14E-04 | 3.12E-02 |
| Dmel_CG32082 | 143.362 | 1.635 | 0.424 | 3.854 | 1.16E-04 | 3.12E-02 |
| Dmel_CG9888 | 413.992 | 1.902 | 0.494 | 3.851 | 1.18E-04 | 3.12E-02 |
| Dmel_CG9220 | 125.885 | 1.424 | 0.371 | 3.840 | 1.23E-04 | 3.18E-02 |
| Dmel_CG9772 | 2989.847 | -0.860 | 0.224 | -3.833 | 1.27E-04 | 3.19E-02 |
| Dmel_CG12283 | 681.554 | 1.430 | 0.375 | 3.819 | 1.34E-04 | 3.29E-02 |
| Dmel_CG5252 | 4020.919 | -0.666 | 0.175 | -3.808 | 1.40E-04 | 3.37E-02 |
| Dmel_CG12763 | 27.517 | 7.141 | 1.880 | 3.799 | 1.45E-04 | 3.41E-02 |
| Dmel_CR44472 | 23.272 | -2.646 | 0.702 | -3.772 | 1.62E-04 | 3.69E-02 |
| Dmel_CG43758 | 309.775 | 1.714 | 0.456 | 3.760 | 1.70E-04 | 3.69E-02 |
| Dmel_CG2275 | 1215.191 | -1.179 | 0.314 | -3.754 | 1.74E-04 | 3.69E-02 |
| Dmel_CG9559 | 4504.461 | -1.078 | 0.287 | -3.754 | 1.74E-04 | 3.69E-02 |
| Dmel_CG1977 | 2140.679 | 1.065 | 0.284 | 3.752 | 1.76E-04 | 3.69E-02 |
| Dmel_CG12375 | 831.121 | -0.777 | 0.208 | -3.739 | 1.85E-04 | 3.80E-02 |
| Dmel_CG40813 | 5.158 | 8.173 | 2.197 | 3.720 | 1.99E-04 | NA |
| Dmel_CG3812 | 1006.282 | -0.794 | 0.215 | -3.698 | 2.17E-04 | 4.33E-02 |
| Dmel_CR46350 | 65.211 | 1.927 | 0.521 | 3.696 | 2.19E-04 | 4.33E-02 |
| Dmel_CG3879 | 12.492 | 3.044 | 0.827 | 3.683 | 2.31E-04 | 4.43E-02 |
| Dmel_CR45530 | 16.389 | 2.296 | 0.624 | 3.680 | 2.33E-04 | 4.43E-02 |
| Dmel_CG12477 | 8.616 | 5.136 | 1.396 | 3.679 | 2.34E-04 | NA |
| Dmel_CG17292 | 1474.083 | -1.032 | 0.281 | -3.674 | 2.39E-04 | 4.45E-02 |
| Dmel_CG2807 | 1130.688 | 0.911 | 0.249 | 3.652 | 2.60E-04 | 4.76E-02 |
| Dmel_CG9379 | 238.648 | 1.353 | 0.371 | 3.643 | 2.69E-04 | 4.79E-02 |

| | | | | | | |
|---|---|---|---|---|---|---|
| Dmel_CG17927 | 1300.359 | 2.092 | 0.574 | 3.641 | 2.71E-04 | 4.79E-02 |
| Dmel_CG6006 | 523.400 | 1.551 | 0.427 | 3.636 | 2.77E-04 | 4.81E-02 |
| Dmel_CG3407 | 971.487 | -0.958 | 0.264 | -3.625 | 2.89E-04 | 4.82E-02 |
| Dmel_CG1462 | 323.655 | 1.723 | 0.476 | 3.621 | 2.94E-04 | 4.82E-02 |
| Dmel_CG13000 | 15.067 | 2.743 | 0.759 | 3.615 | 3.00E-04 | 4.82E-02 |
| Dmel_CG18375 | 181.968 | 1.050 | 0.291 | 3.614 | 3.01E-04 | 4.82E-02 |
| Dmel_CG8262 | 752.433 | -1.356 | 0.375 | -3.614 | 3.02E-04 | 4.82E-02 |
| Dmel_CG6227 | 371.074 | 1.175 | 0.326 | 3.604 | 3.13E-04 | 4.93E-02 |
| Dmel_CG18549 | 1584.276 | -0.675 | 0.188 | -3.597 | 3.22E-04 | 4.93E-02 |
| Dmel_CG5644 | 28.092 | -2.406 | 0.670 | -3.592 | 3.29E-04 | 4.93E-02 |
| Dmel_CG34323 | 38.530 | 2.239 | 0.624 | 3.587 | 3.34E-04 | 4.93E-02 |
| Dmel_CG17078 | 3064.101 | -1.210 | 0.338 | -3.584 | 3.39E-04 | 4.93E-02 |
| Dmel_CR46481 | 1090.588 | 1.301 | 0.363 | 3.581 | 3.43E-04 | 4.93E-02 |
| Dmel_CG3635 | 29.922 | 2.696 | 0.753 | 3.578 | 3.47E-04 | 4.93E-02 |
| Dmel_CG9901 | 6113.781 | -0.527 | 0.147 | -3.577 | 3.47E-04 | 4.93E-02 |
| Dmel_CG9707 | 1943.581 | -0.814 | 0.228 | -3.569 | 3.59E-04 | 5.02E-02 |
| Dmel_CG3171 | 1422.762 | -1.120 | 0.315 | -3.552 | 3.82E-04 | 5.27E-02 |
| Dmel_CG7002 | 154.856 | 2.295 | 0.648 | 3.539 | 4.02E-04 | 5.27E-02 |
| Dmel_CG4620 | 4335.134 | -0.936 | 0.265 | -3.538 | 4.04E-04 | 5.27E-02 |
| Dmel_CG6202 | 3774.453 | -0.627 | 0.177 | -3.537 | 4.04E-04 | 5.27E-02 |
| Dmel_CG8411 | 7127.715 | -0.661 | 0.187 | -3.536 | 4.06E-04 | 5.27E-02 |
| Dmel_CG11941 | 16.019 | 3.035 | 0.859 | 3.535 | 4.08E-04 | 5.27E-02 |
| Dmel_CG12110 | 4136.182 | -0.767 | 0.217 | -3.531 | 4.14E-04 | 5.28E-02 |
| Dmel_CG17839 | 101.907 | 1.620 | 0.459 | 3.526 | 4.22E-04 | 5.29E-02 |
| Dmel_CG1024 | 621.198 | -1.141 | 0.324 | -3.521 | 4.30E-04 | 5.29E-02 |
| Dmel_CG5445 | 1662.056 | -0.783 | 0.223 | -3.518 | 4.36E-04 | 5.29E-02 |
| Dmel_CG16778 | 54.657 | -3.076 | 0.875 | -3.517 | 4.36E-04 | 5.29E-02 |
| Dmel_CG5939 | 354.903 | 1.672 | 0.476 | 3.512 | 4.46E-04 | 5.34E-02 |

**table S19**. *D. melanogaster* genes with Wald Test significant results for ~Genotype*Infection vs ~Genotype+Infection+Genotype*Infection

| gene_id | baseMean | log2FoldChange | lfcSE | stat | pvalue | padj |
|---|---|---|---|---|---|---|
| WD_RS03770 | 149.412 | 0.486 | 0.160 | 3.043 | 2.34E-03 | 7.92E-01 |
| WD_RS05260 | 22.714 | -1.127 | 0.380 | -2.969 | 2.99E-03 | 7.92E-01 |
| WD_RS06475 | 10.609 | 1.217 | 0.449 | 2.710 | 6.72E-03 | 7.92E-01 |
| WD_RS05810 | 49.056 | -0.535 | 0.199 | -2.687 | 7.20E-03 | 7.92E-01 |
| WD_RS04825 | 14.115 | 1.302 | 0.486 | 2.678 | 7.41E-03 | 7.92E-01 |
| WD_RS04740 | 25.586 | -1.220 | 0.461 | -2.646 | 8.13E-03 | 7.92E-01 |
| WD_RS04370 | 80.236 | -0.387 | 0.148 | -2.622 | 8.73E-03 | 7.92E-01 |
| WD_RS01760 | 8.222 | 1.263 | 0.491 | 2.572 | 1.01E-02 | 7.92E-01 |
| WD_RS01300 | 60.484 | 0.563 | 0.223 | 2.519 | 1.18E-02 | 7.92E-01 |
| WD_RS05640 | 10.502 | -1.425 | 0.567 | -2.513 | 1.20E-02 | 7.92E-01 |
| WD_RS02205 | 25.001 | -1.221 | 0.496 | -2.464 | 1.37E-02 | 8.26E-01 |
| WD_RS01390 | 25.646 | 0.699 | 0.303 | 2.306 | 2.11E-02 | 9.96E-01 |
| WD_RS03775 | 1404.573 | 0.354 | 0.161 | 2.207 | 2.73E-02 | 9.96E-01 |
| WD_RS00480 | 8.281 | 1.001 | 0.454 | 2.204 | 2.76E-02 | 9.96E-01 |
| WD_RS05480 | 70.888 | -0.398 | 0.181 | -2.203 | 2.76E-02 | 9.96E-01 |
| WD_RS01110 | 12.261 | -0.982 | 0.447 | -2.198 | 2.80E-02 | 9.96E-01 |
| WD_RS03920 | 32.007 | -0.685 | 0.313 | -2.190 | 2.85E-02 | 9.96E-01 |
| WD_RS05665 | 11.884 | 0.905 | 0.422 | 2.145 | 3.19E-02 | 9.96E-01 |
| WD_RS05520 | 17.489 | 0.821 | 0.388 | 2.118 | 3.42E-02 | 9.96E-01 |
| WD_RS01790 | 16.771 | -1.521 | 0.738 | -2.062 | 3.92E-02 | 9.96E-01 |
| WD_RS01335 | 36.739 | -0.448 | 0.219 | -2.050 | 4.03E-02 | 9.96E-01 |
| WD_RS00655 | 32.020 | -0.467 | 0.228 | -2.046 | 4.07E-02 | 9.96E-01 |
| WD_RS04175 | 9.229 | 0.982 | 0.483 | 2.034 | 4.20E-02 | 9.96E-01 |
| WD_RS01525 | 14.722 | 0.836 | 0.417 | 2.002 | 4.52E-02 | 9.96E-01 |

| | | | | | | |
|---|---|---|---|---|---|---|
| WD_RS01990 | 15.357 | 0.803 | 0.405 | 1.985 | 4.72E-02 | 9.96E-01 |
| WD_RS02935 | 45.227 | 0.477 | 0.240 | 1.985 | 4.72E-02 | 9.96E-01 |
| WD_RS02225 | 22.418 | 0.650 | 0.333 | 1.949 | 5.13E-02 | 9.96E-01 |
| WD_RS04020 | 7.360 | 0.942 | 0.485 | 1.940 | 5.23E-02 | 9.96E-01 |
| WD_RS03070 | 8.315 | 1.286 | 0.665 | 1.934 | 5.32E-02 | 9.96E-01 |

**table S20**. wMel Wolbachia genes Wald Test significant results for ~Genotype vs ~1
